# Unveiling the complexity of post-Roman polity formation using ancient DNA

**DOI:** 10.1101/2025.08.18.670760

**Authors:** Yijie Tian, István Koncz, Norbert Faragó, Corina Knipper, Ronny Friedrich, Deven N. Vyas, Levente Samu, Olga Spekker, Tamás Szeniczey, Tamás Hajdu, Balázs Gusztáv Mende, Péter Tomka, Ildikó Katalin Pap, Dávid Czigány, Rita Radzeviciute, Luca Traverso, Guido Alberto Gnecchi-Ruscone, Paolo Francalacci, Bernd Schöne, Gábor Tóth, Anna Szécsényi-Nagy, Petrus le Roux, Kurt W. Alt, Zuzana Hofmanová, Walter Pohl, Johannes Krause, Tivadar Vida, Patrick J. Geary, Krishna R. Veeramah

## Abstract

The transformation of the Roman world (4th-9th centuries CE) culminating in the dissolution of the Western Roman Empire in the 5th century marked a fundamental transition in European history. "Barbarian" groups and Roman provincial societies merged into enduring political frameworks in some places but not in others. Key questions persist regarding the regionally specific nature of this transformation. Here, we present one of the most densely sampled early medieval datasets to date, enabling the reconstruction of a post-Roman polity at an unprecedented micro-regional resolution. Our integrated genetic and archaeological analysis of two Roman-period (n=68) and five post-Roman (n=246) sites from the Little Hungarian Plain, a former Roman frontier region, reveals a significant increase in Northern European genetic ancestry, likely reflecting a large-scale population movement into the region. Moreover, while all post-Roman sites, dated to the time of the Langobard Kingdom, share similar genetic profiles, burial practices, and material culture, they display distinct patterns of social organization, especially regarding the role that biological relatedness played in each community. Our results suggest that distinct social hierarchies might have existed between sites, and their intensive connectivity probably formed the basis for regional-level organisation within the post-Roman polity. Our study provides a framework for investigating how new polities take shape after the collapse of major political systems. Applied more broadly, this approach can deepen our understanding of societal transformation across early medieval Europe and beyond, offering a richer and more nuanced view of the human past than ever before.

## Introduction

The 4th-6th centuries CE marked a period of substantial demographic, cultural, and political change on the European continent, coinciding with the dissolution of the Western Roman Empire and the emergence of various post-Roman polities within and beyond its former territories^1–3^. While contemporary authors ascribed agency in these processes to “barbarian” peoples or gentes and highlighted the role of migration in these changes, in the past decades some historians have ascribed economic, social and cultural transformations to internal factors rather than to population movements.

The Little Hungarian Plain (henceforth LHP), located between the Alps and the Transdanubian Mountains and both sides of the Danube, provides an ideal case study for understanding these transformations. By the 4^th^ century CE it had become an important border area of the Roman Empire, with multiple major urban centers such as Savaria, Scarbantia and Arrabona, and well-developed military and civic infrastructure. Main trade routes crossed the region along the Danube to the west and the Amber Road running north to south^4–8^. Following the abandonment of the region by the Roman administration in the first half of the 5th century, the territory witnessed multiple political shifts, with the short-lived rule of the Huns and the appearance of groups of Germanic origin, including the Langobards who occupied this area around 500 CE and established a kingdom that lasted until their migration to Italy in 568^6,9–11^. While there is no clear evidence for the continuous habitation of Roman towns^7,8^, the emergence of new communities and the high diversity of archaeological sites show the importance of the region during the 6th century, as it became one of the core territories of this emerging new polity^10–13^.

Archaeological records from the region show changes in settlement structure and material culture suggesting a significant cultural transformation that affected everyday life, burial customs, sustenance strategies and political administration^4,10,11^. During the Roman period, communities formed a dense network via their social, administrative and economic roles that provided the basis of both local and regional organization^7,8,14,15^. The organization of post-Roman, early medieval polities however is more obscure. While 6th-century cemeteries display a picture of stratified communities and diverse long-distance cultural connections^10–12^, the lack of contemporaneous settlements and the silence of detailed written sources on the region indicate that these cemeteries represent small rural communities, providing little understanding of how they formed parts of larger structures^10,11^.

To better understand the development of the LHP during the period described above, we sequenced 314 ancient genomes dated between the 3rd and 6th centuries CE from seven separated sites (Fig 1). By integrating this data with isotopic and archaeological evidence, we aimed to address two key longstanding questions: (1) To what extent do changes described generally in historical sources and observed in the LHP archaeological record reflect underlying large-scale population movements rather than internal cultural and political transformations? (2) What forms of community formation, development and interactions could be observed among contemporaneous sites in a single region? Our results demonstrate a major increase of contemporary Northern European ancestry across the entire region between the 4th and 6th centuries CE, likely reflecting a process connected to the expansion of the Langobard settlement area. We also show that despite very similar genomic ancestry profiles at a regional level, local communities exhibited distinct social hierarchies that were interconnected on cultural, political, and biological levels, forming the foundation of the emerging post-Roman Langobard polity.

**Fig. 1.**
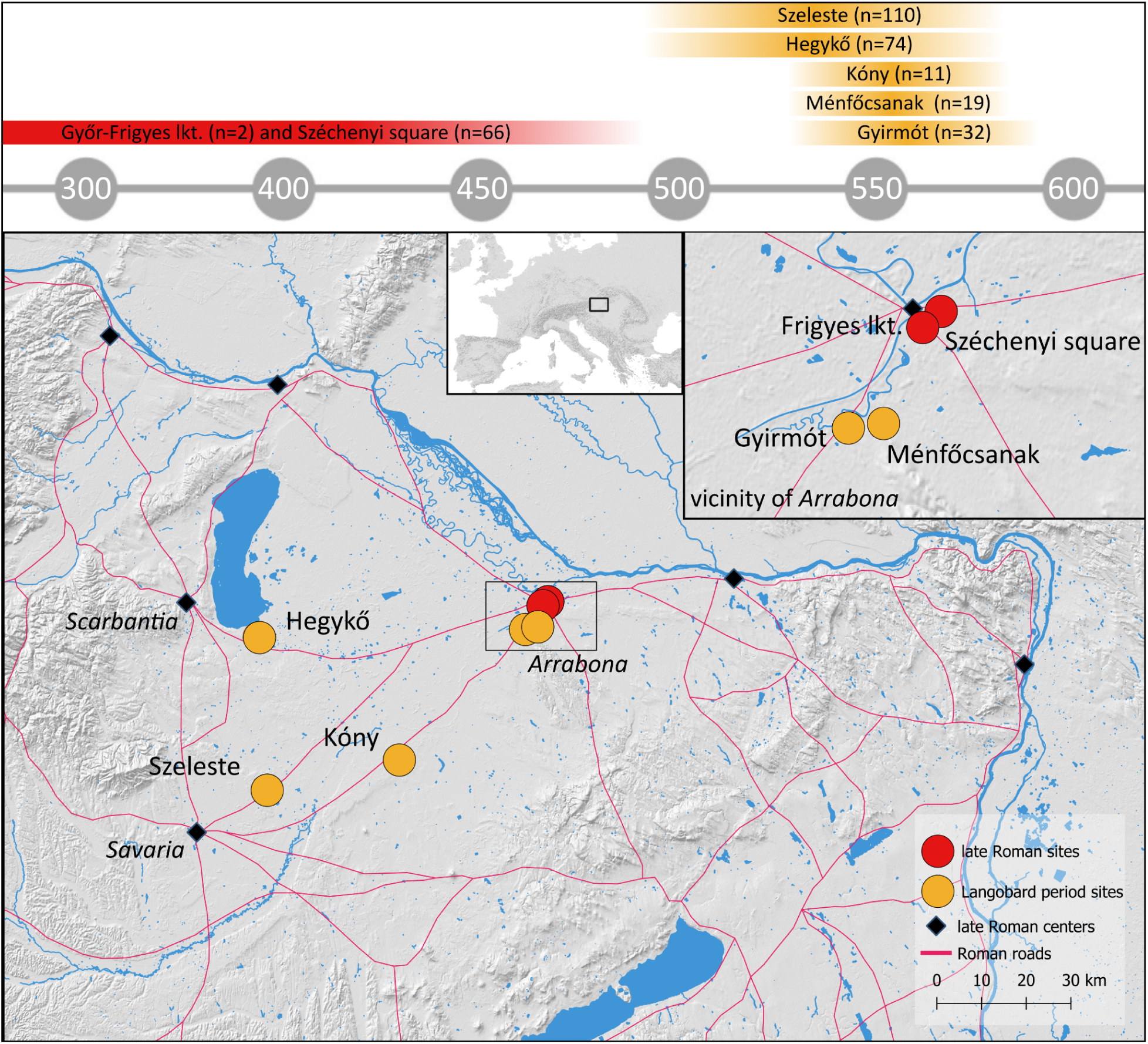
Geographic location and chronology of the sites included in this study. The sites are marked by red and orange circles, with their relative and absolute chronological positions (CE). The number of genetic samples is shown in brackets.

### Dense Sampling Of The LHP

We chose cemeteries with available archaeological and osteological material and conducted a comprehensive sampling strategy, aiming to include all the available individuals from all seven sites (Fig 1, Supplementary Table 1, Supplementary Section S1). The Late Roman period is represented by two sites near the Roman town of Arrabona (modern day Győr, Hungary): Győr-Frigyes lkt. is a small burial group (n=2) dated to the 3rd century CE, while Győr-Széchenyi square belonged to a community (n=66) dated to 4th-5th century CE, the latest phase of the Roman town ^16,17^. Burial sites are rare in the middle/second half of the 5^th^ century—the Hunnic period—when late Roman towns had lost their urban character and role in the region, but a series of newly founded cemeteries emerged around 500 CE^4,11^. During the Langobard occupation in the 6^th^ century, sites in the LHP ranged from large (ca. 100 burials) and small (ca. 20-40 burials) cemeteries to burial groups with only a few graves^10,12^. To represent this heterogeneity, we sampled two large sites, Hegykő (n=74) and Szeleste (n=110), founded at the end of the 5th century and continuously used until the second half of the 6th century^18–20^, and three smaller sites, Gyirmót (n=32), Kóny (n=11) and Ménfőcsanak (n=19), each in use only for a few decades in the middle of the 6th century^6,21^. Archaeological dating of the sites is supported by 38 newly produced radiocarbon dates from Győr-Széchenyi square and the 6th-century sites (Supplementary Table 1). All post-Roman sites are directly connected to late Roman infrastructure and/or earlier urban centers but represent rural communities living outside of the former towns^4,14,15^. They show similarities in their archaeological material, with shared artefact types (most notably brooches) and long-distance connections towards Northern and Western Europe and the Mediterranean^11,12,22–24^.

The comprehensive sampling strategy yielded 314 new ancient DNA (aDNA) genomes from the Late Roman period (n=68, two sites) and the 6th century (n=246, five sites), along with additional ^87^Sr/^86^Sr (n=247, 5 sites), δ^13^C and δ^15^N (n=283, 5 sites) measurements and radiocarbon dates (n=38, 6 sites) (Supplementary Table 1, Supplementary Section S2). Genomic libraries were prepared from petrous bone and underwent partial UDG treatment^25^ followed by an in-solution capture targeting ∼1.2 M single nucleotide polymorphisms (SNPs)^26,27^ (henceforth 1,240 K capture) and Illumina sequencing. The average coverage at these SNPs was ∼1.95×. Analysis of X chromosome-mapped reads in males and mtDNA in both sexes revealed low levels of estimated contamination in all individuals (mean ∼1%).

### Post-Roman Influx Of Northern Ancestry

Historical sources document multiple migratory events in the LHP between the middle of the 5th and 6th centuries that also correspond to significant changes in the archaeological record^4^. The relative lack of later 5th century sites and the absence of evidence for the continuous use of late Roman towns have been interpreted as signs of depopulation, if not total abandonment, of the area following the Roman period^4,7,8^. The appearance of new communities around 500 CE and during the 6th century are generally attributed to the expansion of the Langobards into the region^6,10,11^

To examine evidence of migration as reflected by changes in genomic ancestry, we first conducted principal component analysis (PCA), with the 3rd-5th (n=68) and 6th-century (n=246) individuals projected onto the modern European POPRES reference set^28^. We observed a change in the composition of population genetic ancestry between the two periods: 3-5th century individuals displayed a unimodal distribution on the PC1 axis (Silverman test, k=1: p=0.89), clustering primarily with modern southern European populations. In contrast, the 6th century showed a bimodal pattern (Silverman test, k=1: p=0; k=2: p=0.53) centered around both modern northern and southern European populations, the latter overlapping with the previous period. We also observed the 3rd-5th century samples had a greater variance in PC2 axis (t_test on variance, p<0.001) (Fig. 2a, see Extended Data Fig. 1. for a sample size controlled PCA). Consistent with these differences, the Roman and post-Roman period populations were considered significantly different as assessed by their *F_ST_* (*F_ST_*=0.0021, z-score=9.99).

**Fig. 2.**
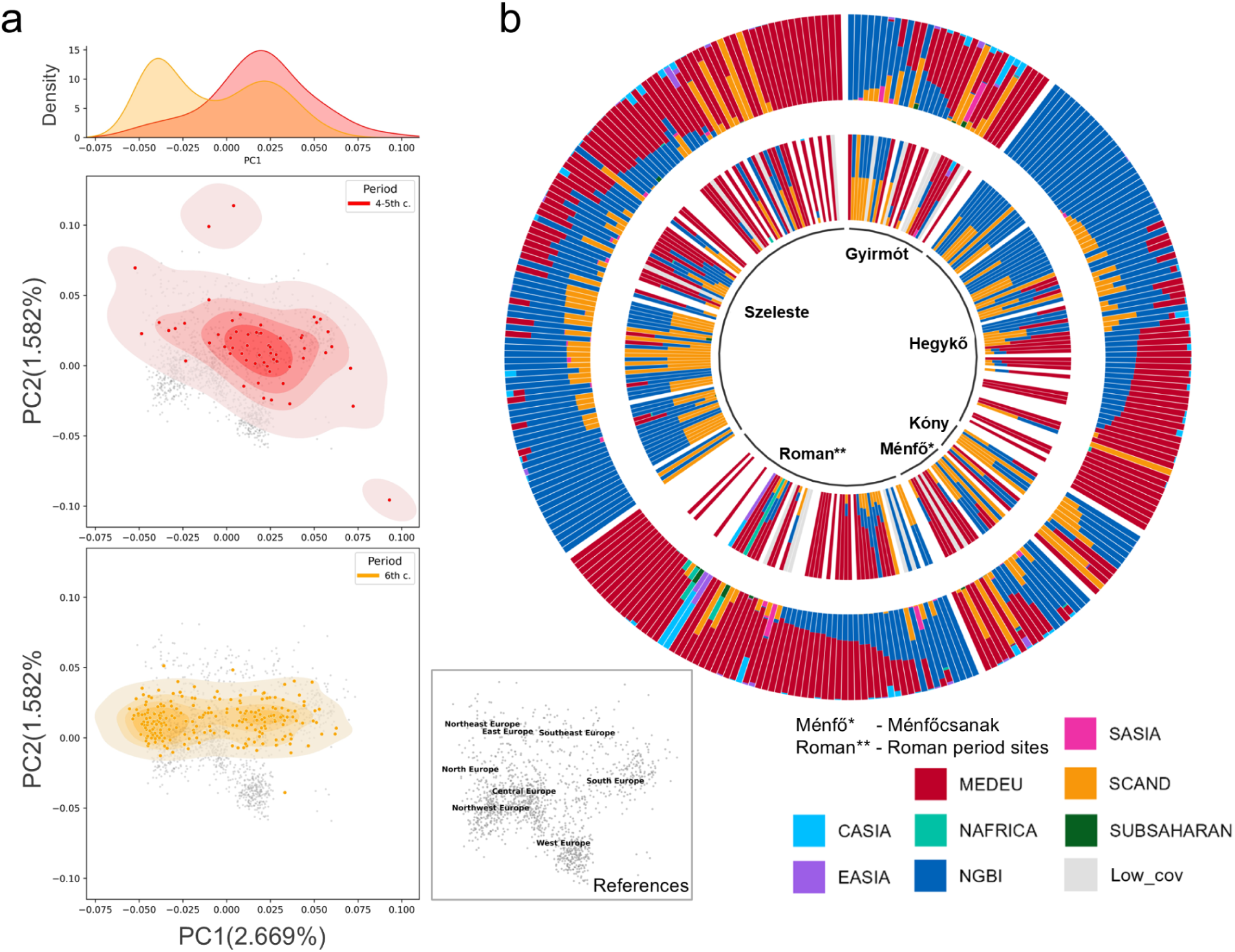
The genetic profiles of ancient individuals in this study. a. PCA of 3rd-5th century and 6th century populations in the Little Hungarian Plain. Black dots in the background represent reference data from POPRES. The Kernel Density Estimation of PC1 is displayed on the top. b. FastNGSadmix results are shown in the outer circle, while qpAdm results are displayed in the inner circle. FastNGSadmix includes all individuals in its plot, whereas qpAdm only displays individuals that can be successfully modeled (p>0.01). Reference (source) groups for both analyses are shown as legend.

We also applied two model-based analyses (fastNGSadmix^29^ and qpAdm^30^) (Fig. 2b) in order to estimate individual genomic ancestry proportions based on a penecontemporary reference panel (modified from Vyas and Koncz et al.^31^, Supplementary Table 2) that consisted of n=219 4-8th CE individuals grouped into eight ancient populations according to their geographic origins. The fastNGSadmix results (Supplementary Table 3 and Supplementary Section S3) were consistent with the PCA: 3rd-5th century sites exhibited a major Southern European (MEDEU) ancestry (64.5%), while 6th-century sites displayed a higher northern ancestry compared with Southern European ancestry (59.3% and 38.6%, respectively). Despite using modern reference populations, the PC1 values were significantly correlated with both ancient Northern and Southern European ancestry (Pearson’s *r* = –0.94 and 0.94, respectively; both p<0.001). We also observed a higher proportion of non-European ancestry (CASIA, EASIA, NAFRICA, SASIA, and SUBSAHARAN) in the 3-5th century sites compared to the 6th century sites (mean=5.9% vs. 1.3%, t-test *p*=0.03). When using qpAdm (Supplementary Table 4) 77% of LHP individuals could be modeled with one, two, or three-source models (*p*>0.01), and yielded results that closely aligned with those from fastNGSadmix (the estimated proportions of all eight ancestries, except for SUBSAHARAN and SASIA, were significantly correlated (*p*<0.001)) (Supplementary section S4).

We then applied qpWave^30^ to test whether the Roman and post-Roman periods can be modeled using the same or distinct ancestral sources (Methods). To achieve this, we first compiled published genomic data from ancient European samples with median ages that predated or were contemporaneous with our two Roman-period LHP sites^32–40^ and grouped them according to their sampling locations (Extended Data Fig. 2), forming nine distinct source populations (Supplementary Table 2). For individuals from the 3rd-5th century, 43 out of 68 could be modeled using one of the penecontemporaneous sources (Fig 3a, Supplementary Table 5), among which only 12% (n=5) could be attributed to Northern European populations (either Northeast_Europe_ia, Northern_Germany_4-8c, or Scandinavia_ia).

**Fig. 3.**
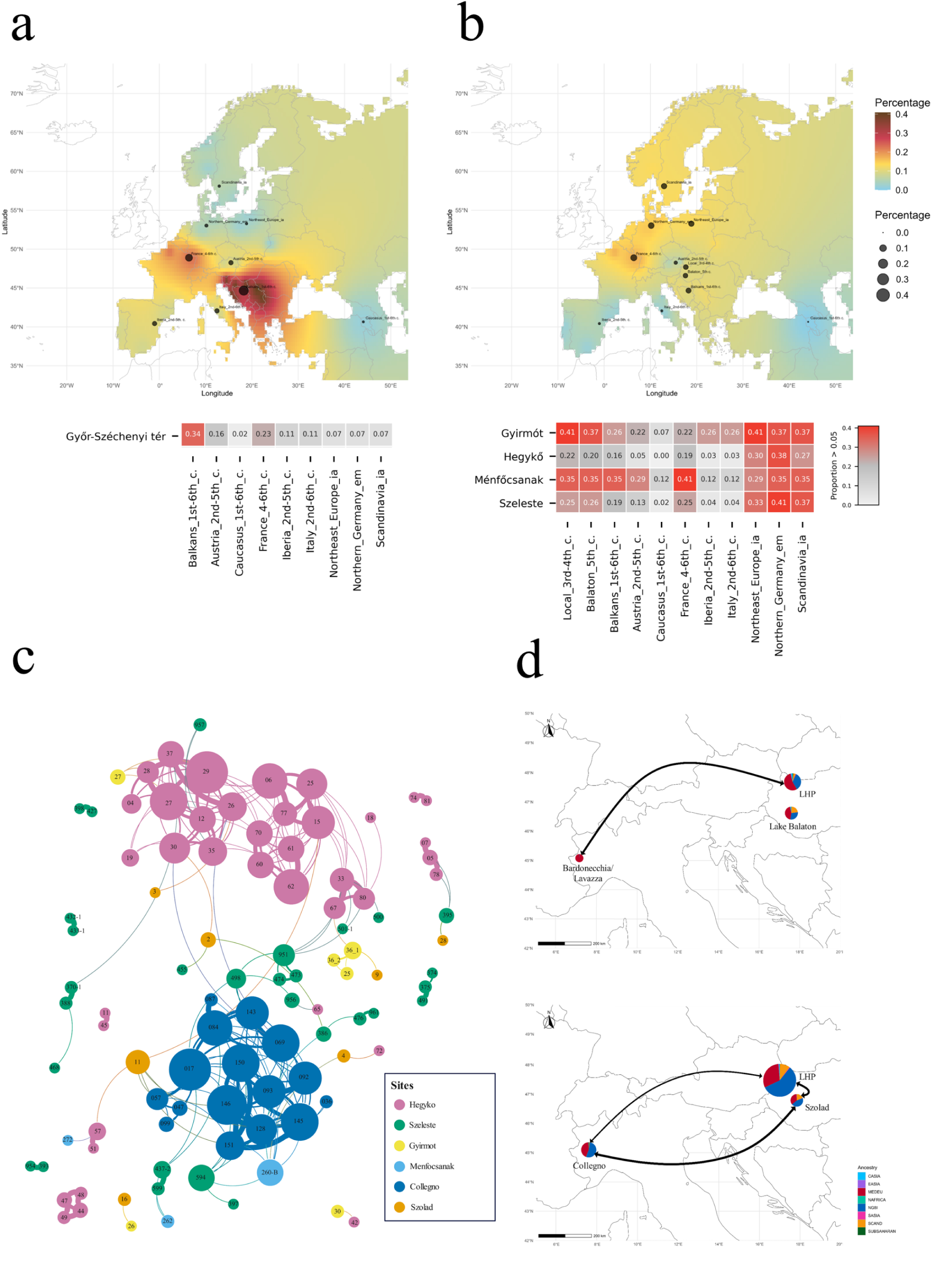
Shifts in genetic ancestry and connection patterns between populations from the Roman and post-Roman periods. a-b, QpWave modeling results for (a) Roman period sites and (b) post-Roman sites. The upper map displays the interpolation results, while the lower one shows the heatmap results. (c) IBD connections (>=12cM) among post-Roman period sites, including published data from Collegno and Szólád. Each node represents an individual, with the size proportional to the number of connections that individual has. Each line (edge) indicates haplotype sharing, and its width corresponds to the normalized pi-hat values (0-0.5) (d) Changes in connections between pre-Lombard period sites (top) and post-Lombard period sites (bottom). The width of the lines reflects the normalized IBD connection strength between each site.

When examining individuals from the 6th century, we incorporated two additional source populations: the 3-5th century populations from the LHP (Local 3rd-4th c.) and the previously published 5th-century Lake Balaton populations (Balaton_5th c.) from the wider region (Supplementary Table 1-2, Fig. 3b)^31^. In stark contrast to the previous period, of the 213 of 246 (87%) 6th century individuals that can be modeled with one source (Supplementary Table 6), 41% (n=87) were attributed to Northern European populations. This notable increase (29%) suggests a possible influx of individuals from the more northern parts of Europe during the 6th century. However, we observed that 20% of individuals could be modeled by “local” groups (Local 3rd-4th c. and Balaton_5th c.) pointing to some autochthonous development of the genetic profile in the post-Roman period.

Similar to the change observed in the genomic data, δ^13^C and δ^15^N data (283 new measurements and 11 earlier published measurements, Supplementary Section S2) show that adult individuals significantly differ between the Roman and post-Roman period (Mann-Whitney test for stochastic equality - δ^13^C: z=5,462; *p*<0.0001; δ^15^N: z=8,292; *p*<0.0001), with higher δ^15^N and δ^13^C values generally attributed to the 4-5th-century individuals. These differences remained significant even after controlling for genetic ancestry using PC1 and PC2 as covariates in a PERMANOVA test (based on a δ¹³C/δ¹⁵N distance matrix, with PC1, PC2, and Period as predictors; *p*=0.001 for Period). These results suggest a change in subsistence strategies and nutrition between the two periods, though this may be linked to the different nature of the sites (for example urban vs rural contexts) rather than just the direct result of a newly arriving population with their own distinct diet. However, variation was also observed between the 6th-century sites (Fig. 4). Moreover, multiple statistically significant correlations were observed between genetic ancestry and either δ^15^N or δ^13^C values (Extended Data Fig. 3): individuals with more Southern European ancestry tended to have higher δ^13^C values in Hegykő (p=0.002) and Szeleste (p=0.009), while at Szeleste individuals with higher Northern European ancestry also demonstrated higher δ^15^N values (p=0.003) suggesting that parts of the population in the 6th century communities might have retained a diet similar to the earlier period.

**Fig. 4.**
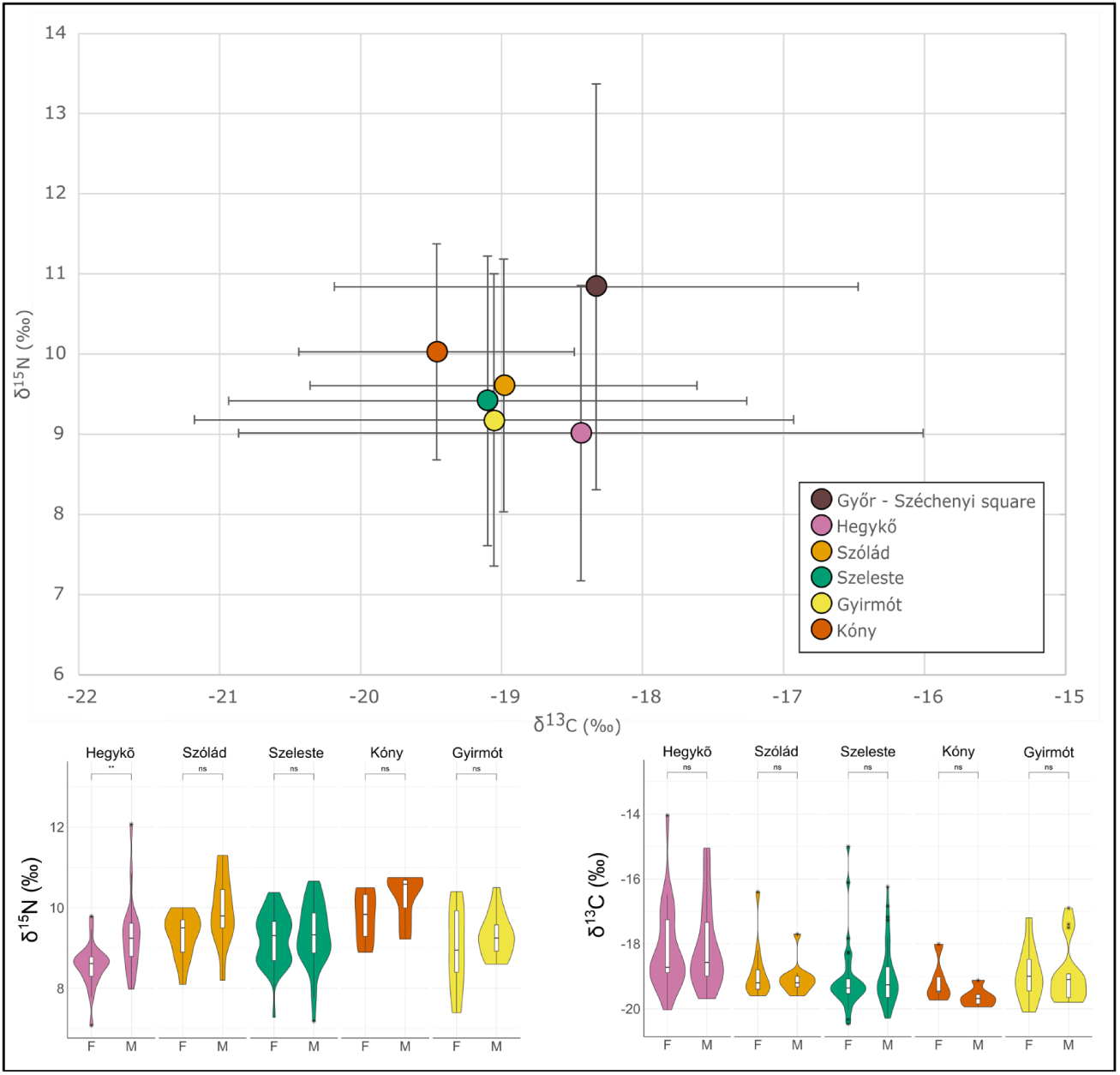
δ¹³C and δ¹⁵N values from the sites in this study. The top panel shows a scatter plot of δ¹³C and δ¹⁵N for all sites, with means and ±2 standard deviations indicated by whiskers. The bottom two panels present the same dataset broken down into δ¹³C and δ¹⁵N values by sex for the 6^th^ century CE site, using combined violin and box plots. Whiskers extend from the box to the largest and smallest data points within 1.5 times the interquartile range; values beyond this range are shown as outliers, marked with circles and stars.

To explore patterns of individual-level mobility across the region in the post-Roman period, we also conducted ^87^Sr/^86^Sr analysis on all available individuals from the 6th-century sites (n=201, Supplementary Table 1, Supplementary Section S2). We defined possible ‘local’ ranges based on environmental samples, measurements from bone and values of non-adult individuals site-by-site. The Sr isotope values showed that the ratio of the individuals outside of ranges identified as possibly ‘local’ varies site by site, between 43 to 74%. These are mostly adult males and females (above 90%) with the exception of Hegykő where it includes 35.3% non-adult individuals. Adult males and females possessed similar ranges across sites with the exception of Gyirmót, while adults and nonadults demonstrated similar ranges with the exception of Hegykő. Interestingly, local versus non-local Sr isotope values showed no correlation with genetic ancestry (p>0.05). Thus while there was a broad influx of a population with predominantly Northern European genomic ancestry into the region, individual-level mobility was apparently not restricted by this population-level process, and individuals with southern ancestry that more likely represented the continuity of Roman period local population were also moving between regions and into these communities as they formed and developed.

Finally, we note that HapROH^41^ identified only 17 out of 314 individuals with at least one Run of Homozygosity segment longer than 5 cM (max - 53cM, mean = 12cM), and of those 17, none were not enriched at any particular site (Supplementary Table 1), pointing to no major evidence of inbreeding within our communities, either during the Roman or post-Roman periods.

### Genetics Supports Langobard Migration

The significant post-Roman increase in northern genomic ancestry and shift in diet in the LHP indicate the possibility of a migratory event and that could be reflected in an enrichment of recent genealogical links between 6th-century individuals in the LHP and specific external regions or sites. Therefore we applied ancIBD^42^ to detect Identity By Descent (IBD) segments indicative of shared recent ancestry up to the 6th degree between individuals in the LHP and 861 penecontemporaneous individuals (Supplementary Table 2) across Eurasia and Africa (Methods).

Overall, given the large number of pairwise comparisons (n=111,930), we found only a very small number (244, ∼0.22%) of IBD connections greater than 12 cMs between post-Roman individuals from the LHP and other contemporaneous populations. A similar proportion was found when examining the Roman period individuals (n=26,691 pairwise comparisons, 41 connections or 0.15%) (Supplementary Table 7, Supplementary Section S5). Of the small number of connections that were observed with post-Roman individuals, those with Northern European individuals predominate (181/244, 74%). However, a) these individuals are also dominant in the reference set (303/861, 35%), b) no particular northern region was enriched, with connections observed with eight different modern northern European countries and c) the highest number of external IBD connections observed for the Roman period also involved northern European individuals (11/24, 45%). This pattern alone would not point to a specific 6th-century migratory event or pulse associated with the Longobards into the LHP from a particular area in Europe, as suggested by McColl et. al^43^, but rather a more general regional southward expansion into the territory of the (former) Western Roman Empire that was possibly accelerated after its fall. However, more targeted sampling of specific Roman-era sites north of the LHP in the future may yet unveil such an event. In addition, the increased IBD connections with northern European populations may simply reflect the smaller effective population size associated with this genomic ancestry.

In comparison, when focusing on sites along the historically documented route of the Langobard migration into northern Italy in the 6th century, we observed markedly different patterns before and after the presumed migration. For sites pre-dating the attested migration in 568—Győr-Széchenyi tér (4-5th century) in the LHP, 5th century sites in Lake Balaton^31^ ,and 5-7th century sites in Bardonecchia and Torino Lavazza in northern Italy^31^ — we only identified two IBD connections (Fig. 3d_top). However, when focusing on the post-Roman sites along the same route—the 6th century LHP sites, Szólád and the 6-8th century site of Collegno—we observed multiple connections that significantly shaped the network constructed from IBD connections between sites from the LHP, Lake Balaton and Northern Italy (Fig. 3c, 3d_bottom). When we incorporated the fastNGSadmix results into the nodes of the IBD network, a clear pattern emerged: the network was primarily structured around individuals with predominantly Northern European ancestry (Extended Data Fig. 4). We then applied FLARE^44^ using reference populations from the 1000 Genomes Project to infer the local ancestry for all the individuals in the network (Methods, Supplementary Section S6) and found that the distribution of IBD segments associated with northern ancestry was significantly larger than that based on southern ancestry (Wilcoxon rank-sum test, p<0.001)(Extended Data Fig 5). The findings suggest that the individuals from different sites were predominantly interconnected through shared northern haplotype segments.

Altogether, our results indicate a substantial increase in Northern European genetic ancestry between the 3rd–5th and 6th centuries in the LHP, most likely driven by the movement of groups from outside the former Empire into the region. The rarity of sites in the late 5th century indirectly suggests that this large-scale demographic shift likely occurred around 500 CE, coinciding with the emergence of newly established communities. However, the data do not support a model of complete population replacement. Instead, they point to the persistence and cohabitation of earlier population groups from the Roman period—who were also locally mobile—alongside incoming groups. These changes could be understood as the results of the expansion of the Langobard Kingdom into the region, described in some detail by the written sources^9,11,45,46^. This interpretation is further supported by IBD links between multiple regions that align with the historically inferred route of Langobard migration from Middle Danube region to Italy^45,46^.

### Diverse Post-Roman Community Formation

The Langobards caught the attention of contemporaneous historians at the end of the 5th century, when they appeared at the Lower Austrian Danube having already settled in Moravia (today’s Czechia). Soon after 500 CE, they established their kingdom in the Middle Danube region^11,46,47^. The LHP became an important core territory of the Langobard Kingdom, as attested by the emergence of a series of new communities throughout this period, placing them on the European political stage for the first time and becoming an important stepping stone before their migration and eventual conquest of Italy^11,45,46^. Archaeological research has described different types of cemeteries in Pannonia based on their chronology, size and cultural contacts^10–12^. Hegykő and Szeleste were founded around 500 CE, and the smaller sites, Gyirmót, Ménfőcsanak and Kóny, were founded in the middle third of the 6th century^6,11,12,21^. While all sites are similar in terms of burial customs, including grave design and funerary representation, this chronological difference is observable in their archaeological material, as certain phenomena that are generally found in 5th century contexts, such as artificial cranial deformation, brick-structure graves or polyhedral earrings with garnet inlays, are only present in the earlier founded sites^13,20,48^. Artifact types dated to the middle of the 6th century, such as S-brooches and rosette-shaped disc brooches show up in all 6th-century sites, while similarities in female dress and weaponry show cultural contacts and suggest that these communities were part of the same distribution networks^12,24^.

The largest pairwise *F_ST_* observed amongst the four 6th-century sites with n>=10 (after adjusting for related individuals) was between Hegykő and Ménfőcsanak (*F_ST_* =8.8×10^-4^), and none were significantly different using a z-score cutoff of 3. Based on the results from fastNGSadmix, qpAdm and PCA (Extended Data Fig. 6), all sites contained individuals with similar profiles of predominantly Northern and Southern European ancestry, resembling previously published data from nearby Szólád in Lake Balaton^49^.

We examined pairwise biological relatedness up to third-degree within the 6th century sites using lcMLkin^50^ and KIN^51^ (Methods, Supplementary Table 8-9, Extended Data Fig. 7, Supplementary Section S7). Hegykő demonstrated the highest ratio of relatedness with 67% of all individuals being part of the 10 inferred pedigrees. There were three pedigrees with more than four individuals, with the largest consisting of 13 individuals. In contrast, at Szeleste only 34% of the individuals belonged to the 14 pedigrees, only two of which contained more than 4 individuals (maximum 5). Gyirmót (n=32) only contained one third-degree related pair (6%), Kóny (n=11) two parent-offspring pairs (36%), and Ménfőcsanak (n=19) had no individuals related within third-degree. Differences in the patterns of more distant biological relatedness (up to 6th degree) were also evident from the ancIBD analysis for samples with 1240k coverage >=1x (n=130) (Fig. 3c). In Hegykő, multiple pedigrees—including all the larger ones—formed a tightly interconnected cluster. In contrast, relatedness at Szeleste exhibited a more fragmented structure with very few connections between pedigrees. The smaller sites showed no additional intra-site connections but did possess links to the larger sites in the LHP, as well as to Szólád and Collegno.

The role biological relatedness played in the formation of these neighboring communities appears to vary significantly, with three general patterns emerging (Fig 5). At Hegykő (as well as the previously published Szólád) pedigrees tended to only include members of homogeneous genetic ancestry, with the larger pedigrees dominated by Northern European genetic ancestry. Members of these genetically homogenous extended pedigrees were also buried together in the same area of the site (Extended Data Fig. 8), indicating that these biological connections could be understood as socially meaningful ties, driving a significant relationship between genetic ancestry and burial proximity (Mantel test, p<0.0001). Moreover, the exclusive presence of weapons, generally interpreted as signs of an elevated social and/or economic status in the period^11,46,52^, among members of these pedigrees suggests that they had an elevated social and/or economic status in the community, and due to the homogeneous genetic ancestry of these pedigrees and the exclusivity of weapons, there is also a significant statistical correlation between Northern European genetic ancestry and the presence of weapons as well (p<0.0001). While these communities were not strictly patrilocal, as multi-generational female lines also exist, these strong correlations between biological relatedness and grave goods are observed only in cases of male burials, suggesting a clear difference in burial representation between the two sexes. The elevated social status of the male members of the large pedigrees is also supported by observed differences in nutrition, as they showed higher δ^15^N values suggesting more access and intake of animal proteins (Fig. 4).

**Fig. 5.**
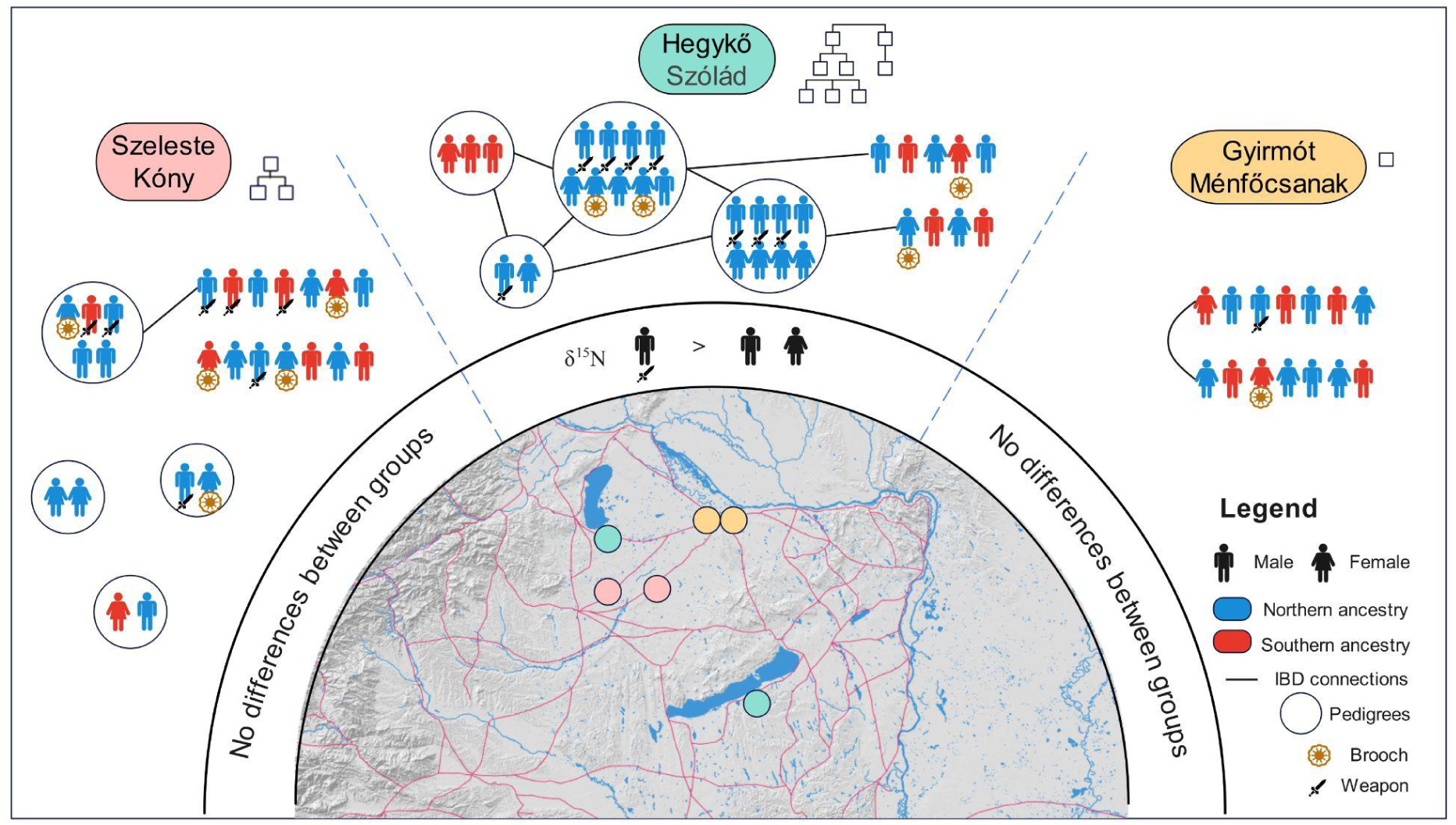
A schematic diagram illustrating the hierarchical organization of 6th-century communities. The outer layer reflects the extent to which each community was structured around biologically related groups. The middle layer indicates whether there are differences in isotope values between different groups. The inner layer displays the geographic distribution of the communities, with colors corresponding to the background colors of the site names.

In contrast, the smaller pedigrees found at Szeleste and Kóny are also more heterogeneous with regard to genomic ancestry. Only members of core families (i.e. parent-offspring sets) are buried together, while more distantly related individuals are buried separately in other parts of the cemeteries. Weapons are no longer restricted to members of pedigrees and also do not show significant correlation with any genetic ancestry. Moreover, no dietary differences are observed between various groups, which might indicate a less stratified community. Further, Ménfőcsanak and Gyirmót exhibit little or no biological relatedness (less than 10%) and again there is no correlation between genetic ancestry and spatial organization of the site or any aspect of burial customs. These sites also contain a lower ratio of weapons (10% and 13.3% among adult males respectively compared to 57.2% at Hegykő or 41.9% at Szeleste) as well as of female burials with gilded jewelry (0% and 16.7% compared to 28% or 21.7%), although burial disturbances could also have significant impact.

### A post-Roman polity in the making?

Contemporaneous written sources describe the gradual expansion of the Langobard settlement area in the Middle Danube region, but its extent, nature and political context have been much debated^10,11,46,47^. Viewed solely through the lens of population genetic structure, the five 6th-century LHP sites appear statistically indistinguishable. However, our comprehensive sampling strategy, integrating pairwise relatedness, burial practices, and historical-archaeological context, reveals that multiple modes of community formation were shaped by interactions between incoming groups enriched in Northern European ancestry and local populations with predominantly southern ancestry, but genetic ancestry does not necessarily correspond with social differences. Rather than existing as separate or parallel entities, these groups cohabited and mixed in varying ways across sites, generating diverse community structures that offer unprecedented insight into the formation and development of an emerging post-Roman polity.

Correlation between weapons, grave organization and extended pedigrees suggest that status inheritance was already established early in the 6th century in some communities, such as Hegykő, and that biological relatedness helped shape social kinship. Dominant lineages exhibit elevated social status that could indicate emerging elite communities of local or even regional importance. In contrast, at nearby Szeleste—archaeologically similar in burial style and material culture—such factors played a more limited role, highlighting the parallel emergence of distinct social models within the same region. Notably, these cemeteries were previously viewed as representing the same rural community type, underscoring the added resolution genomic data brings.

This mosaic of socially and culturally diverse communities likely formed the basis of a new regional post-Roman polity. Written sources describe its involvement in dynastic marital alliances as well as extensive wars with its neighbors, and by the middle of the 6th century it had become a defining regional power with the occupation of Southern Pannonia^46^. At the same time, settlement density increased, and new communities like those of Gyirmót, Kóny and Ménfőcsanak emerged, making great use of surviving late Roman infrastructure, especially roads^14,15^. In these later sites the dearth of within-site biological relatedness suggests top-down community organization, perhaps by political or military elites. This model is described by Paul the Deacon^53^ in the case of the conquest of Cividale in Italy, when the dux of the city handpicked his followers to settle and defend the city. The term *fara*, attested in several written sources in different contexts, may refer to either extended kin groups or mobile military units^54–56^. Our genomic results support the idea that the emerging Langobard society had begun to move beyond kinship-based structures toward more complex forms of social cohesion. Alternatively, the disrupted kinship structure in later cemeteries may reflect broader demographic shocks, such as the Justinianic Plague^57^.

Concurrently, new artefact types emerged which could be understood as local productions^24^. This process is especially visible in the case of S-shaped brooches, one of the most common and fast-changing jewelry types in all Western Europe during the period. While not completely replacing imported types, their widespread adoption suggests the presence of a strong communication network and shared cultural preferences^24^. These developments, together with the appearance of new communities and expansion of the settlement area, indicate the emergence of a shared identity and the consolidation of Langobard political power. Rather than a loose collection of rural settlements, our data suggest a hierarchically structured network of communities.

The complex biological and cultural interactions we observed reflect both the foundation and the consequence of maintaining social cohesion within this post-Roman polity. Our results demonstrate the value of integrating dense genomic data from multiple contemporaneous sites within a defined region with archaeological and historical evidence to uncover the dynamics of early medieval state formation.

## Data Availability

Our newly generated sequence data from 314 individuals are available from the NCBI Sequence Read Archive (SRA) database under accession PRJNA1285465. We accessed the POPRES (Population Reference Sample) dataset collected and published by Nelson et al.^28^ from dbGaP (accession phs000145.v4.p2).

## Acknowledgements

This project has received funding from the European Research Council (ERC) under the European Union’s Horizon 2020 Research and Innovation Programme (HistoGenes, grant 856453 ERC-2019-SyG); from the German Federal Ministry of Education and Research (BMBF) in the framework of Förderung durch das Bundesministerium für Bildung und Forschung (BMBF, grant number: 01 UA 0806B) and from Hungarian Scientific Research Fund (OTKA-NKFI; grant number: NN113157).

## Author contributions

Conceived and led by: I.K., P.J.G., K.R.V., Formal analyses: Y.T., I.K., N.F., D.N.V., P.F., P.l.R, Sample preparation and laboratory work: B.G.M, R.R., L.T., G.A.G-R., R.F., A., S-N.,C.K., K.W.A., B.S., D.C., T.S., T.H. P.T., I.K.P., P.l.R. Visualization:Y.T., I.K., N.F., S.L Writing: Y.T. I.K. N.F. Supervision: Z.H., W.P., J.K., T.V., P.J.G., K.R.V.

## Competing interests

The authors declare no competing interests.

## Methods

### Data Generation and Bioinformatic Processing

Various types of bones (Supplementary Table 1) were sampled from all individuals dated to the Late Roman period (n = 68, from two sites) and the 6th century (n = 246, from five sites). Sample preparation was carried out in the dedicated ancient DNA laboratory facilities at the HUN-REN RCH Institute of Archaeogenomics in Budapest, Hungary. Bone surfaces were decontaminated using UV-C light and mechanically cleaned. Approximately 25–50 mg of bone powder was obtained by drilling or grinding and subsequently transferred to the Max Planck Institute for Evolutionary Anthropology (MPI-EVA) in Leipzig, Germany. DNA extraction, single-stranded library preparation (ssLib), and two rounds of 1240k in-solution capture^27^ were performed at the MPI-EVA Core Unit. Sequencing was conducted in the same facility using the Illumina HiSeq 4000 platform, generating 75-cycle single-end reads. Each library was sequenced to a depth of approximately 20 million reads.

nf-core/eager v2.3.2 was employed for the following further processing of the raw reads ^58^: adapters were removed from the sequenced reads with the AdapterRemoval^59^; BWA v0.7.17 was employed to map the reads to the human reference genome with “bwa aln” (with the parameters: -n 0.01 -l 1024) and BWA samse (in case of single-end reads) or BWA sampe (paired-end)^60^; reads with <30 phred mapping quality were discarded with Samtools v1.9^61^. PCR Duplicate reads were removed with Picard tools MarkDuplicates v.2.26.0 (https://github.com/broadinstitute/picard).

To estimate the amount of C to T taphonomic deamination at the ends of the mapped fragments we used mapDamage v2.0^62^ run on a subset of 100,000 q30-reads.

Genotypes were called using an in-house indent caller that excluded the first and last five bases of each read (www.github.com/kveeramah). The resulting VCF files contained diploid genotypes and genotype likelihoods. Coverage for autosomal 1240K SNPs and the mitochondrial genome was assessed for all individuals using GATK v4.2 DepthOfCoverage.^63^ The number of autosomal 1240K SNPs covered by at least one read with mapping and base quality scores of 30 or higher was determined for each BAM file using samtools depth.^64^ Genetic sex was inferred using the Sex.DetERRmine pipeline, which evaluates relative X and Y chromosome coverage (https://github.com/TCLamnidis/Sex.DetERRmine). Nuclear contamination rates in genetic males were estimated with ANGSD based on the hemizygous X chromosome,^65^ while mitochondrial contamination rates for all individuals were assessed using Schmutzi.^66^

We ran hapROH to estimate runs of homozygosity (ROH) in all 314 individuals from the LHP. The pseudohaploid PLINK files were converted to EIGENSTRAT format using Convertf (https://github.com/argriffing/eigensoft/tree/master/CONVERTF). We used the default global dataset from the 1000 Genomes Project (5008 haplotypes from 2504 individuals) as a reference panel, filtered to include only biallelic 1240K SNPs. The analysis was performed using the settings: min_cm=[4, 8, 12, 20], snp_cm=50, gap=0.5, min_len1=1.0, and min_len2=4.0. We counted all ROH segments longer than 5 cM.

### Mitochondrial and Y Chromosome Haplotype Analysis

For mitochondrial DNA analysis, BAM files from the newly sequenced individuals were filtered to retain only reads mapping to the mitochondrial genome, which were then converted to FASTQ format using samtools.^64^ Mitochondrial genomes were assembled for individuals with more than 1,500 mitochondrial reads using the Mapping Iterative Assembler,^67^ a tool specifically designed for ancient mitochondrial genome assembly. Haplotype assignment was then performed with MitoTool 1.1.2^68^ to determine mitochondrial haplogroups.

Y chromosome haplotypes were estimated based on the 1240K Y chromosome SNPs. Y chromosome VCF files were compiled in Excel, and the phylogenetic position of each Y chromosome SNP was determined using information from published databases (https://isogg.org/, accessed on 08/31/2023).^69–72^ This allowed for the identification of NRY haplogroups, named according to the updated ISOGG (International Society of Genetic Genealogy) nomenclature. Previously unclassified SNPs were assigned to haplogroups using a cladistic approach, accounting for the non-recombining nature of the NRY. Singletons were excluded unless already present in the referenced public databases.

### PCA

We conducted PCA by converting diploid 1240K VCF files to pseudohaploid plink datasets using a custom, in-house script. We integrated modern reference datasets, from imputed POPRES^28^ datasets. SmartPCA^73,74^ was used to perform PCA on the ancient individuals via a procrustes transformation (https://github.com/ShyamieG/). We used seaborn.kdeplot ^75^ function (default setting) to visualize the Kernel Density Estimation.

### Model-Based Clustering Analyses

Model-based clustering analysis was conducted using fastNGSadmix,^29^ applying a penecontemporaneous reference panel including 8 populations that developed from a previously published panel^31^, supplemented with an additional CASIA panel representing Central Asia. The Beagle PL files were generated using vcftools^76^ for all the 314 individuals at 1,082,095 autosomal sites for use by fastNGSadmix. The analysis was run separately for each individual using default parameters 30 times with random seeds, and the run with the highest likelihood chosen as the final result. The admixture plots shown in Fig. 2 utilize the point estimates given by the best run. The confidence intervals for admixture coefficients were also estimated using a simple block bootstrapping procedure.^77^ (Supplementary section S3)

### qpAdm and qpWave analysis

We performed qpWave/qpAdm (v. 1520) of the ADMIXTOOLS package^30^ analysis. The later was performed to validate the results obtained from fastNGSadmix. A random read-indent caller, which excluded the first and last eight bases of each read (https://github.com/kveeramah/), was applied in the analysis for all target, source, and reference populations. For the reference populations (right populations), we adopted 72 prehistoric individuals^26,78–87^ used in a previous study.^31^ The source populations were the same as the reference populations used in the fastNGSadmix analysis. For every test, we used a single individual from the LHP as the target. We first tested a one-source model, then progressively added up to three source populations in a stepwise manner. The process was finalized once the tail probability exceeded 0.01.

We also employed qpWave to model the source of our LHP target individuals using European source populations (Supplementary Table 2), and for the reference (right) populations, we used the same groups used in qpAdm, and we also used a cutoff of 0,01. To model the Roman-period LHP individuals (n=68), we used nine penecontemporaneous Eurasian reference populations. For the 6th-century individuals (n=246), we incorporated the Roman-period LHP populations and the Lake Balaton populations as two additional sources. To minimize biases related to sample size, we retained a maximum of 10 individuals per population and ensured all individuals were unrelated.

For qpAdm results using the same number of reference populations (one, two, or three), each individual modeled with n different combinations was assigned a weight of 1/n per combination. The final ancestry proportion for each reference population was calculated as the sum of its contributions across all combinations. For example, if an individual can be modeled using only one reference population—either NGBI or SCAND—the resulting ancestry is assigned equally: 0.5 NGBI and 0.5 SCAND. In a different case with two reference populations, suppose an individual is modeled with two two-source combinations, such as NGBI (0.3) / MEDEU (0.7) and SCAND (0.2) / MEDEU (0.8). Then, the final ancestry proportions would be 0.15 NGBI, 0.75 MEDEU, and 0.1 SCAND. The final results of genetic ancestry are in Supplementary Table 4 and illustrated in Figure 2b.

For the qpWave results, we used two complementary approaches to visualize the data. In the first approach, we excluded individuals who could not be modeled with any of the reference sources. For each remaining individual, we determined how many sources they could be modeled with and assigned equal proportions to each source (e.g., if an individual could be modeled with both Austria_2nd–5th c. and Balkans_1st–6th c., each source received 0.5). We then averaged the proportions across all individuals to obtain the final ancestry proportion for each period (Roman and post-Roman). To visualize the spatial distribution, we collected geographic coordinates for each source individual (noting that individuals from the same reference group share the same ancestry proportions). Using these data, we applied Inverse Distance Weighting (IDW) interpolation—implemented via the gstat package—to estimate ancestry proportions across mainland Europe. The resulting map is shown in Figure 3a, 3b (upper).

In the second approach, we calculated the proportion of individuals at each site who could be modeled with a given source (p>0.01), excluding Győr-Frigyes and Kóny due to their small sample sizes (n<15). These site-level results are presented a heatmap in Figure 3b (bottom).

For both qpAdm and qpWave analyses, we only plotted results for individuals with a 1240K coverage at least 0.01×.

### Biological relatedness

We utilized a customized version of the lcMLkin software package, following an approach similar in a previous study,^49^ to identify close biological relationships (within the third degree) among individuals at each site. The analysis initially included all individuals, regardless of coverage level, and was performed using genotype likelihood data from 1,076.758 autosomal 1240K sites.

To improve accuracy, allele frequency data from the 1000 Genomes CEU and TSI populations were used as outside population frequency. Relationships involving low-coverage individuals with fewer than 10,000 shared SNPs in an lcMLkin analysis were excluded, as they were likely to be spurious rather than indicative of genuine biological relatedness. To validate the lcMLkin results, we also conducted an independent analysis using KIN.^51^ The analysis was run with default parameters, with the -cnt parameter set to 0. The comparison between KIN and lcMLkin results can be found in Supplementary Table 9 column “lcMLkin_compare” and supplementary Section S7, and detailed description of pedigree reconstruction methods can be found in a previous study.^88^

### Haplotype IBD analysis

We implemented ancIBD^42^ to understand the identical-by-descent haplotype sharing pattern between individuals in the LHP and other contemporaneous populations (Supplementary Table 2) across Eurasia and African^31,39,40,49,88–102^.

Genotypes were called from whole-genome data and imputed for all diploid, dinucleotide sites within the 1000 Genomes Project Phase 3 v5a VCF files using GLIMPSE v1.1^103^, with the 1000 Genomes Project data serving as the imputation reference.^72^ The dataset was then filtered down to the 1,240K positions. Following the steps outlined in ancIBD, we converted VCF files to HDF5 format and identified IBD segments using default parameters. To enhance result reliability, we applied three filtering steps: first, excluding individuals with genomic coverage below 1×, and second, removing those with a GP99 ratio lower than 0.7, as recommended by the original study, finally, we only kept the connections with at least one shared IBD segments long than 12cM.

In the analysis focused on IBD sharing patterns among individuals from the Roman period, we have individuals in the LHP (n=31), 5th century sites of Fonyód, Hács and Balatonszemes around the Lake Balaton^31^ (n=26), and Bardonecchia/Torino-Lavazza^31^ (n=11).

For the post-Roman period, we have individuals in the LHP (n=130), Szólád^49^ (n=24), and Collegno^49,88^ (n=27). To construct the IBD network for post-Roman period individuals, we used Gephi v0.9.7^104^, applying the ForceAtlas2^105^ algorithm with "Prevent Overlap" enabled, an Edge_Weight_influence of 0.4, and Gravity set to 6. Edges were weighted by the adjusted Pi-hat value (calculated by dividing the sum_IBD by the total CM of the genome) and rescaled on a scale of 1 to 5. Node sizes were assigned based on rank, ranging from 10 to 30. Individuals with no IBD connections (degree=0) were excluded from the final network visualization.

### Local ancestry inference

We applied FLARE^44^ to infer local ancestry of individuals in the IBD network. We used modern populations from the 1000 Genomes Project^72^—CEU, GBR, TSI, AFR, EAS, and SAS—as reference populations for inferring local ancestry. To simplify analysis, we grouped the European populations into Northern European (nEUR, including CEU and GBR) and Southern European (TSI). We then analyzed raw IBD data generated by ancIBD^42^, filtering out segments with a density lower than 220/cM or shorter than 12 cM. For each remaining segment, we determined the ancestry status of both individuals using FALRE. At each SNP, we assigned a status based on the degree of ancestry matching between the two individuals across four chromosomes: (1) 1.0 if all four chromosomes share the same ancestry, (2) 0.5 if three chromosomes match, (3) 0.25 if only one chromosome from each individual matches. For each segment, we iterated through all SNPs, recording the lengths associated with each ancestry and their corresponding status. This process was repeated across all segments, with the final ancestry length for each segment calculated as length × status. To visualize the results, we merged all IBD segments for each ancestry and constructed ancestry-specific networks using Gephi.^104^ Extended Data Fig. 5 presents the resulting networks for nEUR and sEUR.

### Mantel test of burial distance and genetic distance

We performed a Mantel test ^106^ in R to assess the relationship between burial distance and genetic distance. Burial distance was calculated using geographic coordinates, while genetic distance was estimated based on pairwise differences in PC1 values, treated as a proxy for genetic proximity.

### Strontium isotope (^87^Sr/^86^Sr) analysis

Tooth enamel used for the Sr analysis was extracted in the laboratory of the Institute of Archaeogenomics, Budapest. For the extraction, first cleaning of the surface of the teeth was required for which a Dremel tool with an abrasion tip was used, followed by a ten-minute ultrasonic bath. After that, the enamel was carefully powdered with the help of a diamond-coated dental drill bit attached to the Dremel tool, until ca. 25–40 mg material was obtained. Approximately 20-30 mg of material from each sample was processed for strontium (Sr) isotope analysis in the MC-ICP-MS Facility, Department of Geological Sciences, University of Cape Town following routine procedures^107^ Enamel powder was weighed in Savillex PFA beakers, covered with 2 ml of concentrated HNO_3_ and closed beakers were placed on a hot plate at 140°C for an hour until complete dissolution. Beakers were opened, dried down and material redissolved in 1.5 ml 2M HNO_3_. Elemental Sr was isolated from these solutions using Triskem Sr.Spec resin and chemical protocols after Pin et al^108^. The isolated Sr fractions from each sample was diluted to 200 ppb Sr in 0.2% HNO_3_ and analysed on a Nu Instruments NuPlasma HR MC-ICP-MS, with reported ^87^Sr/^86^Sr data referenced to bracketing analyses of the international standard reference NIST SRM987 using an ^87^Sr/^86^Sr value of 0.710255^109^. All Sr isotope data were corrected for Rb interference at 87 amu using the measured ^85^Rb signal and the natural ^85^Rb/^87^Rb abundance ratio. Instrumental mass fractionation was corrected using the exponential law and an ^86^Sr/^88^Sr value of 0.1194. Results of repeat analyses of both an in-house carbonate reference material NM95 (^87^Sr/^86^Sr 0.708897±0.000017 2s; n=5) and international reference material NIST SRM1400 (^87^Sr/^86^Sr 0.713134±0.000037 2s; n=4) analysed as unknowns with samples during this study agree with long-term results from this facility (NM95 ^87^Sr/^86^Sr 0.708909±0.000038 2s; n=932, SRM1400 ^87^Sr/^86^Sr 0.713122±0.000036 2s; n=53) and limited published data (SRM1400 ^87^Sr/^86^Sr 0.71308-0.7134; GeoReM database). Elemental Sr measured in total procedural blanks were typical for this facility at < 250 pg and therefore negligible.

### Carbon and Nitrogen isotope analyses

C and N isotope analysis (δ13C, δ15N) was conducted on collagen preferably from ribs, if those were not available on long bones (humerus, femur, tibia, or fibula) at the Curt Engelhorn Centre Archaeometry in Mannheim, Germany or at the Department of Applied and Analytical Palaeontology, Institute of Geosciences at the University Mainz, Germany. Bone samples were cleaned, chemically treated and collagen extracted using a modified Longin method^110^ with modifications as described in Knipper et al.^111^. C and N contents and the stable isotopic compositions were determined in triplicates using a Thermo Flash 2000 Organic Elemental Analyzer coupled to a Thermo Finnigan Mat 253 mass spectrometer or a vario PYRO cube CNSOH elemental analyzer (Elementar) and a precisION isotope ratios mass spectrometer (Isoprime) depending on the lab. Then the raw data were calibrated against the international Standards.

### Radiocarbon dating

A total of 38 human bone samples were radiocarbon dated to provide absolute chronological anchors for the individuals analyzed. All samples were prepared and measured at the Curt-Engelhorn-Center Archaeometry in Mannheim, Germany ^112^. Collagen was extracted following a modified Longin protocol with an additional ultrafiltration step to remove low-molecular-weight contaminants ^113^. Quality control criteria included collagen yield, elemental composition (%C, %N), and atomic C:N ratio. All samples yielded sufficient collagen for AMS measurement, with C:N atomic ratios falling within the accepted range for well-preserved collagen (2.9–3.6; ^114^). Accepted collagen extracts were combusted in an Elementar VarioMicro elemental analyzer interfaced with either an in-house semi-automated graphitization system (MAG) or a commercial AGE3 system (Ionplus AG). The resulting graphite was measured on one of two MICADAS accelerators. Radiocarbon measurements were normalized using Oxalic Acid II (NIST SRM 4990C) as the primary standard. Each batch additionally included process blanks and quality control samples of known radiocarbon age—typically IAEA reference materials or well-characterized bone material routinely used in the laboratory—to monitor background levels, instrument stability, and measurement accuracy. The resulting conventional radiocarbon ages were corrected for isotopic fractionation and normalized to a δ¹³C value of –25‰. Radiocarbon ages were calibrated using the IntCal20 calibration curve ^115^ in OxCal v4.4.4 ^116^.

---

**Extended Data Fig. 1.**
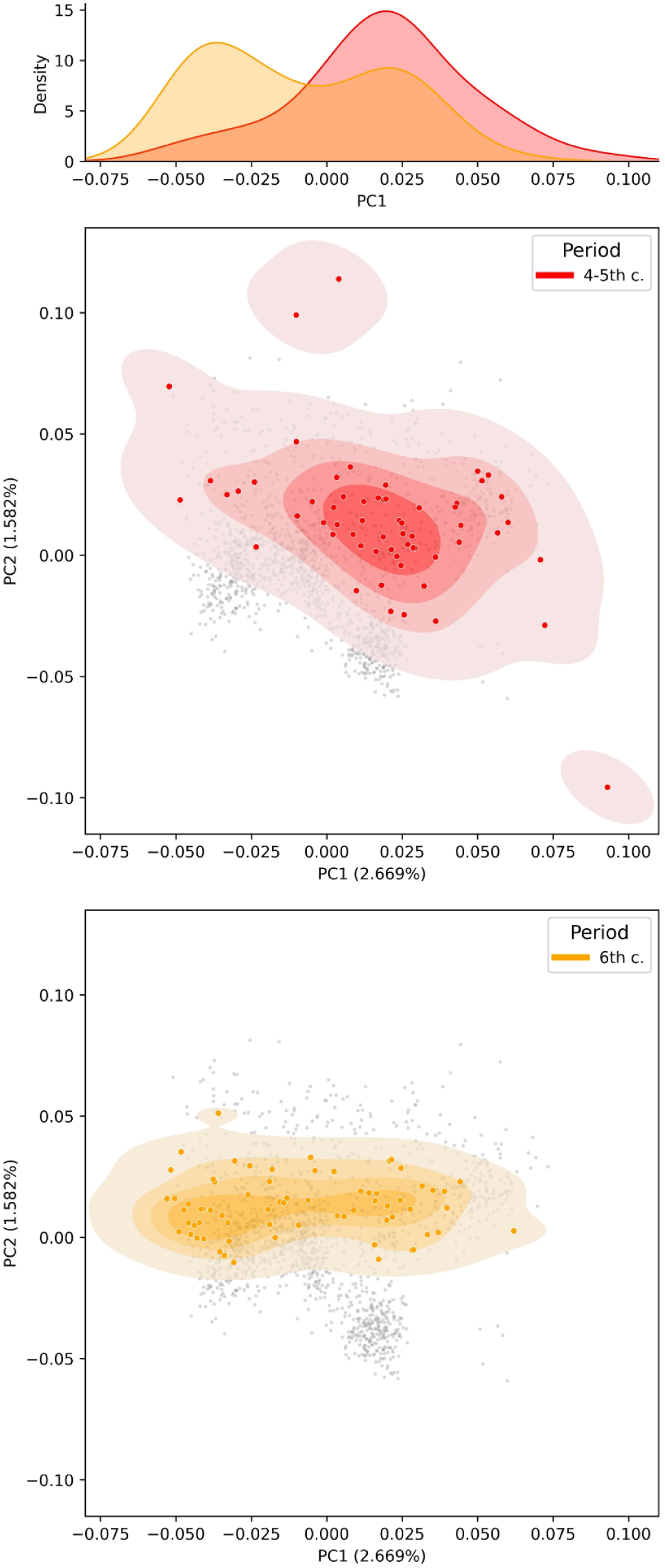
PCA results for 4-5th c. sites (n=68) and a downsampled version of the 6th c. sites (n = 68).

**Extended Data Fig. 2.**
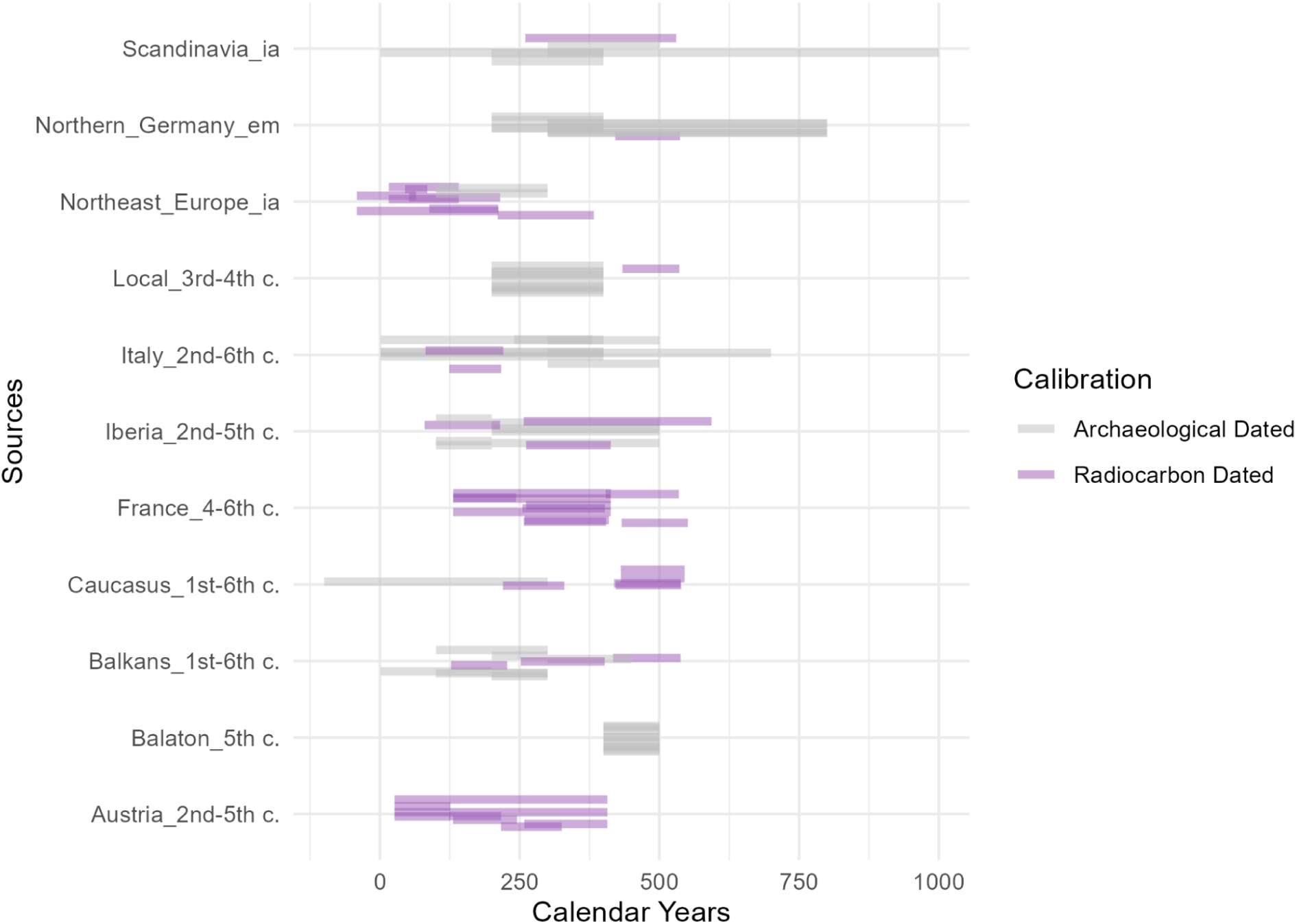
Dating ranges of source individuals and groups included in the qpWave analysis. Each line represents an individual, with colors indicating whether the date is radiocarbon-calibrated. Ia: Iron Age; em: Early Medieval

**Extended Data Fig. 3.**
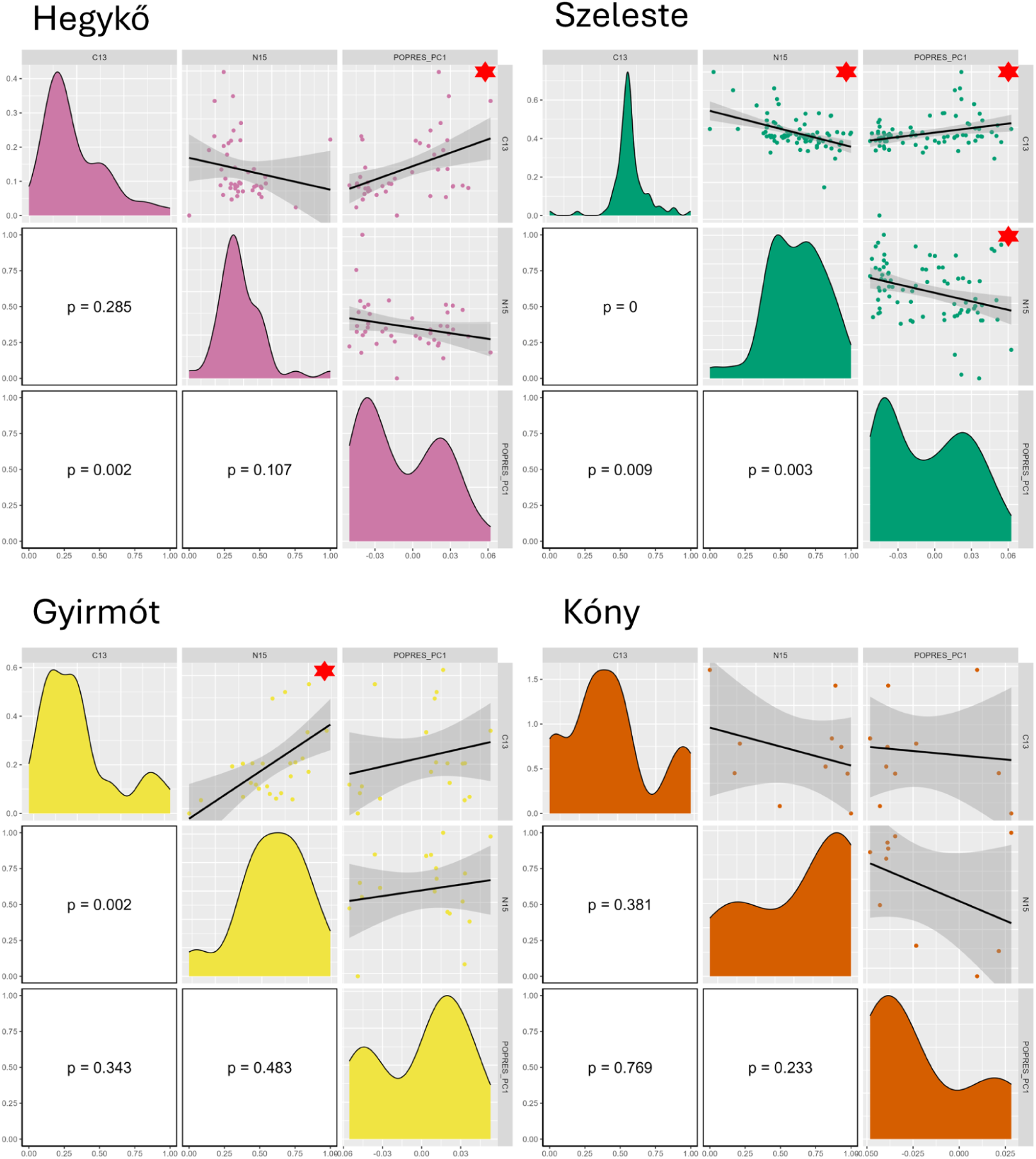
The distribution and linear relationships between PC1, δ¹⁵N, and δ¹³C and PC1 values across four sites. Asterisks indicate statistically significant p-values (threshold=0.05), with exact p-values displayed in the lower left corner.

**Extended Data Fig. 4.**
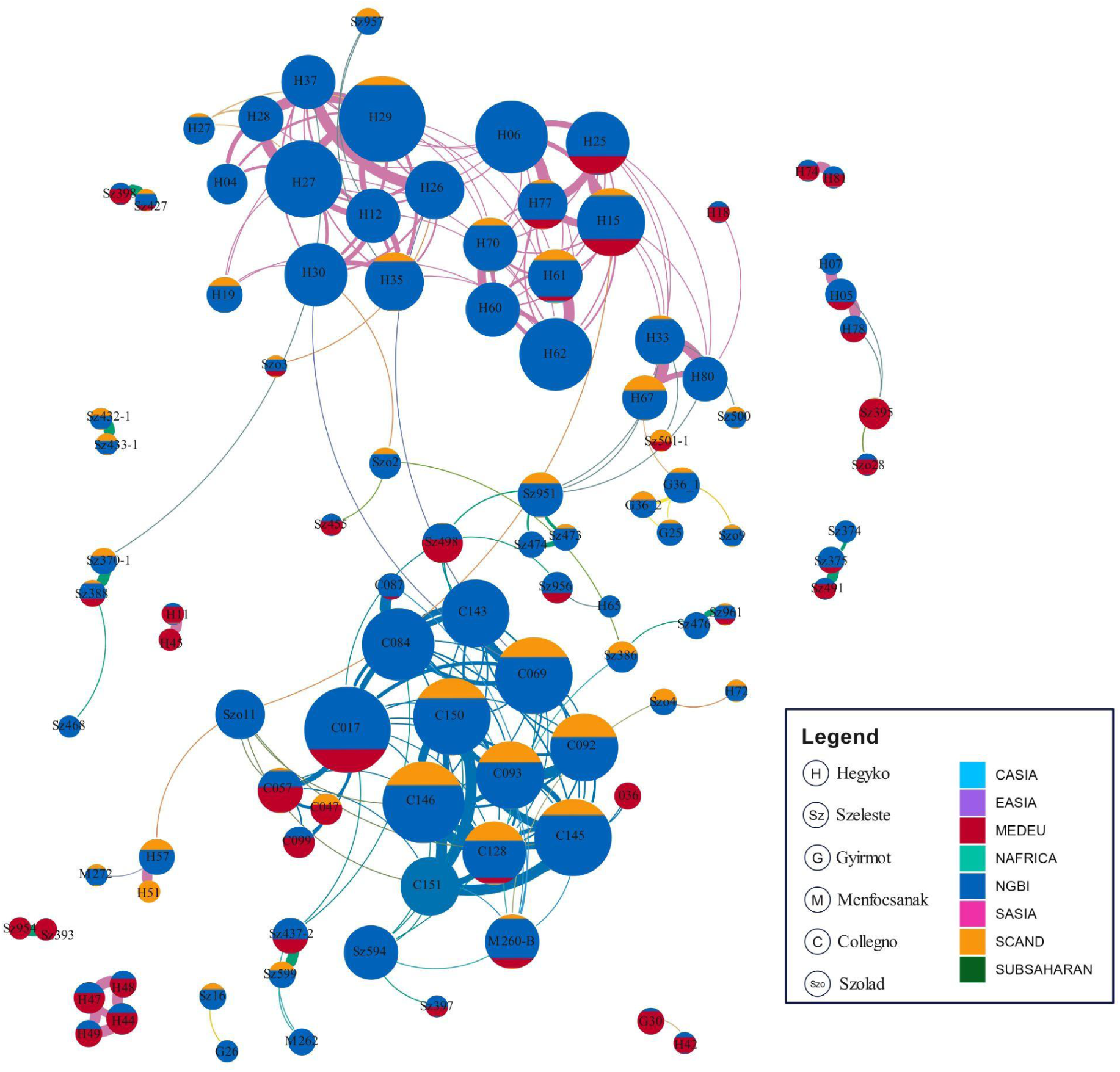
The same IBD network as in Fig. 3c, but colored according to genetic ancestry inferred from fastNGSadmix results.

**Extended Data Fig. 5.**
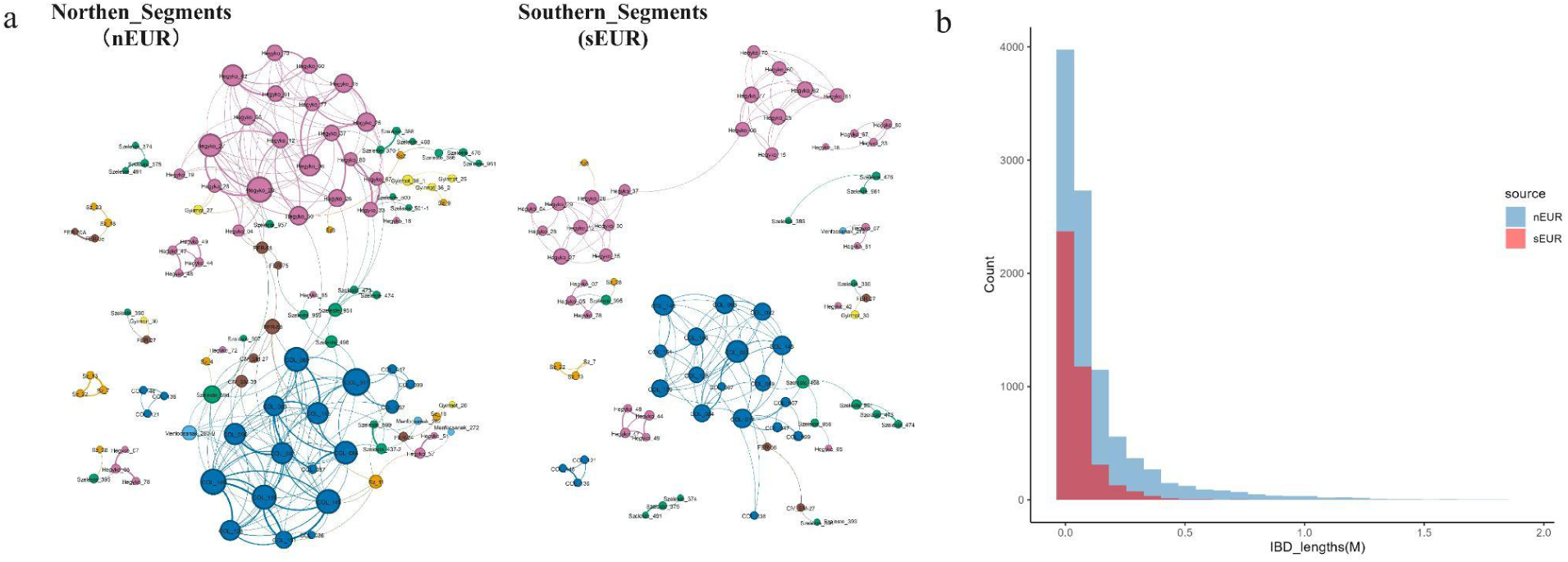
Local Ancestry Inference results using FLARE. a, IBD network based on both Northern ancestry segments (nEUR) and Southern ancestry Segments (sEUR). b, The distribution of nEUR and sEUR.

**Extended Data Fig. 6.**
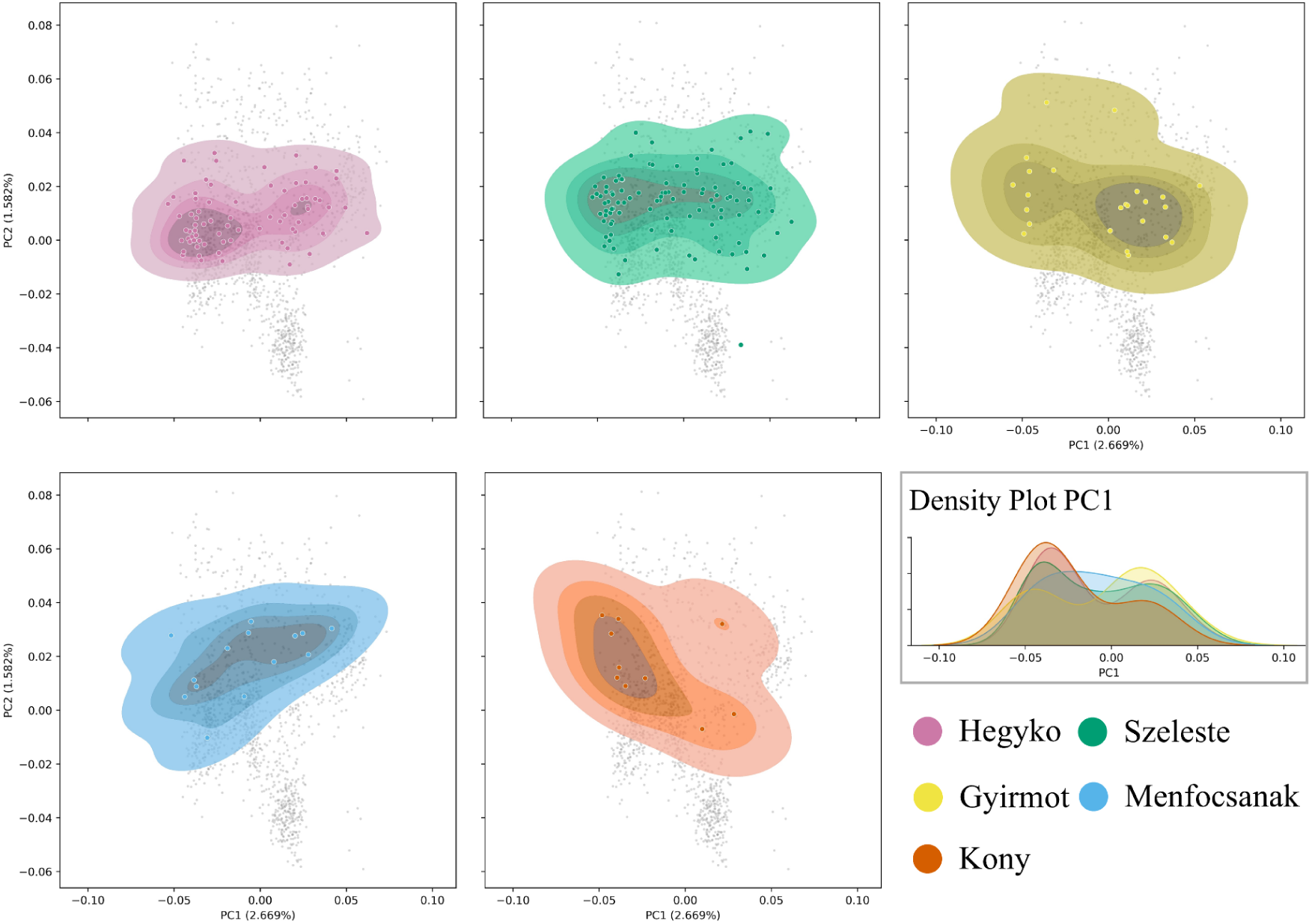
PCA of five 6^th^ century CE sites in the Little Hungarian Plain. It used the same POPRES reference panel and methods as in the Fig. 2.

**Extended Data Fig. 7.**
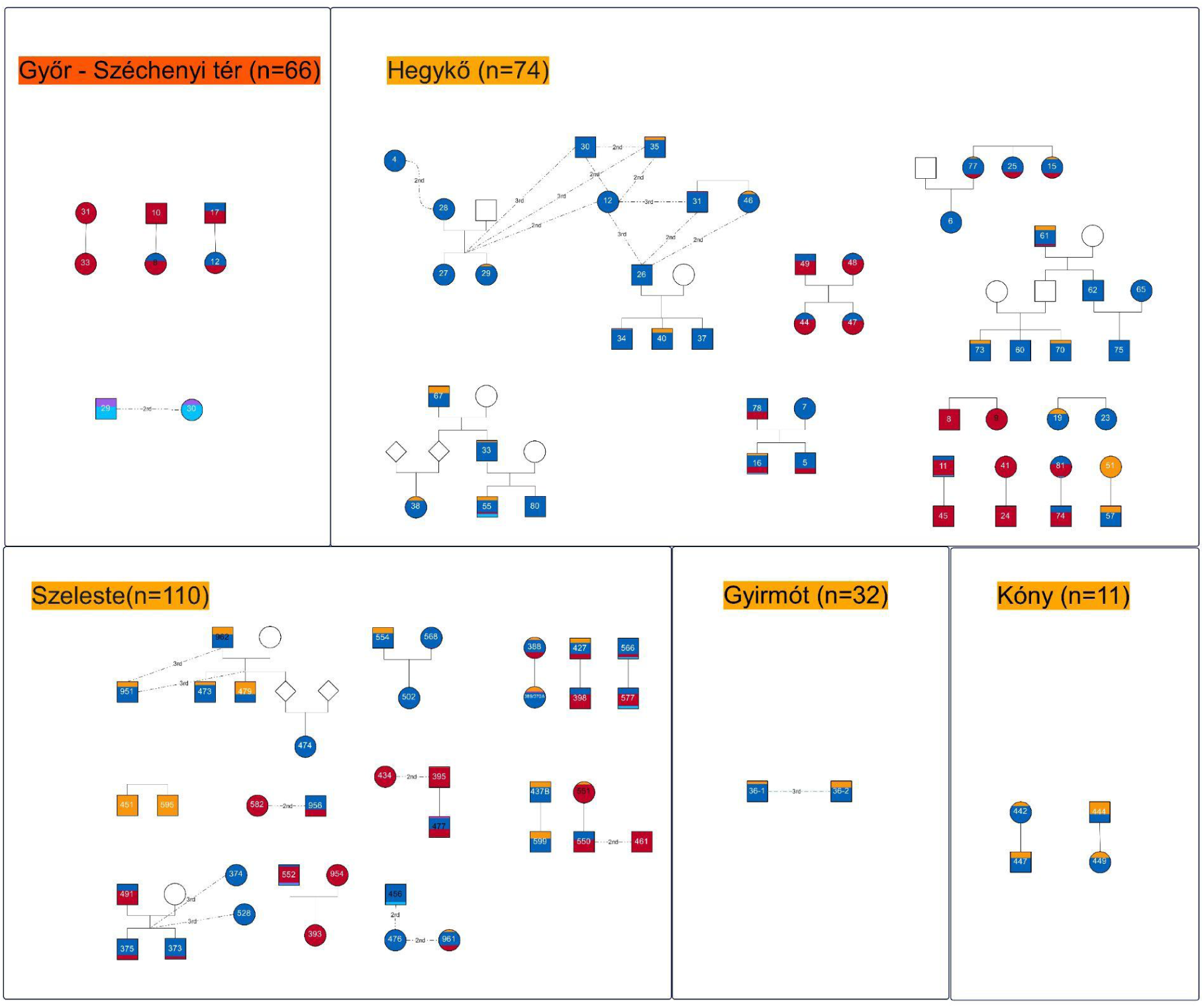
Pedigrees identified at the 6th-century sites. The filled colors of individuals represent fastNGSadmix results from penecontemporaneous panels. Circles indicate females, squares indicate males. Only relationships within third-degree relatedness are shown.

**Extended Data Fig. 8.**
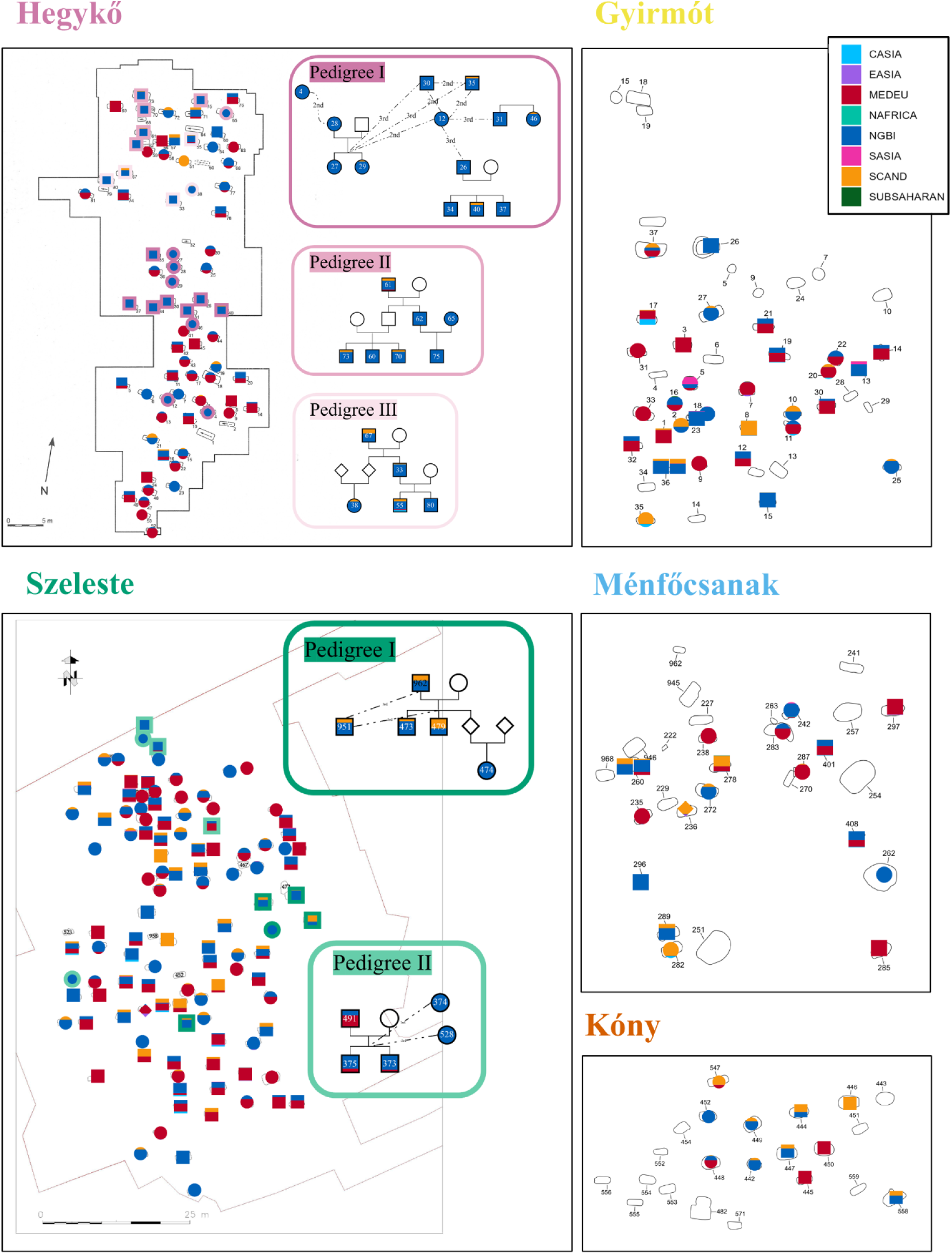
Cemetery maps combined with genetic profiles for individuals in all the five 6^th^ century CE sites. Burials are marked by genetic ancestries derived from fastNGSadmix analysis, with circles representing females and squares representing males. Large pedigrees (n>4) are highlighted, with scaled colors used to distinguish them within each cemetery map.

## Notes

### Competing Interest Statement

The authors have declared no competing interest.

## References

1. Halsall, G. *Barbarian Migrations and the Roman West,* 376–568. (Cambridge University Press, Cambridge, England, 2007).

2. Wickham, C. *Framing the Early Middle Ages: Europe and the Mediterranean*, 400-800. (OUP Oxford, 2006).

3. Meier, M. *Die Geschichte der Völkerwanderung: Europa, Asien und Afrika vom 3. bis zum 8*. Jahrhundert n.Chr. 7. Auflage. (C.H.Beck, Munich, Germany, 2021).

4. Vida, T. Die Zeit zwischen dem 4. und dem 6. Jahrhundert im mittleren Donauraum aus archäologischer Sicht. in Römische Legionslager in den Rhein- und Donauprovinzen - Nuclei spätantik-frühmittelalterlichen Lebens? (eds. Witschel, C. & Konrad, M.) 615–650 (Bayerischen Akademie der Wissenschaften bei C.H.Beck, München, 2011).

5. Hardt, M. Pannonien im Spannungsfeld zwischen Römer- und Völkerwanderungszeit - eine geschichtliche Einführung. in Keszthely-Fenékpuszta im Kontext spätantiker Kontinuitätsforschung zwischen Noricum und Moesia (ed. Heinrich-Tamáska, O.) 15–28 (Verlag Marie Leidorf, Rahden, Westf., 2011).

6. Tomka, P. Langobardenforschung in Nordwestungarn. in Die Langobarden. Herrschaft und Identität. Ergebnisse eines vom 2. bis 4. November 2001 in Wien abgehaltenen internationalen Symposiums. (eds. Pohl, W. & Erhart, P.) 247–264 (Verlag der österreichischen Akademie der Wissenschaften, Vienna, 2005).

7. Tomka, P. Kulturwechsel der spätantiken Bevölkerung eines Auxiliarcastells: Fallbeispiel Arrabona. in Zentrum und Peripherie – Gesellschaftliche Phänomene in der Frühgeschichte (eds. Friesinger, H. & Stuppner, A.) 389–409 (Verlag der österreichischen Akademie der Wissenschaften, Vienna, 2004).

8. Tomka, P. Eine römische Stadt und ihre barbarische Peripherie: Scarbantia. in Romania Gothica II. The Frontier World. Romans, Barbarians and Military Culture (ed. Vida, T.) 587–615 (Martin Optiz Kiadó, Budapest, 2015).

9. Pohl, W. Migration und Ethnogenesen der Langobarden aus Sicht der Schriftquellen. in Kulturwandel in Mitteleuropa: Langobarden, Awaren, Slawen. Akten der internationalen Tagung in Bonn vom 25. bis 28. Februar 2008 (eds. Bemmann, J. & Schmauder, M.) 1–12 (Habelt, R, Bonn, Germany, 2008).

10. Vida, T. Aufgaben und Perspektiven der Langobardenforschung in Ungarn nach István Bóna. in Kulturwandel in Mitteleuropa : Langobarden, Awaren, Slawen : Akten der Internationalen Tagung in Bonn vom 25. bis 28. Februar 2008 (eds. Schmauder, M. & Bemmann, J.) 343–362 (Deutsches Archäologisches Institut. Römisch-Germanische Kommission (Berlin) by Habelt, Bonn, 2008).

11. Bóna, I. A langobardok története és régészeti emlékei. in Hunok - Gepidák - Langobardok (eds. Bóna, I., Cseh, J., Nagy, M., Tomka, P. & Tóth, Á.) 102–162 (JATE Magyar Őstörténeti Kutatócsoport, Szeged, 1993).

12. Koncz, I. A hegykői 6. Századi temető időrendje és kapcsolatrendszere. Archaeol. Ért. 139, 71–98 (2014).

13. Bendeguz, T., Wiltschke-Schrotta, K. & Binder, M. The Langobardian period cemetery of Vienna-Mariahilfer Gürtel. With a contribution to artificial cranial deformation in the western Carpathian Basin. Jahrbuch des Römisch-Germanischen Zentralmuseums Mainz 57, (2014).

14. Vida, T. Late Roman territorial organisation and the settlement of the barbarian gentes in Pannonia. Hortus Artium Mediev. 13, 319–331 (2007).

15. Virágos, R. Tájrégészeti megközelítések a dunántúli 5-6. századi régészeti lelőhelyek értelmezésében. Archaeol. Ért. 133, 199–221 (2008).

16. Hajdu, T. et al. Bone tuberculosis in Roman Period Pannonia (western Hungary). Mem. Inst. Oswaldo Cruz 107, 1048–1053 (2012).

17. Bíró, S. & Tomka, P. Über die mysteriöse ‘schwarze Schicht’ und das sog. ‘hunnenzeitliche Gräberfeld’ von Győr-Széchenyi Platz. in Na hranicích impéria = Extra fines imperii: Jaroslavu Tejralovi k 80. narozeninám 53–72 (Masarykova univerzita – Archeologický ústav AV ČR, Brno, Brno, 2017).

18. Koncz, I. 568 — A historical date and its archaeological consequences. Acta Archaeol. 66, 315–340 (2015).

19. Tóth, G. & Pap, I. K. Germán temető a Nyugat-Dunántúlon. in Trendek és eredmények a biológiai kutatás és oktatás terén (eds. Nagy, M. & Poráčová, J.) 22–27 (J. Selye University, Komarno, 2016).

20. Pap, I. K. Savaria keleti temetője és a szelestei germán temető épített és tegulás sírjai (Savaria’s eastern cemetery and the built and tegula tombs of the Szeleste Langobard cemetery). Savaria 38, 91–105 (2016).

21. Vaday, A. The Langobard cemetery at Ménfőcsanak. Antaeus 33, 163–242 (2015).

22. Vida, T. Spätantike Buntmetallgefäβe im langobardenzeitlichen Pannonien. in Macht des Goldes, Gold der Macht. Herrschafts und Jenseitsrepräsentation zwischen Antike und Frühmittelalter im Mittleren Donauraum. Aktend des 23. Internationalen Symposiums der Grundprobleme der frühgeschichtlichen Entwicklung im mittleren Donauraum, Tengelic, 16.-19.11.2011. (eds. Hardt, M. & Heinrich-Tamáska, O.) 339–360 (Verlag Bernhard Albert Greiner, Weinstadt, 2013).

23. Horváth, E. Gemstone and glass inlaid fine metalwork from the Carpathian Basin: the Hunnic and Early Merovingian Periods. Diss. archaeol. ex Inst. Archaeol. Univ. Rolando Eötvös Nomin. 275–302 (2013).

24. Koncz, I. About brooches and networks: Some remarks on the female dress in 6th century Pannonia. Diss. archaeol. ex Inst. Archaeol. Univ. Rolando Eötvös Nomin. 163–176 (2018).

25. Rohland, N., Harney, E., Mallick, S., Nordenfelt, S. & Reich, D. Partial uracil-DNA-glycosylase treatment for screening of ancient DNA. Philos. Trans. R. Soc. Lond. B Biol. Sci. 370, 20130624 (2015).

26. Mathieson, I. et al. Genome-wide patterns of selection in 230 ancient Eurasians. Nature 528, 499–503 (2015).

27. Fu, Q. et al. DNA analysis of an early modern human from Tianyuan Cave, China. Proc. Natl. Acad. Sci. U. S. A. 110, 2223–2227 (2013).

28. Nelson, M. R. et al. The Population Reference Sample, POPRES: a resource for population, disease, and pharmacological genetics research. Am. J. Hum. Genet. 83, 347–358 (2008).

29. Jørsboe, E. & Hanghøj, K. Albrechtsen, fastNGSadmix: Admixture proportions and principal component analysis of a single NGS sample. Bioinformatics 33, 3148–3150 (2017).

30. Patterson, N. et al. Ancient admixture in human history. Genetics 192, 1065–1093 (2012).

31. Vyas, D. N. et al. Fine-scale sampling uncovers the complexity of migrations in 5th-6th century Pannonia. Curr. Biol. 33, 3951–3961.e11 (2023).

32. Antonio, M. L. et al. Stable population structure in Europe since the Iron Age, despite high mobility. bioRxiv 2022.05.15.491973 (2022) doi:10.1101/2022.05.15.491973.

33. Olalde, I. et al. A genetic history of the Balkans from Roman frontier to Slavic migrations. Cell 186, 5472–5485.e9 (2023).

34. Lazaridis, I. et al. A genetic probe into the ancient and medieval history of Southern Europe and West Asia. Science 377, 940–951 (2022).

35. Olalde, I. et al. The genomic history of the Iberian Peninsula over the past 8000 years. Science 363, 1230–1234 (2019).

36. Posth, C. et al. The origin and legacy of the Etruscans through a 2000-year archeogenomic time transect. Sci. Adv. 7, eabi7673 (2021).

37. Antonio, M. L. et al. Ancient Rome: A genetic crossroads of Europe and the Mediterranean. Science 366, 708–714 (2019).

38. Stolarek, I. et al. Genetic history of East-Central Europe in the first millennium CE. Genome Biol. 24, 173 (2023).

39. Gretzinger, J. et al. The Anglo-Saxon migration and the formation of the early English gene pool. Nature 610, 112–119 (2022).

40. Margaryan, A. et al. Population genomics of the Viking world. Nature 585, 390–396 (2020).

41. Ringbauer, H., Novembre, J. & Steinrücken, M. Parental relatedness through time revealed by runs of homozygosity in ancient DNA. Nat. Commun. 12, 5425 (2021).

42. Ringbauer, H. et al. Accurate detection of identity-by-descent segments in human ancient DNA. Nat. Genet. 56, 143–151 (2024).

43. McColl, H., et al. Steppe Ancestry in western Eurasia and the spread of the Germanic Languages. bioRxiv (2024) doi:10.1101/2024.03.13.584607.

44. Browning, S. R., Waples, R. K. & Browning, B. L. Fast, accurate local ancestry inference with FLARE. Am. J. Hum. Genet. 110, 326–335 (2023).

45. Pohl, W. Alboin und der Langobardenzug nach Italien. Aufstieg und Fall eines Barbarenkönigs. in Sie schufen Europa. Historische Portraits von Konstantin bis Karl dem Großen (ed. Meier, M. (hrsg ).) 216–227 (C.H.Beck, München, 2007).

46. Christie, N. The Lombards: The Ancient Longobards. (Wiley, 1999).

47. Jarnut, J. Geschichte der Langobarden. (Kohlhammer, Stuttgart, Germany, 1982).

48. Knipper, C. et al. Coalescing traditions-Coalescing people: Community formation in Pannonia after the decline of the Roman Empire. PLoS One 15, e0231760 (2020).

49. Amorim, C. E. G. et al. Understanding 6th-century barbarian social organization and migration through paleogenomics. Nat. Commun. 9, 3547 (2018).

50. Lipatov, M., Sanjeev, K., Patro, R. & Veeramah, K. R. Maximum Likelihood Estimation of Biological Relatedness from Low Coverage Sequencing Data. bioRxiv 023374 (2015) doi:10.1101/023374.

51. Popli, D., Peyrégne, S. & Peter, B. M. KIN: a method to infer relatedness from low-coverage ancient DNA. Genome Biol. 24, 10 (2023).

52. Jarnut, J. Langobardische Eliten in der sozialen Praxis. in Théorie et pratiques des élites au Haut Moyen Âge. Conception, perception et réalisation sociale (eds. Bougard, F., Goetz, H.-W. & Le Jan, R.) vol. 13 283–290 (Brepols, 2011).

53. *Monumenta Germaniae Historica. Scriptores Rerum Langobardicarum et Italicarum Saec* VI-IX. (Hannover, 1878).

54. Bognetti, G. P. *L’influsso Delle Istituzioni Militari Romane Sulle Istituzioni Longobarde Del Secolo VI E La Natura Della ‘fara’*. (Milano, 1967).

55. Murray, A. C. *Germanic Kinship Structure. Studies in Law and Society in Antiquity and in the Early Middle Ages*. (Pontifical Institute of Medieval Studies, Toronto, 1983).

56. Haubrichs, W. »Leudes, fara, faramanni und farones«: Zur Semantik der Bezeichnungen für einige am Konsenshandeln beteiligte Gruppen. (2020) doi:10.11588/VUF.2017.0.71813.

57. *Plague and the End of Antiquity*. (Cambridge University Press, Cambridge, England, 2006).

58. Fellows Yates, J. A., et al. Reproducible, portable, and efficient ancient genome reconstruction with nf-core/eager. PeerJ 9, e10947 (2021).

59. Schubert, M., Lindgreen, S. & Orlando, L. AdapterRemoval v2: rapid adapter trimming, identification, and read merging. BMC Res. Notes 9, 88 (2016).

60. Li, H. & Durbin, R. Fast and accurate short read alignment with Burrows-Wheeler transform. Bioinformatics 25, 1754–1760 (2009).

61. Li, H. et al. The Sequence Alignment/Map format and SAMtools. Bioinformatics 25, 2078–2079 (2009).

62. Jónsson, H., Ginolhac, A., Schubert, M., Johnson, P. L. F. & Orlando, L. mapDamage2.0: fast approximate Bayesian estimates of ancient DNA damage parameters. Bioinformatics 29, 1682–1684 (2013).

63. Mckenna, A. The genome analysis toolkit: A mapreduce framework for analyzing nextgeneration DNA sequencing data. Genome Res 20, 1297–1303 (2010).

64. Danecek, P. et al. Twelve years of SAMtools and BCFtools. Gigascience 10, (2021).

65. Korneliussen, T. S., Albrechtsen, A. & Nielsen, R. ANGSD: Analysis of next generation sequencing data. BMC Bioinformatics 15, 356 (2014).

66. Renaud, G., Slon, V., Duggan, A. T. & Kelso, J. Schmutzi: estimation of contamination and endogenous mitochondrial consensus calling for ancient DNA. Genome Biol. 16, 224 (2015).

67. Briggs, A. W. et al. Targeted retrieval and analysis of five Neandertal mtDNA genomes. Science 325, 318–321 (2009).

68. Fan, L. & Yao, Y.-G. An update to MitoTool: using a new scoring system for faster mtDNA haplogroup determination. Mitochondrion 13, 360–363 (2013).

69. Francalacci, P. Low-pass DNA sequencing of 1200 Sardinians reconstructs European Ychromosome phylogeny. Science 341, 565–569 (2013).

70. Karmin, M. et al. A recent bottleneck of Y chromosome diversity coincides with a global change in culture. Genome Res. 25, 459–466 (2015).

71. Poznik, G. D. et al. Sequencing Y chromosomes resolves discrepancy in time to common ancestor of males versus females. Science 341, 562–565 (2013).

72. 1000 Genomes Project Consortium et al. A global reference for human genetic variation. Nature 526, 68–74 (2015).

73. Price, A. L. et al. Principal components analysis corrects for stratification in genome-wide association studies. Nat. Genet. 38, 904–909 (2006).

74. Patterson, N., Price, A. L. & Reich, D. Population structure and eigenanalysis. PLoS Genet. 2, e190 (2006).

75. Waskom, M. seaborn: statistical data visualization. J. Open Source Softw. 6, 3021 (2021).

76. Danecek, P. et al. The variant call format and VCFtools. Bioinformatics 27, 2156–2158 (2011).

77. Kunsch, H. R. The Jackknife and the Bootstrap for General Stationary Observations. Ann. Stat. 17, 1217–1241 (1989).

78. Lazaridis, I. et al. Genomic insights into the origin of farming in the ancient Near East. Nature 536, 419–424 (2016).

79. Lazaridis, I. et al. Ancient human genomes suggest three ancestral populations for present-day Europeans. Nature 513, 409–413 (2014).

80. Haak, W. et al. Massive migration from the steppe was a source for Indo-European languages in Europe. Nature 522, 207–211 (2015).

81. Allentoft, M. E. et al. Population genomics of Bronze Age Eurasia. Nature 522, 167–172 (2015).

82. Gamba, C. et al. Genome flux and stasis in a five millennium transect of European prehistory. Nat Commun 5: 5257. Preprint at (2014).

83. Hofmanová, Z. et al. Early farmers from across Europe directly descended from Neolithic Aegeans. Proc. Natl. Acad. Sci. U. S. A. 113, 6886–6891 (2016).

84. Mathieson, I. et al. The genomic history of southeastern Europe. Nature 555, 197–203 (2018).

85. Olalde, I. et al. Derived immune and ancestral pigmentation alleles in a 7,000-year-old Mesolithic European. Nature 507, 225–228 (2014).

86. van de Loosdrecht, M. et al. Pleistocene North African genomes link Near Eastern and sub-Saharan African human populations. Science 360, 548–552 (2018).

87. Fu, Q. et al. The genetic history of Ice Age Europe. Nature 534, 200–205 (2016).

88. Tian, Y. et al. The role of emerging elites in the formation and development of communities after the fall of the Roman Empire. Proc. Natl. Acad. Sci. U. S. A. 121, e2317868121 (2024).

89. Antonio, M. L. et al. Stable population structure in Europe since the Iron Age, despite high mobility. Elife 13, e79714 (2024).

90. Damgaard, P. de B., et al. 137 ancient human genomes from across the Eurasian steppes. Nature 557, 369–374 (2018).

91. Flegontov, P. et al. Palaeo-Eskimo genetic ancestry and the peopling of Chukotka and North America. Nature 570, 236–240 (2019).

92. Gnecchi-Ruscone, G. A. et al. Ancient genomic time transect from the Central Asian Steppe unravels the history of the Scythians. Sci. Adv. 7, eabe4414 (2021).

93. Gnecchi-Ruscone, G. A. et al. Network of large pedigrees reveals social practices of Avar communities. Nature 629, 376–383 (2024).

94. Harney, É. et al. Ancient DNA from the skeletons of Roopkund Lake reveals Mediterranean migrants in India. Nat. Commun. 10, 3670 (2019).

95. Jeong, C. et al. A dynamic 6,000-year genetic history of Eurasia’s Eastern Steppe. Cell 183, 890–904.e29 (2020).

96. Lazaridis, I. et al. The genetic history of the Southern Arc: A bridge between West Asia and Europe. Science 377, eabm4247 (2022).

97. Marcus, J. H. et al. Genetic history from the Middle Neolithic to present on the Mediterranean island of Sardinia. Nat. Commun. 11, 939 (2020).

98. Maróti, Z. et al. The genetic origin of Huns, Avars, and conquering Hungarians. Curr. Biol. 32, 2858–2870.e7 (2022).

99. Patterson, N. et al. Large-scale migration into Britain during the Middle to Late Bronze Age. Nature 601, 588–594 (2022).

100. Sikora, M. et al. The population history of northeastern Siberia since the Pleistocene. Nature 570, 182–188 (2019).

101. Veeramah, K. R. et al. Population genomic analysis of elongated skulls reveals extensive female-biased immigration in Early Medieval Bavaria. Proc. Natl. Acad. Sci. U. S. A. 115, 3494–3499 (2018).

102. Wang, C.-C. et al. Genomic insights into the formation of human populations in East Asia. Nature 591, 413–419 (2021).

103. Rubinacci, S., Ribeiro, D. M., Hofmeister, R. J. & Delaneau, O. Publisher Correction: Efficient phasing and imputation of low-coverage sequencing data using large reference panels. Nat. Genet. 53, 412 (2021).

104. Bastian, M., Heymann, S. & Jacomy, M. Gephi: An Open Source Software for Exploring and Manipulating Networks. ICWSM 3, 361–362 (2009).

105. Jacomy, M., Venturini, T., Heymann, S. & Bastian, M. ForceAtlas2, a continuous graph layout algorithm for handy network visualization designed for the Gephi software. PLoS One 9, e98679 (2014).

106. Oksanen, J., et al. Vegan: Community Ecology Package. (2020).

107. Scott, M., Le Roux, P., Sealy, J. & Pickering, R. Lead and strontium isotopes as palaeodietary indicators in the Western Cape of South Africa. S. Afr. J. Sci. 116, (2020).

108. Pin, C., Gannoun, A. & Dupont, A. Rapid, simultaneous separation of Sr, Pb, and Nd by extraction chromatography prior to isotope ratios determination by TIMS and MC-ICP-MS. J. Anal. At. Spectrom. 29, 1858–1870 (2014).

109. Waight, T., Baker, J. & Peate, D. Sr isotope ratio measurements by double-focusing MC-ICPMS: techniques, observations and pitfalls. Int. J. Mass Spectrom. 221, 229–244 (2002).

110. Longin, R. New method of collagen extraction for radiocarbon dating. Nature 230, 241–242 (1971).

111. Knipper, C. et al. What is on the menu in a Celtic town? Iron Age diet reconstructed at Basel-Gasfabrik, Switzerland. Archaeol. Anthropol. Sci. 9, 1307–1326 (2017).

112. Kromer, B., Lindauer, S., Synal, H.-A. & Wacker, L. MAMS – A new AMS facility at the Curt-Engelhorn-Centre for Achaeometry, Mannheim, Germany. Nuclear Instruments and Methods in Physics Research Section B: Beam Interactions with Materials and Atoms vol. 294 11–13 Preprint at 10.1016/j.nimb.2012.01.015 (2013).

113. Brown, T. A., Nelson, D. E., Vogel, J. S. & Southon, J. R. Improved Collagen Extraction by Modified Longin Method. Radiocarbon 30, 171–177 (1988).

114. DeNiro, M. J. Postmortem preservation and alteration of in vivo bone collagen isotope ratios in relation to palaeodietary reconstruction. Nature 317, 806–809 (1985).

115. Reimer, P. J. et al. The IntCal20 Northern hemisphere radiocarbon age calibration curve (0–55 cal kBP). Radiocarbon 62, 725–757 (2020).

116. Ramsey, C. B. Bayesian Analysis of Radiocarbon Dates. Radiocarbon 51, 337–360 (2009).

