## Supplementary Information for "Unveiling the complexity of post-Roman polity formation using ancient DNA"

### S.1. Description of sites

**S1.1. Gyirmót,** Győr–Moson–Sopron County, Hungary

mid 6th century CE

37 graves, 37 individuals, 36 aDNA samples

The site was excavated in 1994 as part of the rescue excavations between 1990 and 2012 connected to the building of the main road 83. The site is located on an elevated ridge near the Roman road in the vicinity of the Roman town Arrabona (today's Győr). By the 6th century, Arrabona lost its urban character and its military and administrative role, it is even questionable to what extent was it inhabited during this period, so the small site at Gyirmót probably represents a smaller, rural community that settled along the Roman road connecting Arrabona and Scarbantia. The cemetery is completely excavated. Most of the unearthed graves showed signs of contemporary disturbance: the traces of secondary openings were clearly observable in the soil, skeletons were generally out of position, only parts remained in situ. Due to the disturbance, the archaeological material is also fragmentary. Male burials generally did not contain many grave goods, just belt elements (mostly buckles), iron knives and contents of a pouch, such as tweezers, fire strikers, etc. Among females, more richly furnished burials with bow brooches, S-shaped brooches and various types of disc brooches were observed. The archaeological dating of the site is based on the small finds, mostly female jewellery, especially the earlier mentioned brooches that are characteristic finds of the mid 6th century. The cemetery is one of those sites in the region, similarly to Ménfőcsanak and Kóny, that were established a few decades after the initial settlement of the Langobards in the region in the mid-6th century.

The site is unpublished, but an extensive report is available written by the excavator, Péter Tomka. The archaeological and skeletal material is currently stored in the Rómer Flóris Művészeti és Történeti Múzeum in Győr.

Tomka, Péter: Langobardenforschung in Nordwestungarn. In Pohl, W. & Erhart, P. (Hrsg.): *Die Langobarden. Herrschaft und Identität. Ergebnisse eines vom 2. bis 4. November 2001 in Wien abgehaltenen internationalen Symposiums.* Vienna: Verlag der Österreichischen Akademie der Wissenschaften 2005, 247–264.

Tomka, Péter: Kulturwechsel der spätantiken Bevölkerung eines Auxiliarcastells: Fallbeispiel Arrabona. In Friesinger, H. & Stuppner, A. (Hrsg.): *Zentrum und Peripherie – Gesellschaftliche Phänomene in der Frühgeschichte*. Vienna: Verlag der Österreichischen Akademie der Wissenschaften 2004, 389–409.

**S1.2. Győr - Frigyes laktanya (GYR),** Győr–Moson–Sopron County, Hungary

3rd-4th century CE

3 individuals, 3 aDNA samples

Three Roman period burials dating to the second half of the 3rd to 4th centuries CE were excavated at the Győr-Frigyes laktanya site in western Hungary in 2009. The site probably belonged to a smaller, rural settlement near Arrabona, situated along the trade route connecting Arrabona and Savaria, two important Roman period centres in the Little Hungarian Plain. Arrabona, established on the site of an earlier Celtic settlement, was one of the most significant auxiliary forts along the Danube Limes. The individuals – an adult male and two adult females – were buried in simple pit graves without any notable grave goods. Osteological analysis revealed severe and distinctive lesions on one of the individuals and identified it as the result of bone tuberculosis.

The site is archaeologically unpublished; the osteological information was published by Tamás Hajdu et al. 2012. The archaeological and skeletal material is currently stored in the Rómer Flóris Művészeti és Történeti Múzeum in Győr.

Hajdu, Tamás *et al.:* Bone tuberculosis in Roman Period Pannonia (western Hungary). *Mem. Inst. Oswaldo Cruz* 107 (2012) 1048–1053.

**S1.3. Győr - Széchenyi square (GYS),** Győr–Moson–Sopron County, Hungary

$4^{th}$-$5^{th}$ century CE

at least 80 graves, at least 80 individuals, 66 aDNA samples

In the area of present-day Győr, the eastern cemetery of the vicus (civilian settlement) presumably belonging to the Roman period auxiliary camp of Arrabona has been uncovered. The site has been known since 1949, and burials have been found in several stages since then—about 80 in total, of which 66 have been fully excavated over the past decades. The auxiliary camp of Arrabona and its associated civilian settlement were an important centre in the region during the Roman period. Although the vicus was gradually abandoned during the 4th century—likely due to the turbulent nature of the period—the civilian population probably moved into the better-protected military camp, which likely remained in use until the early 5th century.

The cemetery's dating is based on its relation to the so-called ‘dark earth’ (*schwarze Schicht, sötét réteg*), a dark stratum observed across Europe that separates Roman and medieval layers. A significant portion of the burials were dug into this ‘dark earth’, which is dated by coins from the early 5th century. Several instances of stratigraphic superposition were observed in the cemetery (partly why not all skeletons could be tied to a clear archaeological context), suggesting long-term use of the burial site.

Graves are generally very poorly furnished; the most common artefacts are simple iron buckles and double-sided combs. Only the late antique basket-shaped earring from grave 31 provides any substantial dating evidence. Stones and Roman bricks found in some of the graves reflect late Roman burial practices, but the presence of artificial cranial deformation in several cases clearly indicates usage during the mid-5th century, during the Hunnic period.

The site is unpublished, but extensive reports are available written by the excavators, Péter Tomka and Szilvia Bíró. The archaeological and skeletal material is currently stored in the Rómer Flóris Művészeti és Történeti Múzeum in Győr.

Tomka, Péter: Kulturwechsel der spätantiken Bevölkerung eines Auxiliarcastells: Fallbeispiel Arrabona. In Friesinger, H. & Stuppner, A. (Hrsg.): Zentrum und Peripherie – Gesellschaftliche Phänomene in der Frühgeschichte. Vienna: Verlag der österreichischen Akademie der Wissenschaften 2004, 389–409.

Bíró, Szilvia & Tomka, Péter: A Győr, Széchenyi téri fekete réteg rejtélyei. In Csóka úrtól Gáspár atyáig. Ünnepi kötet Csóka Gáspár OSB 75. Születésnapjára. Győr: Szent Mór Bencés Perjelség 2013, 521-538.

Bíró, Szilvia & Tomka, Péter: Über die mysteriöse ‘schwarze Schicht’ und das sog. ‘hunnenzeitliche Gräberfeld’ von Győr-Széchenyi Platz. In: *Na hranicích impéria = Extra fines imperii: Jaroslavu Tejralovi k 80. Narozeninám*. Brno: Masarykova univerzita – Archeologický ústav AV ČR, Brno 2017, 53–72.

**S1.4. Hegykő - Mező utca (HMU),** Győr–Moson–Sopron County, Hungary

late 5$^{th}$- mid 6$^{th}$ century CE

81 graves, 81 individuals, 74 aDNA samples

The cemetery at Hegykő is perhaps the most extensively researched site from the Langobard-period Pannonia. The site has been known since 1955, and excavations were carried out in several phases between 1959 and 1962. The cemetery is located near the southern shore of Lake Fertő (*Neusiedlersee*), approximately 18 km from the Roman town of Scarbantia (modern-day Sopron). The cemetery, which contains 81 graves, escaped the typical disturbance of graves—a fundamental problem of this period—making it particularly suitable for the study of chronological, cultural, and social questions. Burial customs at Hegykő differ from those observed in "classical Langobard" cemeteries: the graves are simple pit graves, considered shallow for the period, and do not feature the wooden structures interpreted elsewhere as "houses of the dead." Food offerings are found in very few cases, but there is a relatively high number of objects decorated with crosses, primarily belt buckles, which archaeological research interpreted as symbols of Christianity. Due to its distinctive characteristics, the cemetery gave its name to the so-called ‘Hegykő Group’, and was described as a community where surviving late antique groups, pre-Langobard Germanic peoples (Heruli, Suebi, etc.), and Langobards lived together. However, both this type of ethnic interpretation and the defining characteristics of the ‘Hegykő Group’ have come under criticism in recent research.

From a chronological standpoint, the site is of exceptional importance. In Transdanubia, one of the long-standing questions of archaeological research concerns the relationship between 5th- and 6th-century sites and the chronological gap between them. The polyhedral earrings with garnet inlays found in graves 3 and 23 at Hegykő are primarily known from 5th-century contexts, suggesting that, like the cemetery at Szeleste, Hegykő is one of the sites that likely began around the end of the 5th century and might represent the connection between the two horizons. This early foundation may explain the differences in burial customs observed in a community whose roots may go back to the 5th century. Based on women's jewelry, particularly the brooches, it can be assumed that the cemetery was abandoned sometime in the second half of the 6th century.

The cemetery is divided into two spatially separate grave groups, which show no demographic or socio-cultural differences. Research has primarily interpreted them as family units. Among both male and female graves, there are exceptionally rich burials. In grave 18, the remains of a woman buried in the so-called "four-brooch" dress (*Vierfibeltracht)* were found—two rosette disc fibulae fastened her outer garment, while a pair of bow fibulae were found near the pelvis. Her clothing was also adorned with an ornamented belt pendant ending in a silver-mounted decorative key and faceted pendant, and she was buried with a footed glass vessel that has parallels in northern Italy. Grave 34 at Hegykő is one of the richest male burials from the period in Pannonia. The young deceased was buried with an axe, a scale (with Roman coins used as weights), an ornate comb, and a copper bowl (so called *Perlrandbecken*) likely originating from the Mediterranean. The cemetery's finds indicate complex cultural and trade connections. In addition to the Mediterranean links mentioned in connection with the two richly furnished burials, disc and bow brooches found at the site suggest primarily Western European, Merovingian connections. Interestingly, the S-shaped brooches that are among the most common in the region are entirely absent from this site. The high number of imported items may suggest that the community played some role in the trade along the Danube, which could also explain the cemetery's overall wealth.

The site is published posthumously by the excavator István Bóna. The osteological material was first analysed and published by István Kiszely and later reevaluated by Irene Barbiera and Olga Spekker. The archaeological and skeletal material is currently stored in the Rómer Flóris Művészeti és Történeti Múzeum in Győr.

Barbiera, Irene: *Changing Lands in Changing Memories. Migration and identity during the Lombard Invasions*. Firenze 2005.

Bóna István: Das langobardenzeitliche Gräberfeld von Hegykő, Komitat Győr-Sopron. In Anreiter, P.–Bartosiewicz, L.–Jerem, E.–Meid, W. (Eds.): *Man and the animal World. Studies in Archaeozoology, Archaeology, Anthropology and Palaeolinguistics in memoriam Sándor Bökönyi.* Budapest 1998.

Bóna István – B. Horváth Jolán: Hegykő-Mező utca. In Bóna, I. & B. Horváth, J.: Langobardische Gräberfelder in West-Ungarn. Monumenta Germanorum Archaeologica Hungariae vol. 6. Budapest 2009, 31-57.

Horváth Eszter: Gemstone and glass inlaid fine metalwork from the Carpathian Basin: the Hunnic and Early Merovingian Periods. *Dissertationes Archaeologicae* 3(1) (2013) 275–302.

Kiszely István: *The Anthropology of Lombards*. BAR International Series 1979.

Koncz, István: A hegykői 6. századi temető időrendje és kapcsolatrendszere. (The chronology and cultural contacts of the 6th century cemetery at Hegykő) *Archaeologiai Értesítő* 139 (2014) 71–98.

Tomka, Péter: Problémák a Hegykő-csoport körül (The Hegykő problem). In Kovács, László & Révész, László & Bollók, Á. & Gergely, K. & Kolozsi, B. & Pető, Zs. & Szenthe, G. (eds): *Népek és kultúrák a Kárpát-medencében: Tanulmányok Mesterházy Károly tiszteletére*. Budapest: Magyar Nemzeti Múzeum 2016, 185-190.

**S1.5. Kóny – Markotai út (KMU),** Győr–Moson–Sopron County, Hungary

mid 6th century CE

20 (22?) graves, 20 individuals, 11 aDNA samples

During the rescue excavations preceding the construction of a motorway, 22 graves – two of them did not contain any human remains – dated to the 6th century came to light in 2014. The site is located close to the Roman road connecting Arrabona (today's Győr) and Scarbantia (today's Sopron) about ca. 25 kms east from the former. The cemetery is completely excavated, two of the graves were destroyed before the excavation by modern construction. All graves show the signs of contemporaneous disturbance: the traces of secondary openings were clearly observable in the soil, skeletons were generally out of position, only parts remained in situ. Due to the disturbance, the archaeological material is fragmentary. Archaeological dating of the site is based on a well-preserved pot with stamped-in decoration, shield-on-tongue belt buckles (*Schilddornschnallen*), single-sided combs and other small finds, all characteristic of the middle third of the 6th century, so the site belongs to the same horizon with the nearby Gyirmót and Ménfőcsanak cemeteries. Both the archaeological material, as well as the small number of burials suggest a short occupation of use.

The site is unpublished. The archaeological and skeletal material is currently stored in the Rómer Flóris Művészeti és Történeti Múzeum in Győr.

**S1.6. Ménfőcsanak (MFC),** Győr–Moson–Sopron County, Hungary

mid 6th century CE

25 inhumations and 3 cremations, 28 individuals, 19 aDNA samples

The Ménfőcsanak site, similarly to Gyirmót that is only ca. 1 km apart, was excavated between 1995 and 1997 as part of the rescue excavations between 1990 and 2012 connected to the building of the main road 83. The cemetery is located on al elevated ridge near the Roman road in the vicinity of the Roman town Arrabona (today's Győr). The graves were heavily disturbed, and the area was also flooded sometime after the 6th century resulting in very bad preservation of both the archaeological and osteological material. Among the grave goods placed in the burials, primarily simple, everyday items (buckles, iron knife, tweezers, beads, spindle whorl) have survived, while the more valuable items, in the case of both men and women, fell victim to disturbance. Ménfőcsanak is one of the few cemeteries – together with Kajdacs and Tamási – described as biritual in 6$^{th}$-century Pannonia. While cremation burials do appear together at these sites, their dating and thus their exact connection to the inhumation burials is dubious and requires further analysis. Similarly to Gyirmót and Kóny, the Ménfőcsanak cemetery was probably used by a rural community for a short period of time in the middle of the 6$^{th}$ century.

The site is published by the excavator Andrea Vaday. The osteological material was analysed and published by Balázs Gusztáv Mende. The archaeological and skeletal material is currently stored in the Rómer Flóris Művészeti és Történeti Múzeum in Győr.

Vaday, Andrea: The Langobard cemetery at Ménfőcsanak. *Antaeus* 33 (2015) 163–242.

Mende, Balázs Gusztáv: Brief overview of the migration period population from Ménfőcsanak. *Antaeus* 33 (2015) 243-248.

**S1.7. Szeleste (SZL),** Vas County, Hungary

late 5$^{th}$- mid 6$^{th}$ century CE

112 graves, 112 individuals, 110 aDNA samples

With 112 graves the cemetery at Szeleste, excavated in 2013, is the largest known cemetery from the first half of the 6th century in Transdanubia (present-day Western Hungary). The site is located about 20 km west of the Roman-era town of Savaria (today's Szombathely). Although by the 6th century Savaria likely lost its urban character and administrative role, as a provincial centre and due to the nearby Amber Road, it was an important regional hub during the Roman period.

In Transdanubia, researchers have long been intrigued by the chronological gap between sites associated with Germanic groups—such as the Heruli, Rugii, Suebi, and Goths—dating to the second half of the 5th century, and the newly founded cemeteries of the 6th century. The latest cemeteries belonging to the earlier group—such as those at Soponya and Balatonszemes—can be dated at most to the last decades of the 5th century. In contrast, regarding the 6th-century cemeteries (with the exception of the cemetery at Hegykő), the possibility of an origin in the 5th century has not yet been raised. This results in a shorter hiatus in Northern Transdanubia, and a longer, possibly generational gap in Southern Transdanubia, where researchers date the foundation of 6th-century cemeteries to the middle third of the century.

In Northern Transdanubia, alongside the previously mentioned Hegykő, the Szeleste cemetery may hold the key to solving this chronological gap. From the northern part of the cemetery, six individuals with artificially deformed skulls were recovered. The practice of artificial cranial deformation was widespread in the Carpathian Basin during the 5th century, but despite being well-known in neighbouring regions such as Lower Austria, it is not known from 6th-century sites in the area. Other features suggestive of a 5th-century origin include the presence of three graves lined with Roman bricks and *tegulae*, a practice not known from other 6th-century Transdanubian sites. These have parallels in 5th-century cemeteries (e.g., Mözs), and their origin can be traced to Late Roman cemeteries, such as those in nearby Savaria. However, in most cases, both the 5th-century and Szeleste examples lack the structural regularity of Late Roman brick graves. The existence of brick-lined graves and cranial deformation provides only indirect evidence for the cemetery's 5th-century beginnings. Based on the recovered finds, the cemetery clearly remianed in use at least until the middle third of the 6th century.

The most important datable grave goods in the cemetery are the fibulae. These include well-known types from the period: pairs of S-shaped fibulae of the Schwechat-Pallersdorf type, single-zoned disc brooch or pairs of rosette disc fibulae. The S-fibula type is known from 11 sites in the Carpathian Basin, making it the most common small fibula type of the era. The disc fibulae have good parallels in the Hegykő and Szentendre cemeteries, and the rosette form was widespread in both Western Europe and Transdanubia. A much rarer type is represented by a pair of bow brooches with bird-head ends (so called *Vogelkopffibel*), with the closest parallels in the eastern Carpathian Basin, specifically in the Szolnok-Szanda cemetery. This fibula type was widespread in present-day France and Germany, with the geographically closest and most accurate parallels found in present-day Czechia. Overall, based on both bow brooches and small fibula types, the site forms a transitional link between Lower Austrian and Transdanubian sites and shows clear cultural connections toward Western Europe, the Mediterranean, and Scandinavia.

The site is unpublished. The archaeological and skeletal material is currently stored in the Savaria Megyei Hatókörű Városi Múzeum in Szombathely.

Pap, Ildikó Katalin: Savaria keleti temetője és a szelestei germán temető épített és tegulás sírjai (Savaria’s eastern cemetery and the built and tegula tombs of the Szeleste Langobard cemetery). *Savaria* 38 (2016) 91–105.

Tóth, Gábor & Pap, Ildikó Katalin: Germán temető a Nyugat-Dunántúlon. In Nagy, M. & Poráčová, J. (eds): *Trendek és eredmények a biológiai kutatás és oktatás terén*. Komarno: János Selye University 2016, 22–27.

### S2. Sr, C and N isotope analysis

#### S2.1. The geological and pedological setting of Northwestern Transdanubia and the Little Hungarian Plain

In the present-day territory of Hungary, Quaternary geological development has produced thick strata in large areas, which can be classified into two main facies: hilly and flatland areas (Jámbor 2012)[1]. The region under study is one of these, with Quaternary formations predominating in the North Transdanubian region and the Little Hungarian Plain. The hilly areas, and in general most of the Transdanubian region, are characterised by loess deposits of varying thickness during the Pleistocene, which are dissected by rivers and other watercourses and the gravel-sand sediments deposited by them (Jámbor 2012; Lóczy 2015; Csillag & Sebe 2015)[1–3]. In western and northern Transdanubia, these loess layers are deposited at approximately 0-50 m thickness and represent mostly the younger Pleistocene (Jámbor 2012; Gábris & Nádor 2007)[1,4]. Their subsoil is covered by river sand and gravel layers of older Pleistocene age. Of our sites, only Szeleste is located in such a micro-region, namely the Gyöngyös Plain.

The flatland areas comprise most of Hungary's territory and can be further classified into three facies: the Great Plain, the Győr Basin and the Dráva Basin. The Quaternary sequence is much thicker in these areas, with a maximum of 450 m in the Győr basin (Gábris & Nádor 2007; Jámbor 2012; Csillag & Sebe 2015)[1,3,4]. The Danube and the Rába deposited most of the alluvium, creating extensive and thick alluvial fans during the Pleistocene and Holocene. Special mention should be made of the depressions and basins around Lake Fertő and Hanság, where a layer of lacustrine, marshy, peaty sediment was formed up to a thickness of about 20 m (Jámbor 2012)[1]. Among our sites, Hegykő is located in this area, while Kóny and Gyirmót are situated on the alluvial cone of the Rába, on the Csorna plain.

The geological and soil characteristics of the micro-regions comprising each site can be broadly characterised within this framework, but it is also worth considering them separately.

*Fertő-Basin (Hegykő)*

It is a small micro-region, bordered by the Hanság to the east, the Ikva Plain to the south, and the Fertő Hills and the Sopron Basin to the west (Dövényi 2010)[5]. A considerable part of its surface is still swampy and covered with water; the other part is a low floodplain. Before the regulation of Lake Fertő, almost the entire area was covered with water at high water levels. The soil is peaty in the marshy parts and covered with floodplain silt, boggy clay and meadow clay in the areas protected from water. Underneath are thick layers of Pleistocene sand and gravel.

*Csorna-Plain (Kóny, Gyirmót)*

The area is bordered by the Pápa-Devecser Plain to the south and east, the Kapuvár Plain to the west and the Hanság and Moson Plain to the north (Dövényi 2010)[5]. The vast majority of its surface is high and low floodplain, and the formation of the area itself can be linked to the alluvial fan of the River Rába. The soils are typically river silt, marshy and meadow clay, peat and sand. Underlying layers of sand and gravel of Pleistocene age are 50-100 m thick.

*Gyöngyös-Plain (Szeleste)*

It is a relatively large micro-region, bordered by the Rába alluvial plain in the south and east, the Vas mountain and the Kőszeg foothills in the west, and the Répce plain in the north (Dövényi 2010[5]). The surface of the area is relatively flat, with only a slight slope to the southeast, mainly covered by glacial loam and loess. Towards the south, the glacial strata are mixed with late Pleistocene gravel layers, accompanied by solifluction and cryoturbation. These gravel layers, deposited mainly by the rivers Gyöngyös, Répce and Rába, played a decisive role in the formation of the present form of the landscape.

#### S2.2. Bioavailable Sr and its relationship with the environment

Research on the mobility of the $^{87}$Sr isotope is based on the fact that it is formed in the parent rock when the radioactive isotope $^{87}$Rb decays, and then is transported by further weathering processes to the soil and groundwater, and then through plants to the whole food chain (Bentley 2006; Price et al. 2002)[6,7]. In general, the older the rock, the more the ratio of $^{87}Sr$ to $^{86}Sr$ may shift in favour of the former. Although this isotope is also radioactive, being the only one of the other Sr isotopes, its half-life is so long (4.88-1010 years) that in practice no decay can be expected, Moreover, it does not undergo significant fractionation in the food chain, so in theory for any narrow, broader region we should obtain a typical $^{87}Sr/^{86}Sr$ ratio, which is consistent with the local geological context (Bentley 2006)[8]. The reality is different, even the determination of the value of the parent rock is problematic since it matters which minerals were measured in the given piece of rock (Bentley 2006)[8]. Moreover, weathering processes are also highly dependent on many factors, so typically the bioavailable Sr value for a given region, when examined, does not fully overlap with that measured in the parent rock (Price et al. 2002; Makarewicz & Sealy 2015)[7,9]. Thus, traditionally, one of the most obvious solutions for determining the local baseline, in addition to contemporaneous human and faunal measurements, is to examine a recurrent environmental sample (Maurer et al. 2012)[10].

In our study region, very few measurements have been made on archaeological material based on the literature, but fortunately, such data are available from the site of Kóny 85 Enese (Supplementary Table S1), which is very close to our sites (Deparmentier et al. 2021)[11]. Unfortunately, the sample set is not very numerous, only one human bone, three dentin samples from animal teeth of the same age and four shells of the same age were measured. The literature indeed suggests that these are the most suitable proxies for determining locally detectable bioavailable strontium when it is not possible to take environmental samples (Price et al. 2002; Knipper 2004; Bentley 2006; Brönimann et al. 2018)[7,8,12]. The reported data have a rather wide range, from 0.71095 to 0.70975, with an average of 0,7099.

| Site | Age | Material | Type | Feature | 87Sr/86Sr | |
|---|---|---|---|---|---|---|
| Kóny 85 Enese | TLBK | human bone | Femur | 56 | 0.70984 | 0.00001 |
| Kóny 85 Enese | no data | shell | Gastropoda | 505/491 (Bag: 1125) | 0.71056 | 0.00004 |
| Kóny 85 Enese | no data | shell | Bivalvia | 547/533 (Bag: 1427) | 0.71087 | 0.00002 |
| Kóny 85 Enese | no data | shell | Bivalvia | 56/55 (Bag: 167) | 0.71095 | 0.00005 |
| Kóny 85 Enese | no data | shell | Bivalvia | 300/286 (Bag: 560) | 0.70975 | 0.00006 |
| Kóny 85 Enese | no data | animal enamel | Sus domesticus | 642/609 (Bag: 1059) | 0.70989 | 0.00002 |
| Kóny 85 Enese | no data | animal enamel | Sus domesticus | 776 / 741 (Bag: 2158) | 0.70998 | 0.00002 |
| Kóny 85 Enese | no data | animal enamel | Sus scrofa | 642 / 609 (Bag: 1059) | 0.70990 | 0.00003 |

Supplementary Table S1. $^{87}Sr/^{86}Sr$ values from the site Kóny 85 Enese

There is no other information available for Kisalföld, nor for the wider North-Western Transdanubian region. Another lesson from the micro-regional baselines experimentally established for Hungary, supported by a variety of geographic, hydrological and geological factors, is that the relationship between bioavailable strontium and the environment is far from clear (Deparmentier et al. 2021)[11]. With all this in mind, the averaged values of 0.708-0.710 (Bentley et al. 2004[6]) for Central European loess soils and 0.7096 (Giblin 2009)[13] for the Danube can be taken into account only as an indication.

#### S2.3. $^{87}Sr/^{86}Sr$ isotopic data at Kóny

From the Kóny site, we have a total of 14 measurements from 14 individuals, taken from their tooth enamel (Supplementary Dataset S1, Supplementary Fig. 1). The lowest value, 0.70909, is from a Maturus male buried in grave 445, while the highest value, 0.7131, was measured from the teeth of a male buried in grave 558. The mean of the samples is 0.71086, and the median is 0.71055. The sex ratio is relatively even, with 6 males and 5 females, in addition to two children (graves 442 and 552) and one juvenile-aged individual of indeterminate sex buried in grave 553. In the lack of extensive child datasets or bone or faunal measurements, we could only rely on the 2SD values of the whole population. There are no outliers between the values, and the data set can be considered homogeneous. There is also no significant difference between the values for men and women; both their median (women 0.71055, men 0.71099) and their mean (women 0.71103, men 0.71074) are considered similar. As the data series were found to be normally distributed by the Shapiro-Wilk test, their means are comparable by analysis of variance. The result of the ANOVA performed is p=0.9683; df=1;8; F=0.001685, i.e. there is no significant difference between the means of men and women. The difference is only in the two gender ranges, with men's values covering a slightly larger interval than women's values. At the same time, the Kolmogorov-Smirnov test run shows that there is no significant difference between their distributions (D=0.333; p=0.847).

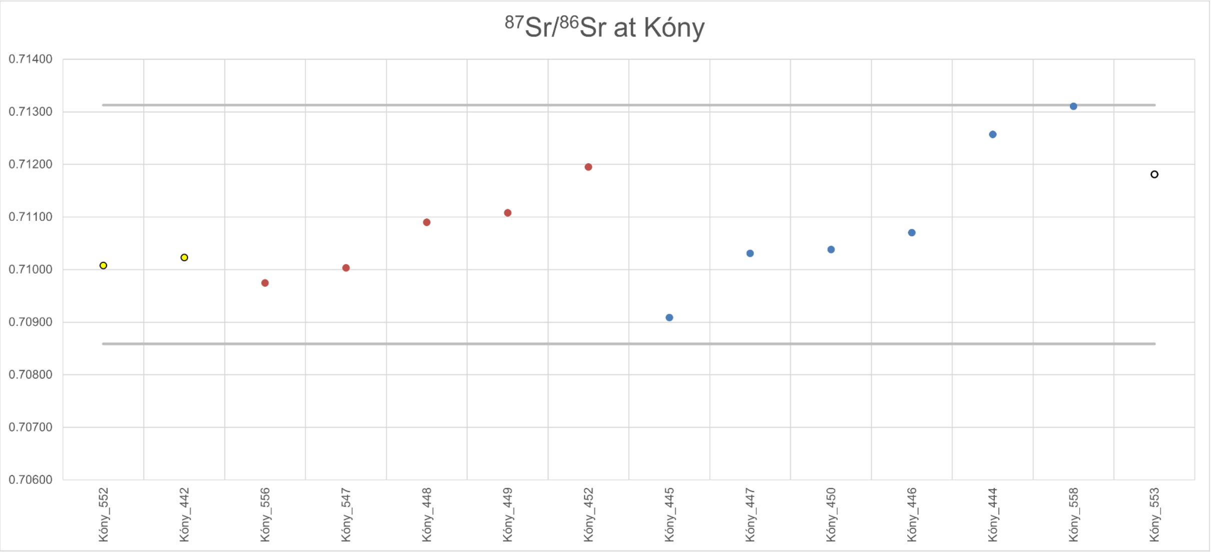

Supplementary Fig. 1. 87Sr/86Sr isotopic data at Kóny

The age distribution, apart from the under-representation of children and young adults, is quite heterogeneous, and therefore, for such a small sample size, only 1-3 individuals are included in each category. In addition, the categories with 2-3 values show very wide intervals, with individual measurements falling far apart so that they reflect the range of the population as a whole. Such categories are Adultus I, Adultus II-Maturus I, Maturus I, Maturus II-Senile. Unfortunately, the data set representing the ages does not lend itself to serious statistical comparisons, but a visual inspection of the values does not reveal any outliers.

A moderate correlation was observed for within-cemetery genetic relatedness and heavy isotopic data. The strontium values of the adult male 1 buried in grave 447 and his direct descendant, the infant child 1 found in grave 442, were found to be very similar, with a difference of only 0.00008 between the two. The other first-degree relationship showed a slightly larger difference (0.0015) between the adult male of grave 444 and the mature female of grave 449.

**S2.4. 87Sr/86Sr isotopic data at Gyirmót**

From the Gyirmót site, we have a total of 40 strontium measurements from 27 individuals (Supplementary Fig. 2). In two cases, three measurements were taken from one individual (graves 25 and 31), which, in addition to two teeth, also represent a sample from a long bone. Four individuals also have long bone data, but only one tooth was measured (graves 11, 14, 21 and 32). A further five individuals had two teeth sampled, these being graves 15, 17, 20, 27 and 35. The smallest value in the whole data set belongs to the male juvenile-age deceased in grave 15 (0.70704), while the largest value belongs to the male juvenile-age deceased buried in grave 30 (0.71623). This is the only outlier in the whole data set. The mean of the series is 0.71045, while the median is 0.70984. In the case of Gyirmót, we also have a few sub-adult measurements, so to define the local baseline, we relied on measurements from bones and their 2SD data, which interestingly matched the data for men quite well.

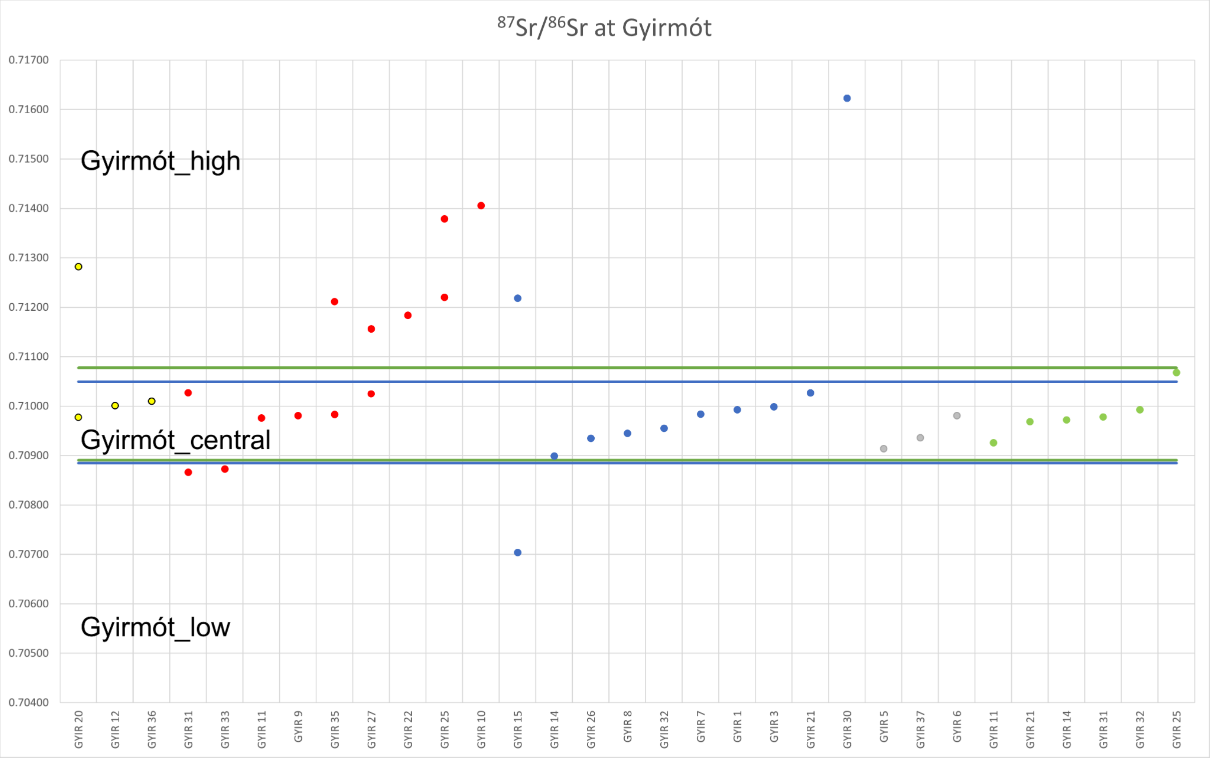

Supplementary Fig. 2. 87Sr/86Sr isotopic data at Gyirmót

The distribution of men and women is even, with nine in the former group and eleven in the latter, plus three children (20, 12, 36) and four individuals of unspecified gender (5. An interesting pattern has emerged in the breakdown of men and women, with men forming a relatively homogeneous group and, apart from outliers (15 and 30), falling into a narrower band. Women, on the other hand, show a broader range of values, with at least four distinct groups. Samples from children, bones and unidentifiable individuals all show similarities with men, so in this respect, women reflect a unique pattern. However, despite the lack of visual discrimination, an interesting result is obtained when the above groups are statistically tested, using exclusively the results coming from teeth, as, despite the former assumption, there is no significant difference between the medians (Mann-Whitney: H=5.011; p=0.286). Due to the lack of a normal distribution, the means cannot be compared, but the distributions can, and their analysis yields a surprising result, as there is no difference between males and females in this respect (Kolmogorov-Smirnov: D=0.4125; p=0.1051). Another interesting addition is that the values for males, individuals of indeterminate sex, coincided very well with the results from the bone samples, together allowing the definition of a very marked and well-defined local baseline.

Also, when examining the age distribution, only samples from teeth were included to ensure that other biasing effects did not affect the results. The largest groups were Adultus I and Adultus II, with 8 individuals in the former and 7 in the latter. The distribution of the samples was not at all similar, however, with the former group occupying a very narrow zone on the scale (between 0.70866 and 0.7127) and the latter a relatively wide zone (between 0.70919 and 0.71378). The other age groups were broadly similar, falling into one of these two distributions, but with no more than 2-4 individuals. The Adultus-Senilis group was similar to Adultus I, and Adultus I-II, Juvenis, Maturus I, and Infans I-II were similar to Adultus II.

It is also worth mentioning individuals with double or even triple measurements. The first molar (36) of the Infans I-II individual buried in grave 20 showed a significant difference from the value measured in the second molar (37), and the second value falls in the middle of the zone defined as the local baseline. This suggests that in later life he may have lived only in the narrower neighbourhood and been reared elsewhere in his younger childhood. Another possibility is that we see the mother's ratio reflected in this measurement. Among the females, there were three double measurements, the first being grave 31, with both teeth (16 and 18 molars) within the local zone but at the two extreme ends of the zone. The woman buried in grave 27 reflects a different situation. In her earlier years, this individual may also have spent more time away from this region before living and being buried here.

Unfortunately, it is difficult to reconcile the strontium isotope results with the genetic results, as apart from a third-degree linkage, no first-degree links and hence pedigrees within the community could be reconstructed. Moreover, of the Infans I child and the Juvenis buried in grave 36, only the former yielded results, so we cannot comment on the relationship between the two from an isotopic perspective

**S2.5. 87Sr/86Sr isotopic data at Szeleste**

From the site of Szeleste, we have a total of 96 Sr measurements, of which 90 were taken on human teeth and 6 from archaeological animal bones (Supplementary Fig. 3). Only one tooth from each human skeleton was measured, except for the individual Infans I from grave 550, whose teeth 16 and 75 were also measured, but the difference between the two measurements was negligible (0.71121 and 0.71116). The mean of the whole data set is 0.71098, the median is 0.71117, the minimum value is 0.70796 (Adultus male from grave 534), and the maximum is 0.71456 (Maturus female from grave 434). The whole data set covers a wide range, and the standard deviation is small (0.001285), yet its distribution is homogeneous and continuous, with perhaps the only measurement from grave 434 being minimally different from the others. In the case of Szeleste, we could rely not only on the abundance of children but also on faunal data to define the local baseline. The 2SD boundaries for the children defined a narrower middle range, while the 2SD boundaries for the animals outlined a wider range.

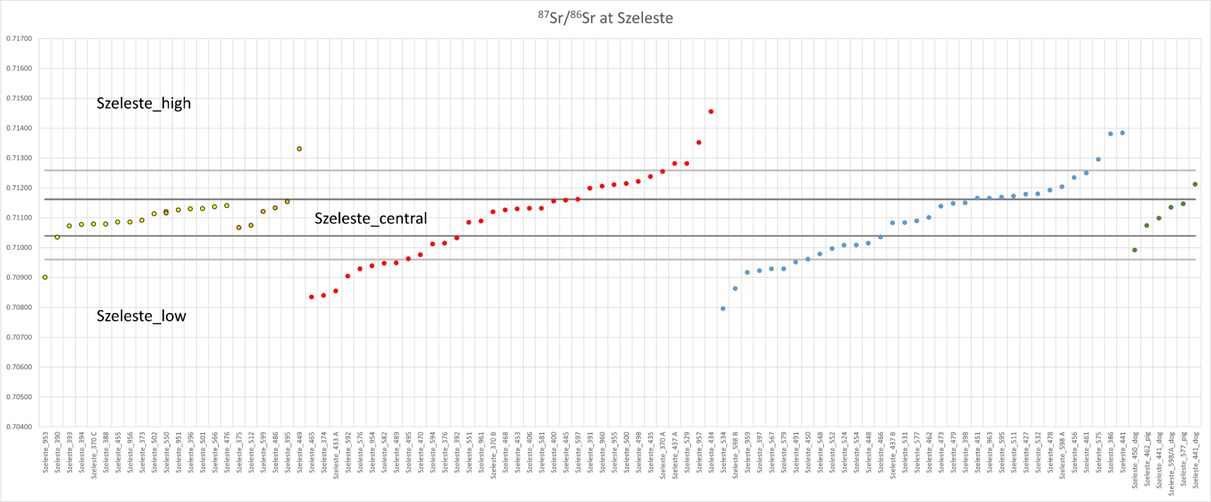

Supplementary Fig. 3. 87Sr/86Sr isotopic data at Szeleste

The sex distribution is evenly balanced, with the same number of adult males for every 34 adult females, 16 measurements of 15 Infans I individuals and 6 Infans II children, and a separate group of six animal data to compare directly with these groups. The sub-groups are not characterised by outlier values either; the only one that stands out in a negative sense from the Infans I group is the measurement of the child buried in grave 953, who is too remote from the other individuals in the group. Excluding this measurement from the analysis, the Infans I group already followed a normal distribution, so there was no obstacle to performing the ANOVA. However, as Levene's test for homogeneity of variances (p=0.0001575) was not satisfied, a Welch F test was used (F=0.3658; df=18.89; p=0.8299), which indicated that there was no significant difference between the group means. Pairwise Tukey's tests were performed to look for additional possible between-group differences, but were unsuccessful, with significant agreement found in all subgroup pairwise comparisons.

When comparing the different age groups, we found a diverse mix of groups. The group with the largest number of items is represented by Maturus individuals, with 34. Second in line is the Adultus group, with 24 individuals, while third place goes to the Infans I group, with the 16 children already mentioned. The Infans II group is made up of three subgroups, and in addition to the three children classified as Infans II, it also includes two Infans I-II and one Infans II-Juvenile. Other categories include Juvenis (4), Adultus-Maturus (3) and Maturus-Senilis (3). No outlier was found in any group except the Infans I category mentioned above, so it was considered worthwhile to compare the Adultus and Maturus categories statistically, as there was a noticeable difference in the medians and distributions in the plot. However, although Maturus individuals appeared to have a higher Sr rate, the tests did not confirm our hypothesis. The F-test showed that the variances of the two groups did not differ significantly (F=1.1929; p=0.667), and the t-test confirmed that their means did not differ significantly (t=1.6955; p=0.09554), thus indicating that there is no difference between the two groups.

In the case of Szeleste, genetic testing has revealed a first-degree relationship in several cases, although in most cases, no extensive relationships have been identified. For the most part, only nuclear families have been reconstructed, so that the correlation between the Sr rate and genetic relatedness can only be investigated partially. One of the largest family trees includes the head of the family buried in grave 962, his two sons (graves 473 and 479), and their niece (474) from an unknown sibling. Unfortunately, of these individuals, measurements are only from the two brothers, but the results confirm their close descent, with a difference of only 0.00009 between the two measurements. Moreover, both of them are located within the local range defined by Infans I, in the upper half of it. A similar situation was observed for graves 451 and 595, where the two were also brothers, and their measurements were also very close to each other, with a difference of only 0.00003. These two measurements were, albeit slightly, outside the locally defined zone, but still within the wider local range defined by the animals. Another nuclear family is formed by the individuals buried in graves 552, 954 and 393, of which only the father (552) and mother (954) have Sr measurements. The difference between the two results is 0.00058, which is still not a large difference in the overall range of the dataset, and both individuals are outside the narrower local range defined by Infans. The last kinship relationship worth examining in terms of Sr isotope was between the males of grave 427 (father) and grave 398 (son). Again, the difference between the two measurements is relatively small, 0.00029, but while technically the father is outside the local range of Infans I, the son is located inside.

#### S2.6. $^{87}Sr/^{86}Sr$ isotopic data at Hegykő

At Hegykő, we have a total of 88 strontium isotope measurements from 69 individuals (Supplementary Fig. 4). Of the individuals with multiple samples, we also have data from 9 cases of bone inorganic material, so apart from children, we have more solid data on the local baseline. These individuals represent roughly equal proportions of children (graves 5, 74), females (graves 21, 28, 53, 54) and males (graves 20, 42, 69). For the majority of these individuals, we have data from only one tooth in addition to bone, but for the female in grave 21, we have three measurements altogether from the left femur and two from different teeth. In total, we have ten individuals in our data set for whom we have mobility data from two different teeth. In addition to 2SD data from Infans I children, 2SD data from Infans II individuals were also used to define local baselines, but the narrowest zone was still the 2SD data from measurements of the inorganic bone material.

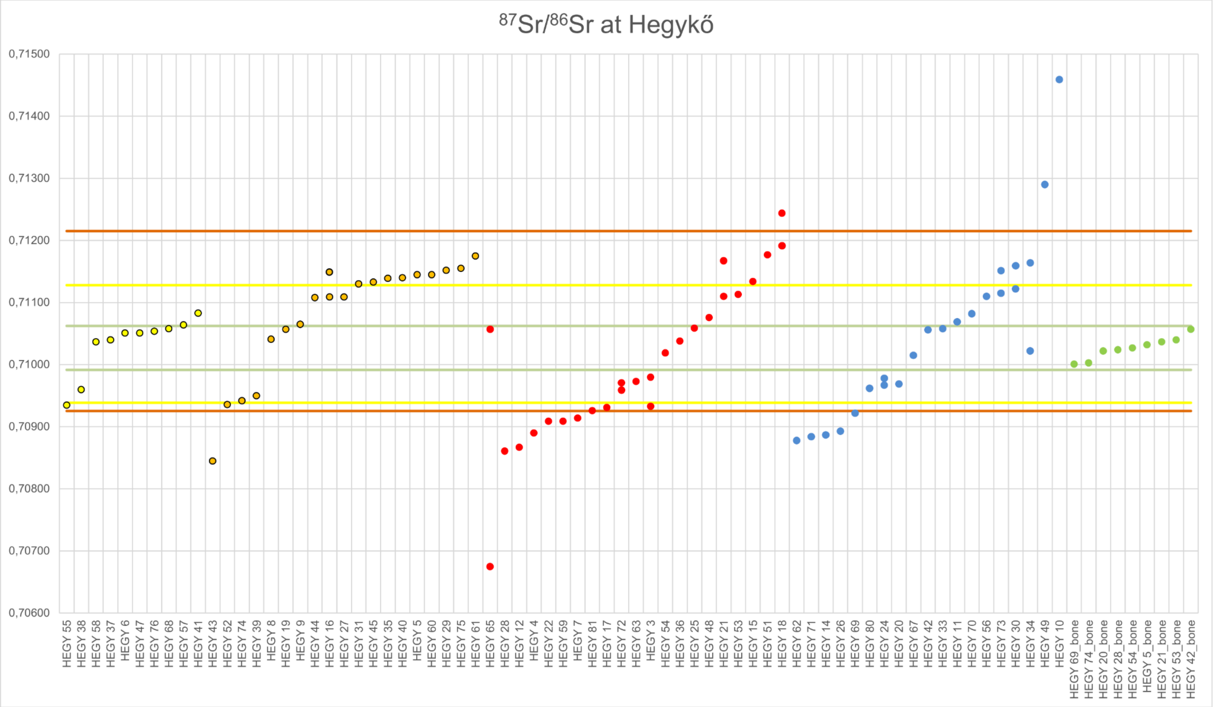

Supplementary Fig 4. 87Sr/86Sr isotopic data at Hegykő

The maximum value of the whole data set is 0.71459 (10th grave Maturus male), while the minimum is 0.70675 (65th grave Maturus female), both of them can be considered outliers, but the other measurements rather form a homogeneous array. The mean is 0.71039, the median is 0.71046, and the standard deviation is 0.00112. If only the sampled teeth are considered, the ratio of males to females seems balanced, with 23 males for every 26 females, but the proportion of children is markedly high, with a total of 30 infants I (n=10) and II (n=20). The individual categories do not show a normal distribution, so they were compared with each other using Kruskall-Wallis and paired Mann-Whitney tests. The result of the former test showed no significant differences between males, females, Infans I, Infans II and bone-derived values (H=7.687; p=0.1037), whereas the latter showed moderate differences between the Infans II category and the other groups, which disappeared after sequential Bonferroni correction.

In terms of ages, the picture is very varied, with many anthropological age categories represented, but in fact, most categories only include one or two individuals. Four of them can be highlighted and statistically compared: the Adult I-II category, where we have 12 measurements, the Infans II group, where we have 20 measurements, the Infans I group, where we have 10 results, and the Maturus I-II category, with 7 results. In the statistical comparison, the outcome was similar to the one mentioned above, with a significant difference between the age categories based on the Kruskal-Wallis test (H=7.879; p=0.04854), and the Mann-Whitney paired tests also showed that the main reason for this was due to the elements of the Infans II group, with significantly higher strontium ratios than the other three groups. In synchrony with the above, these differences disappeared again by applying the sequential Bonferroni correction. The values for the other age categories, with one or two exceptions, are scattered within the very wide Adultus I-II range, so there is no age pattern or trend is reflected in the Sr values.

Thanks to the numerous pedigrees, we also have the opportunity to compare genetic and isotopic data in more detail. At the same time, one of the pedigrees, in which we find many nuclear families linked by many third-degree relationships, represented only a few sampled individuals, all of whom were associated with the Infans II age. In the case of graves 27 and 29, there are two female siblings, and there was not even a large difference (0.0004) between their values, but they are separated by the local baseline defined by the 2SD of the Infans I values. The 8-9 year old girl buried in grave 27 can thus still be considered local, while her sister buried in grave 29 has a value outside the narrow local zone. The other elements of the same pedigree, the individuals buried in graves 31, 35 and 40, are only loosely related, but it is remarkable how close their values are to each other compared to the two sisters mentioned above, so that they all occupy a narrow range of values outside the local boundary. Unfortunately, we also have only sporadic data on another extensive pedigree: in addition to the grandmother buried in grave 61, we have also measured the grandchildren buried in graves 60 and 75, who are also cousins. The values of all three of them were within the aforementioned range, a little above the locality threshold.

The children buried in graves 5 and 16 were part of a small nuclear family as siblings, and we have double measurements of both of them. The bone measurement of the former child represented the centre of the narrow local zone defined by the bones, but the sample from tooth 46 was already in the positive band outside the local zone outlined above. Both values from grave 16 were located in the same band, also, around the upper border of the local baseline. The siblings buried in graves 8 and 9 also represented a fragment of an elementary family, but their Sr isotope values were within the narrow local range defined by the human bones, in relative proximity to each other. In relation to parent-child relationships, four family tree fragments also carry information, so we have important data on this as well. The difference between the Maturus age male found in tomb 11 and his son found in tomb 45 is 0.00064, which means that the father is still within the narrow local range, while the son is outside the local range of Infans I. The distance between the strontium values of the mother in tomb 41 and the son in tomb 24 is larger, 0.0009, yet both are within the local range of Infans I. The mother-son pair of graves 81 and 74 is a more interesting case, although their Sr values are very close, they both fall outside the Infans I local range, but in a more negative direction. The last parent-child pair is the pair of graves 51 and 57, with Sr values of 0.00113, i.e. the largest difference. The child is on the border of the narrow local range defined by the bones, but the mother is far beyond the local range of the Infans I's, in the more positive range.

Among the double-measured individuals, the largest difference was found for the woman buried in grave 65, with her tooth 16 value of 0.70675, while her tooth 38 value was 0.71057. This means that she was probably later in the area of the future cemetery, since her tooth 38 is located in the local band defined by the bones, while her earlier years were spent at a considerable distance, since her tooth 16 can be interpreted as an outlier for the whole community. The second largest distance between two teeth from the same individual was from Juvenis associated with grave 34, interestingly, the situation here was similar to the previous one. The 36th tooth suggests that he was not raised locally, but may have spent his later life in the local area, at least as tooth 18 suggests. The value of the individual tooth of Maturus II buried in grave 28 was also very far from the local values, suggesting that he was not locally raised, but the sample from his bone confirmed the narrower local range. A similar situation is shown by individuals 74, 5, 21, 53, all of whom had tooth values in a range outside the narrow local range, but had their bone values in a very narrow local range.

#### S2.7. Comparative $\delta^{13}C$ and $\delta^{15}N$ data at Győr, Szeleste, Gyirmót, Hegykő and Kóny

$\delta^{13}C$ and $\delta^{15}N$ isotopic ratios are commonly used in archaeology to study ancient diets. Thanks to the fractionation of these isotopes, an insight can be acquired about the basic photosynthetic pathway of the food web of the ancient organism, and its position in it (Katzenberg 2008; Lee-Thorp 2008)[14,15]. Millet (Panicum miliaceum), domesticated in East Asia in the mid-6th millennium, had already become an intensively cultivated plant in Europe by the beginning of the 1st millennium, but its $C_4$ photosynthetic way differs from that of $C_3$ photosynthetic way of temperate cereals (Ventresca Miller & Makarewicz 2019; Martin et al. 2021)[16,17] In this sense, $\delta^{13}C$ is a good indicator of the different crop diets, with a more positive value (>-18‰) reflecting a higher rate of direct or indirect millet intake. Generally, marine-derived diets have the same directional effect; however, freshwater diets can be reflected in variable $\delta^{13}C$ values, sometimes resulting in more negative values (<-21‰) than herbivores. $\delta^{15}N$ is a predictor of the position of the former organism in the food web; generally speaking, an increase of +2 - 6‰ per trophic level can be expected (Hedges & Reynard 2007; Lee-Thorp 2008; Makarewicz & Sealy 2015)[9,15,18].

When analysing the diet data, we first wanted to know whether the aggregated 6th century data, i.e. the population data of Szeleste (n=105), Hegykő (n=75), Gyirmót (n=35) and Kóny (n=19), together show any deviation from the late Roman data of Győr (n=60). Only measurements that met the commonly used collagen quality criteria in the literature (Ambrose 1991, van Klinken 1999)[19,20] were considered. First, comparing $\delta^{13}C$ values, we were immediately confronted with the lack of a normal distribution of the data series, so we used the Mann-Whitney test. The result was a significant difference (U=3811; p<0.0001), i.e. overall, we find significantly higher $\delta^{13}C$ values in the late antique period than in the 6th century (Supplementary Fig. 5), which is most likely due to the different rates of millet consumption. It is worth noting, however, that we have a large number of outliers in the 6th century, which are in a range that suggests even more pronounced millet consumption than the values for Győr.

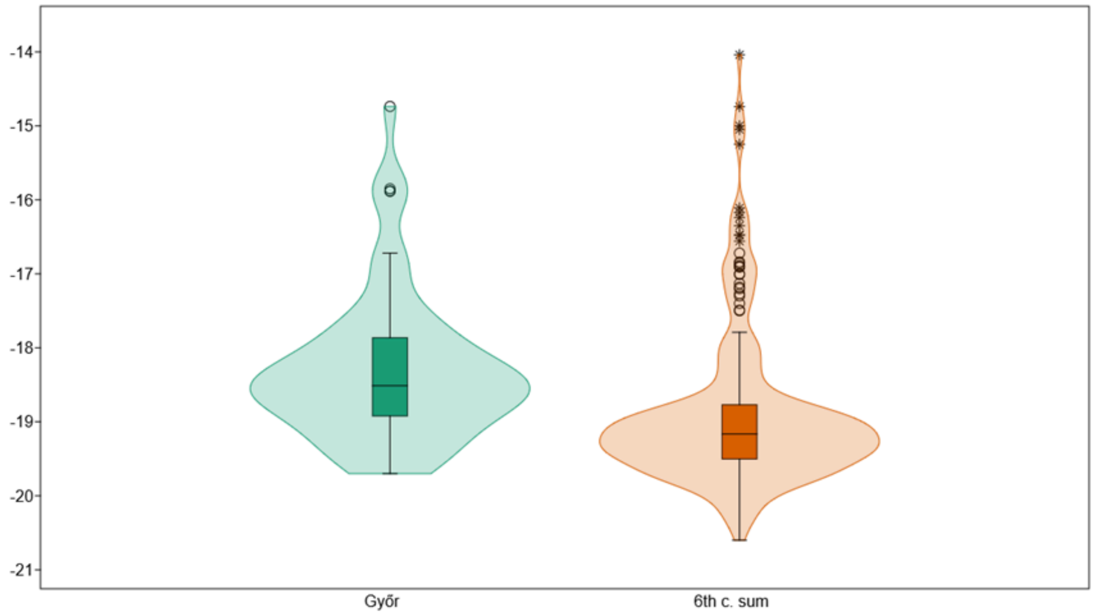

Supplementary Fig. 5 Comparison of δ$^{13}$C values between 4-5th century site and 6th century sites

There is also a characteristic difference in δ$^{15}$N values (Supplementary Fig. 6), but we did not find as many outliers as for δ13C values, i.e. our data set turned out to be much more homogeneous. Due to the lack of a normal distribution, we again resorted to the Mann-Whitney test ($U=2148$; $p<0.0001$), and the result confirmed our expectations, with significantly higher protein intakes among individuals from the Győr site.

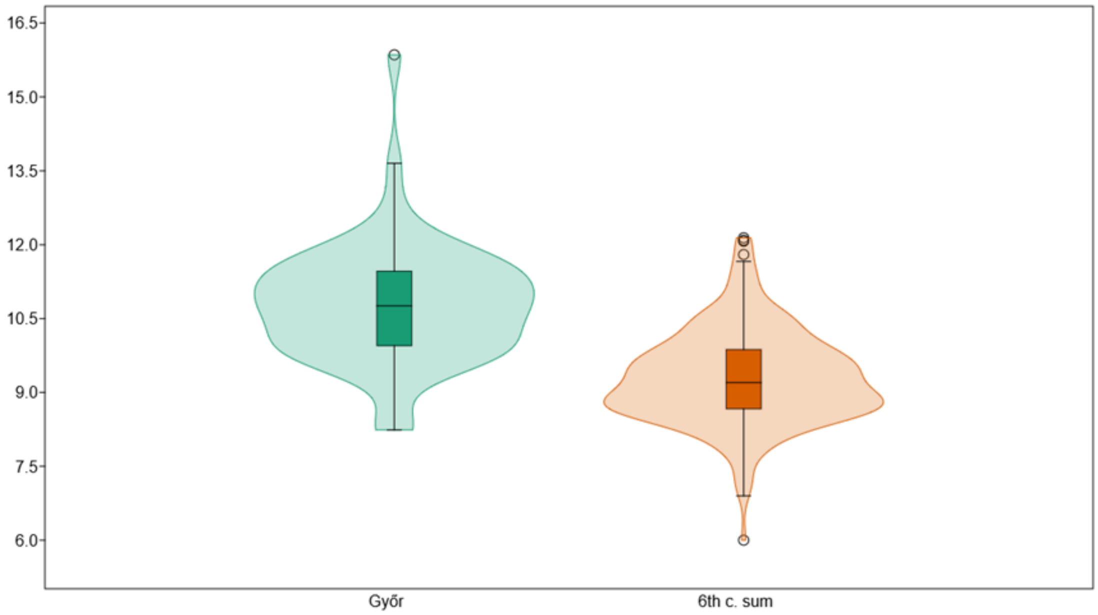

Supplementary Fig. 6. Comparison of δ$^{15}$N values between 4-5th century site and 6th century sites

We looked for further reasons for the significant difference by using site-by-site disaggregation, and first looked at the δ$^{13}$C values (Supplementary Table S2, Supplementary Fig. 7). Mainly outliers are found in the populations of Hegykő and Szeleste, and these are in a more positive range than the Győr values, yet the median values of Gyirmót and Kóny, which show fewer such outliers, seem to fall much further away from the Győr median. In the absence of a normal distribution, we used paired Mann-Whitney tests and sequential Bonferroni correction, and the results confirmed our observations. The difference between Győr and Hegykő is no longer significant, and the difference between Szeleste and Kóny is negligible. Gyirmót is also related to Kóny and Szeleste, but in all other combinations, there is a significant difference between the sites.

| Mann-Whitney pairwise p=0.05 (red-after sequential Bonferroni significance) | Győr | Gyirmót | Hegykő | Kóny | Szeleste |
|---|---|---|---|---|---|
| Győr | | 3.652E-05 | 0.02548 | 2.356E-06 | 2.707E-09 |

| | | | | | |
|---|---|---|---|---|---|
| Gyirmót | 3.652E-05 | | 0.0002664 | 0.1106 | 0.836 |
| Hegykő | 0.02548 | 0.0002664 | | 1.809E-06 | 4.737E-08 |
| Kóny | 2.356E-06 | 0.1106 | 1.809E-06 | | 0.04209 |
| Szeleste | 2.707E-09 | 0.836 | 4.737E-08 | 0.04209 | |

Supplementary Table S2. Paired Mann-Whitney tests on $\delta^{13}C$ values between 5 sites

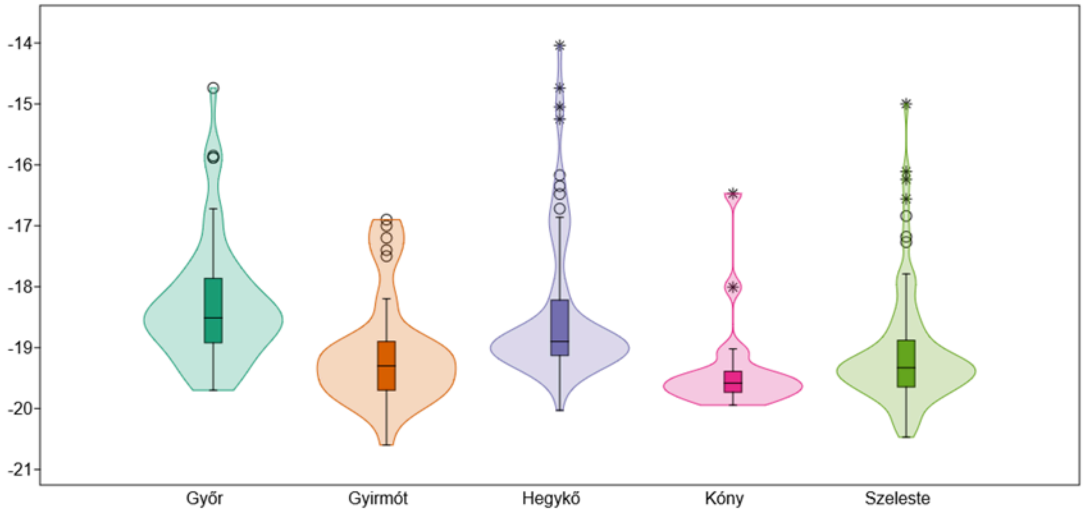

Supplementary Fig. 7. Comparison of $\delta^{13}C$ values across five sites

$\delta^{15}N$ values were subjected to a similar analysis, and here the picture is somewhat consistent, with all sites except Kóny in a much more negative range (Supplementary Table S3, Supplementary Fig. 8). It is noteworthy that in this case, too, Hegykő has a more significant amount of outliers, but here there is a clear difference between the average values for Győr and Hegykő. Together with Gyirmott and Szeleste, they are clearly in a different band. The Mann-Whitney paired tests carried out show clear evidence of similarity between Szeleste and Gyirmót, and after the sequential Bonferroni correction, the difference between Hegykő and Gyirmót is no longer significant. However, the combination of all other sites and their differences remained significant, and Győr and Kóny are no exception.

| Mann-Whitney pairwise p=0.05 (red-after sequential Bonferroni significance) | Győr | Gyirmót | Hegykő | Kóny | Szeleste |
|---|---|---|---|---|---|
| Győr | | 1.641E-09 | 1.403E-14 | 0.005602 | 9.056E-13 |
| Gyirmót | 1.641E-09 | | 0.02863 | 0.0003826 | 0.5206 |
| Hegykő | 1.403E-14 | 0.02863 | | 2.284E-06 | 0.0001492 |
| Kóny | 0.005602 | 0.0003826 | 2.284E-06 | | 0.000595 |
| Szeleste | 9.056E-13 | 0.5206 | 0.0001492 | 0.000595 | |

Supplementary Table S3. Paired Mann-Whitney tests on $\delta^{15}N$ values between 5 sites.

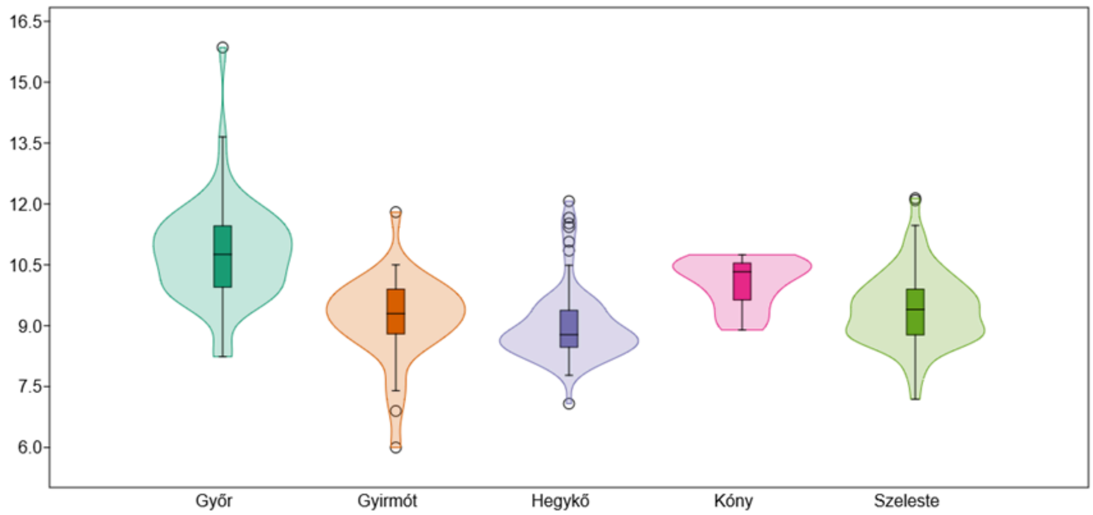

Supplementary Fig. 8. Comparison of $\delta^{15}N$ values across five sites

Further breaking down our dataset into adult males and females without sub-adults (Supplementary Table S4, Supplementary Fig. 9), but this time sticking only to the 6th century sites, we sought to answer the question of whether the different behaviour of Hegykő and Kóny is sex-specific. In addition, at this level, we felt justified in including the reported data from

Szólád (n=25), since many similarities can be found between Szólád and Hegykő (Alt et al. 2014)[21]. Apparently, there is no difference in the case of Hegykő concerning $\delta^{13}C$, with the quartiles shown in the boxplots occupying very similar bands, while in the case of Kóny, some divergence is noticeable. Only a moderate number of outliers are observed, with most cases coming from the group of women and men found in Szeleste; in this respect, Szólád shows even more unique cases for both sexes. The data isolated at each site show no gender differentiation, with perhaps only Kóny showing some variation between males and females.

| Mann-Whitney pairwise p=0.05 (red-after sequential Bonferroni significance) | Hegykő-F | Hegykő-M | Szólád-F | Szólád-M | Szeleste-F | Szeleste-M | Kóny-F | Kóny-M | Gyirmót-F | Gyirmót-M |
|---|---|---|---|---|---|---|---|---|---|---|
| Hegykő-F | | 0.9436 | 0.0185 | 0.002005 | 0.0001 | 0.001367 | 0.04633 | 0.0009911 | 0.04712 | 0.01196 |
| Hegykő-M | 0.9436 | | 0.02127 | 0.003707 | 0.0001627 | 0.00155 | 0.04506 | 0.001398 | 0.0614 | 0.01357 |
| Szólád-F | 0.0185 | 0.02127 | | 0.8483 | 0.1736 | 0.571 | 0.4299 | 0.00742 | 0.9752 | 0.5926 |
| Szólád-M | 0.002005 | 0.003707 | 0.8483 | | 0.1773 | 0.702 | 0.4269 | 0.009701 | 0.7896 | 0.9173 |
| Szeleste-F | 0.0001 | 0.0001627 | 0.1736 | 0.1773 | | 0.409 | 0.969 | 0.06572 | 0.2805 | 0.4472 |
| Szeleste-M | 0.001367 | 0.00155 | 0.571 | 0.702 | 0.409 | | 0.6464 | 0.04992 | 0.7308 | 0.8695 |
| Kóny-F | 0.04633 | 0.04506 | 0.4299 | 0.4269 | 0.969 | 0.6464 | | 0.1207 | 0.6671 | 0.6094 |
| Kóny-M | 0.0009911 | 0.001398 | 0.00742 | 0.009701 | 0.06572 | 0.04992 | 0.1207 | | 0.05694 | 0.06269 |

| | | | | | | | | | | |
|---|---|---|---|---|---|---|---|---|---|---|
| Gyirmót-F | 0.04712 | 0.0614 | 0.9752 | 0.7896 | 0.2805 | 0.7308 | 0.6671 | 0.05694 | | 0.9529 |
| Gyirmót-M | 0.01196 | 0.01357 | 0.5926 | 0.9173 | 0.4472 | 0.8695 | 0.6094 | 0.06269 | 0.9529 | |

Supplementary Table S4. Paired Mann-Whitney tests conducted on $\delta^{13}C$ values among five sites. focusing on adult males and females

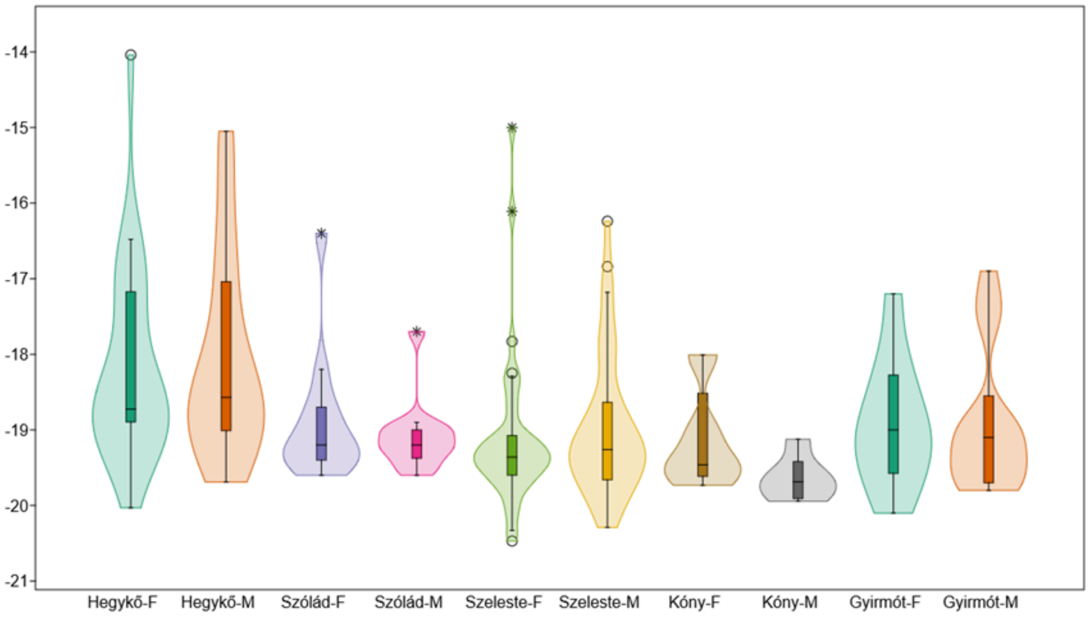

Supplementary Fig. 9. Comparison of $\delta^{13}C$ values across five sites, focusing on adult males and females

In the statistical analysis, most of the categories showed a non-normal distribution, so again, paired Mann-Whitney tests were used. The results faithfully reflect the observations on the boxplot, with the Hegykő site different from all other sites. However, after performing a sequential Bonferroni correction, most of these significant differences disappear, remaining only for the female data associated with Szeleste. A similar phenomenon is observed for the male data associated with Kóny, where we see differences for several other sites, but after the correction, a significant difference remains for the female data from Hegykő.

The breakdown of $\delta^{15}N$ values by sex revealed exciting differences (Supplementary Table S5, Supplementary Fig. 10). The differences between the sexes at individual sites seem to be much more marked in this respect, with Hegykő and Szólád in particular being good examples. In the case of Hegykő, the significant number of outliers with high values seems to be largely associated with men, so it is not surprising that they disappear in the male-female breakdown. Surprisingly, the distribution of each category followed a normal distribution in the vast majority of cases, and in the remaining two cases was just below the significance threshold, so instead of a Mann-Whitney test we tried ANOVA, which revealed a significant difference (df=181 [9,172]; F=5,338; p<0,0001) with equal variance (Levene's test: p=0,1246).

| Tukey's Q pairwise p=0.05 | Hegykő-F | Hegykő-M | Szólád-F | Szólád-M | Szeleste-F | Szeleste-M | Kóny-F | Kóny-M | Gyirmót-F | Gyirmót-M |
|---|---|---|---|---|---|---|---|---|---|---|
| Hegykő-F | | 0.0435 | 0.07774 | 1.528E-05 | 0.0207 | 0.002522 | 0.03201 | 3.344E-05 | 0.8707 | 0.07071 |
| Hegykő-M | | | 1 | 0.312 | 1 | 1 | 0.9454 | 0.1093 | 0.9894 | 1 |
| Szólád-F | | | | 0.5574 | 1 | 1 | 0.9795 | 0.2165 | 0.9832 | 1 |
| Szólád-M | | | | | 0.1183 | 0.2755 | 1 | 0.995 | 0.08415 | 0.5092 |
| Szeleste-F | | | | | | 0.9998 | 0.874 | 0.04531 | 0.995 | 1 |
| Szeleste-M | | | | | | | 0.961 | 0.1005 | 0.944 | 1 |
| Kóny-F | | | | | | | | 0.9794 | 0.6356 | 0.9748 |
| Kóny-M | | | | | | | | | 0.02737 | 0.1919 |
| Gyirmót-F | | | | | | | | | | 0.9845 |
| Gyirmót-M | | | | | | | | | | |

Supplementary Table S5. Paired Mann-Whitney tests conducted on $\delta^{15}N$ values among five sites. focusing on adult males and females

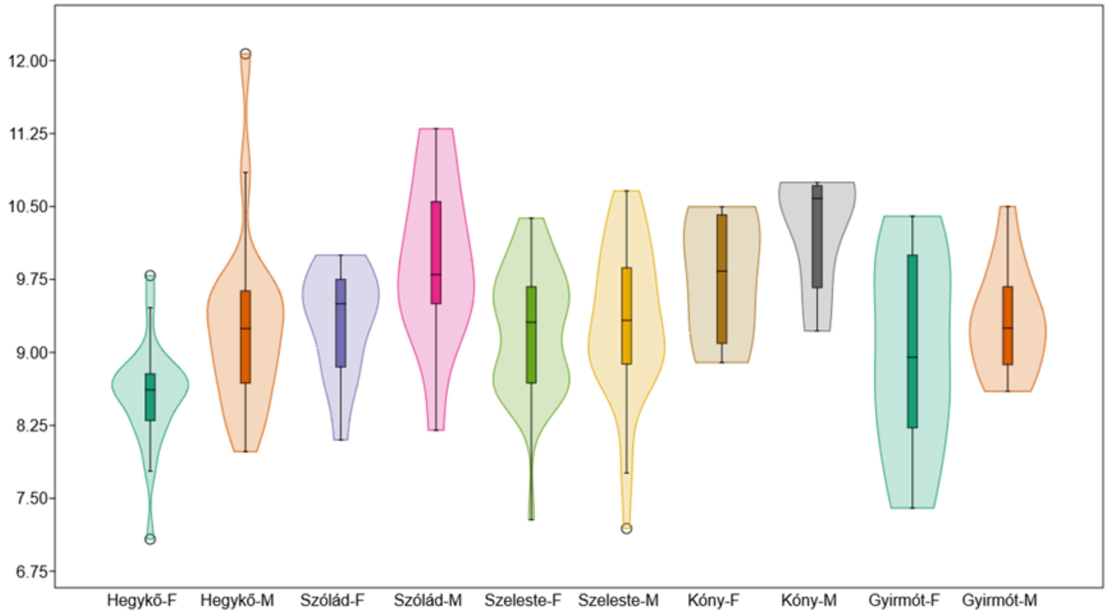

Supplementary Fig. 10. Comparison of $\delta^{15}N$ values across five sites, focusing on adult males and females

Thanks to the pairwise Tukey's tests, it is also clear that the results of the women associated with Hegykő cause the greatest fluctuation in the data set, with all categories except those associated with Gyirmót differing significantly or very close to the significance level.

One more aspect we were curious about was the result of comparing dietary data across sites, and this concerned the dietary habits of men buried with weapons. For this purpose, we collected separately the armed men of Szólád (n=11), Hegykő (n=11) and Szeleste (n=18) and compared them with the non-armed adult population of the same sites. In terms of $\delta^{13}C$ data, the separateness of the unarmed individuals from Hegykő is immediately striking, as well as the extremely wide range of their data (Supplementary Table S6, Supplementary Fig. 11). The difference from the individuals with weapons from the same sites is remarkable, but not observed in Szólád and Szeleste, where the diet is more consistent. In the absence of normal distributions, Mann-Whitney pairwise tests were performed and our suspicion was confirmed: the difference in the non-armed individuals in the Hegykő is significant, and it is significant for almost all other subgroups.

| Mann-Whitney pairwise p=0.05 (red-after sequential Bonferroni significance) | Szólád-weapons | Szólád-no weapons | Hegykő-weapons | Hegykő-no weapons | Szeleste-weapons | Szeleste-no weapons |
|---|---|---|---|---|---|---|
| Szólád-weapons | | 0,9058 | 0,06559 | 0,000836 | 0,3448 | 0,4877 |

| | | | | | | |
|---|---|---|---|---|---|---|
| Szólád-no weapons | 0,9058 | | 0,1093 | 0,0005762 | 0,3216 | 0,328 |
| Hegykő-weapons | 0,06559 | 0,1093 | | 0,07024 | 0,01614 | 0,04464 |
| Hegykő-no weapons | 0,000836 | 0,0005762 | 0,07024 | | 8,564E-05 | 7,58E-06 |
| Szeleste-weapons | 0,3448 | 0,3216 | 0,01614 | 8,564E-05 | | 0,8122 |
| Szeleste-no weapons | 0,4877 | 0,328 | 0,04464 | 7,58E-06 | 0,8122 | |

Supplementary Table S6. Paired Mann-Whitney tests conducted on $\delta^{13}C$ values among three sites, focusing on males with weapons and other adults without weapons.

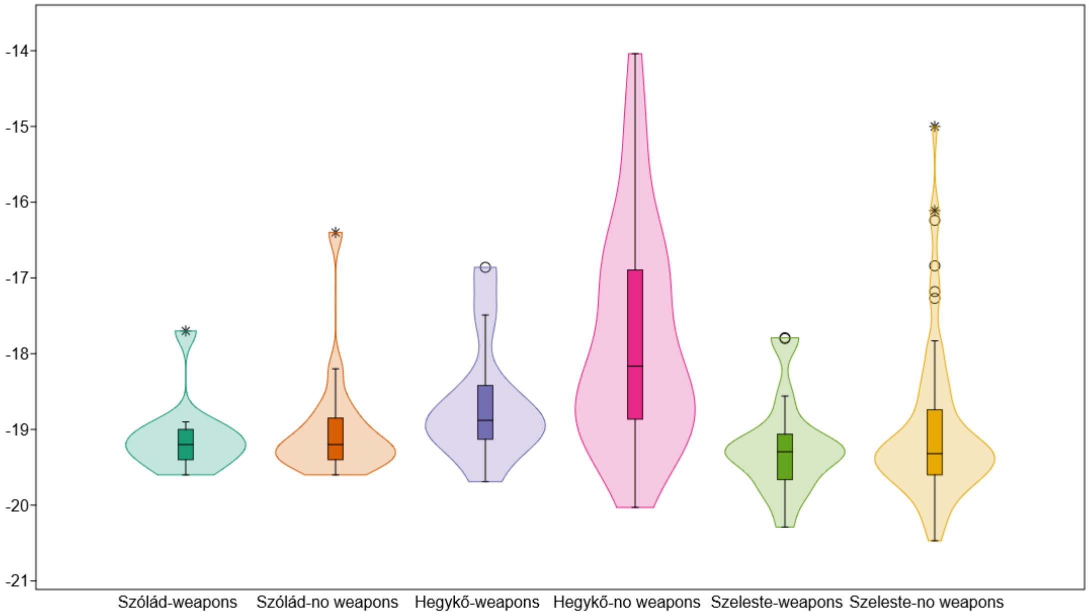

Supplementary Fig. 11. Comparison of $\delta^{13}C$ values across three sites, focusing on males with weapons and other adults without weapons.

There is also much greater fluctuation in $\delta^{15}N$ values within sites, and apparently higher protein intakes were associated with the armed male group (Supplementary Table S7, Supplementary Fig. 12). In this respect, the population of Hegykő seems to be in the lead, and this phenomenon is moderately observed for Szeleste. A positive result was obtained when testing for a normal distribution, so there was no obstacle to running the ANOVA. The key lesson is that the unarmed population of Hegykő is significantly different from almost all other categories and sites, including the armed population of Hegykő. The other interesting phenomenon is that no significant difference is found among armed males from any site. The third important observation is that in Seleste this differentiation does not reach a significant level between the armed and unarmed population.

| Tukey's Q pairwise p=0.05 | Szólád-weapons | Szólád-no weapons | Hegykő-weapons | Hegykő-no weapons | Szeleste-weapons | Szeleste-no weapons |
|---|---|---|---|---|---|---|
| Szólád-weapons | | 0,04374 | 0,7297 | 2,384E-07 | 0,5012 | 0,001409 |
| Szólád-no weapons | | | 0,727 | 0,01849 | 0,7386 | 0,9902 |
| Hegykő-weapons | | | | 0,000413 | 1 | 0,2576 |
| Hegykő-no weapons | | | | | 4,849E-05 | 0,004596 |
| Szeleste-weapons | | | | | | 0,1713 |
| Szeleste-no weapons | | | | | | |

Supplementary Table S7. Paired Tukey's Q tests conducted on $\delta^{15}N$ values among three sites, focusing on males with weapons and other adults without weapons.

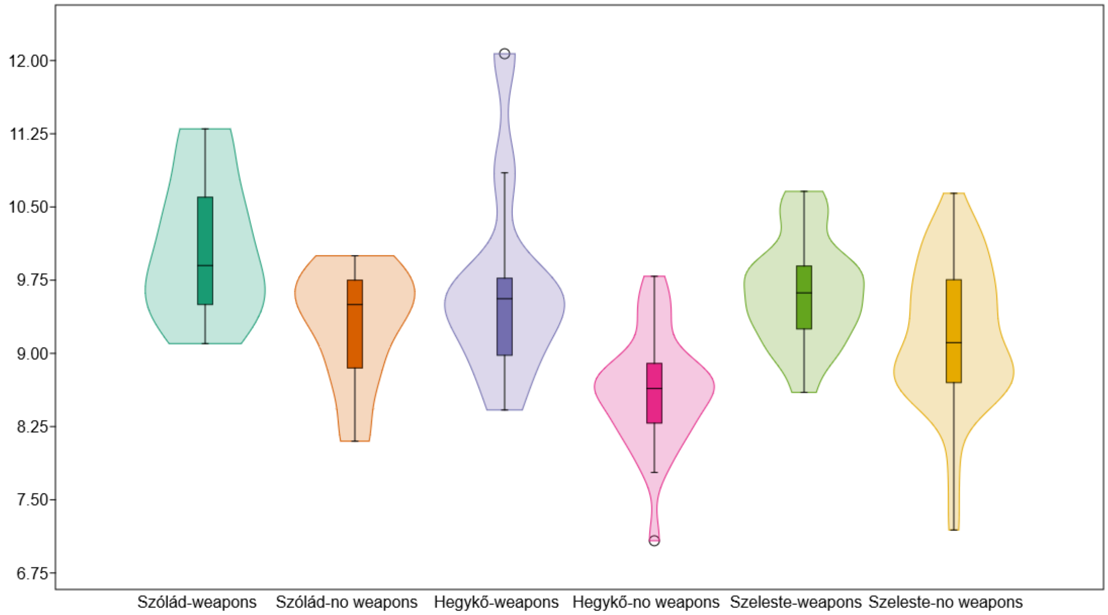

Supplementary Fig. 12. Comparison of $\delta^{15}N$ values across three sites, focusing on males with weapons and other adults without weapons.

**S3. Bootstrap analysis for fastNGSadmix results**

In order to assess the reliability of our fastNGSadmix admixture coefficients, we implemented a simple block bootstrapping procedure to estimate confidence intervals. A block bootstrap approach was chosen in order to incorporate the potentially inflated error arising from the correlation structure of neighboring SNPs being in linkage disequilibrium. Bootstrapping was performed over 10Mb non-overlapping blocks (as such the genome is divided into 300 discrete blocks across all autosomes), where 300 blocks are sampled randomly with replacement. We ran 50 bootstrap iterations for each individual and plotted the 90% confidence intervals (ranging from the 5th to the 95th quantile estimates) and the 50% confidence intervals (ranging from the 25th to the 75th quantile estimates) for each ancestry component across all 314 individuals. The mean SD for the eight ancestries are 0.2552% (SUBSAHARAN), 0.3274% (NAFRICA), 0.3840% (EASIA), 0.8138% (SASIA), 0.9026% (CASIA), 7.333% (MEDEU), 9.018% (SCAND) and 13.72% (NGBI). Non-Europeans ancestries had much smaller SDs. (Supplementary Fig. 14 and 15).

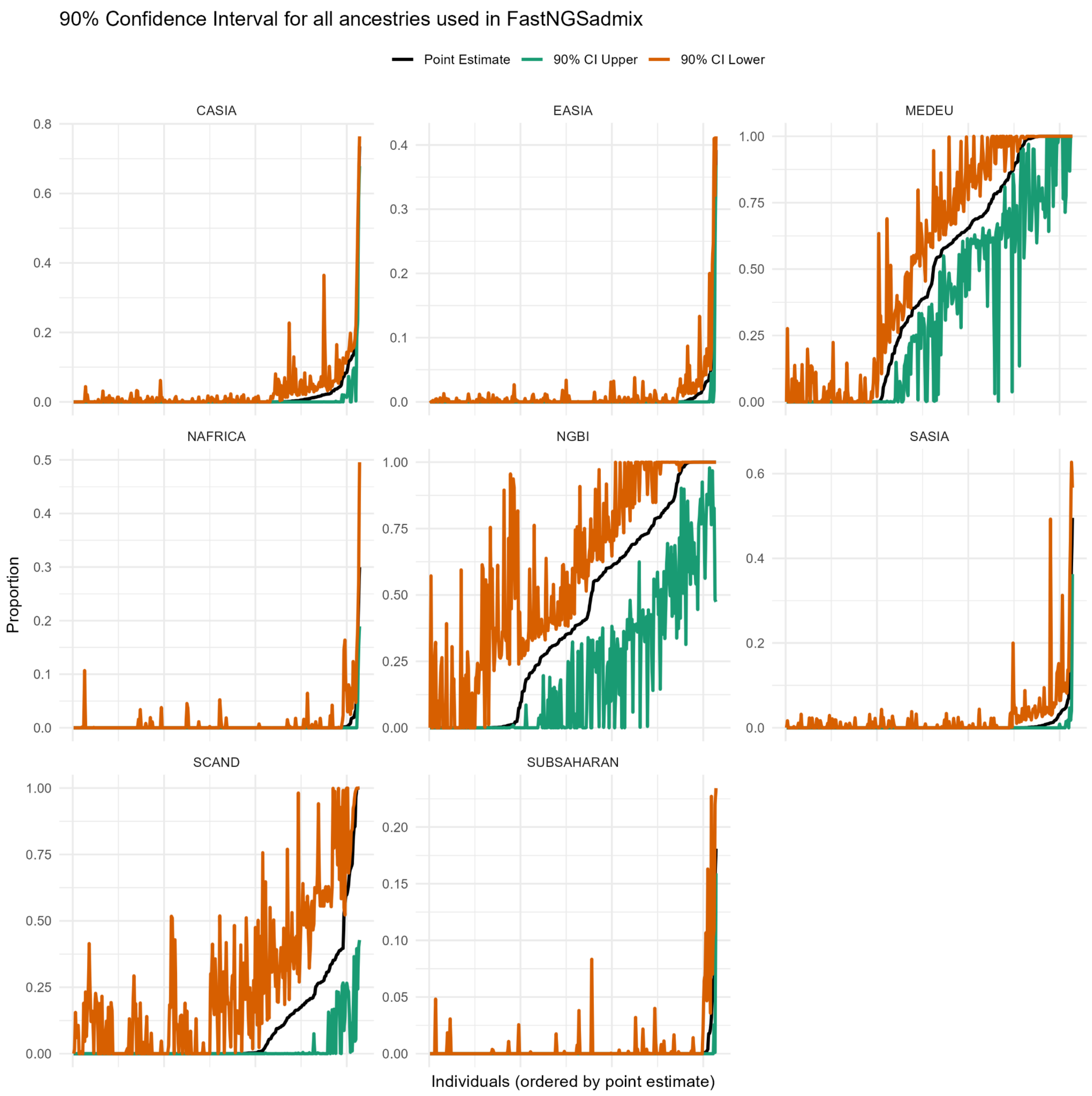

Supplementary Fig. 14. Bootstrap results for all ancestries, including point estimates and 90% confidence intervals

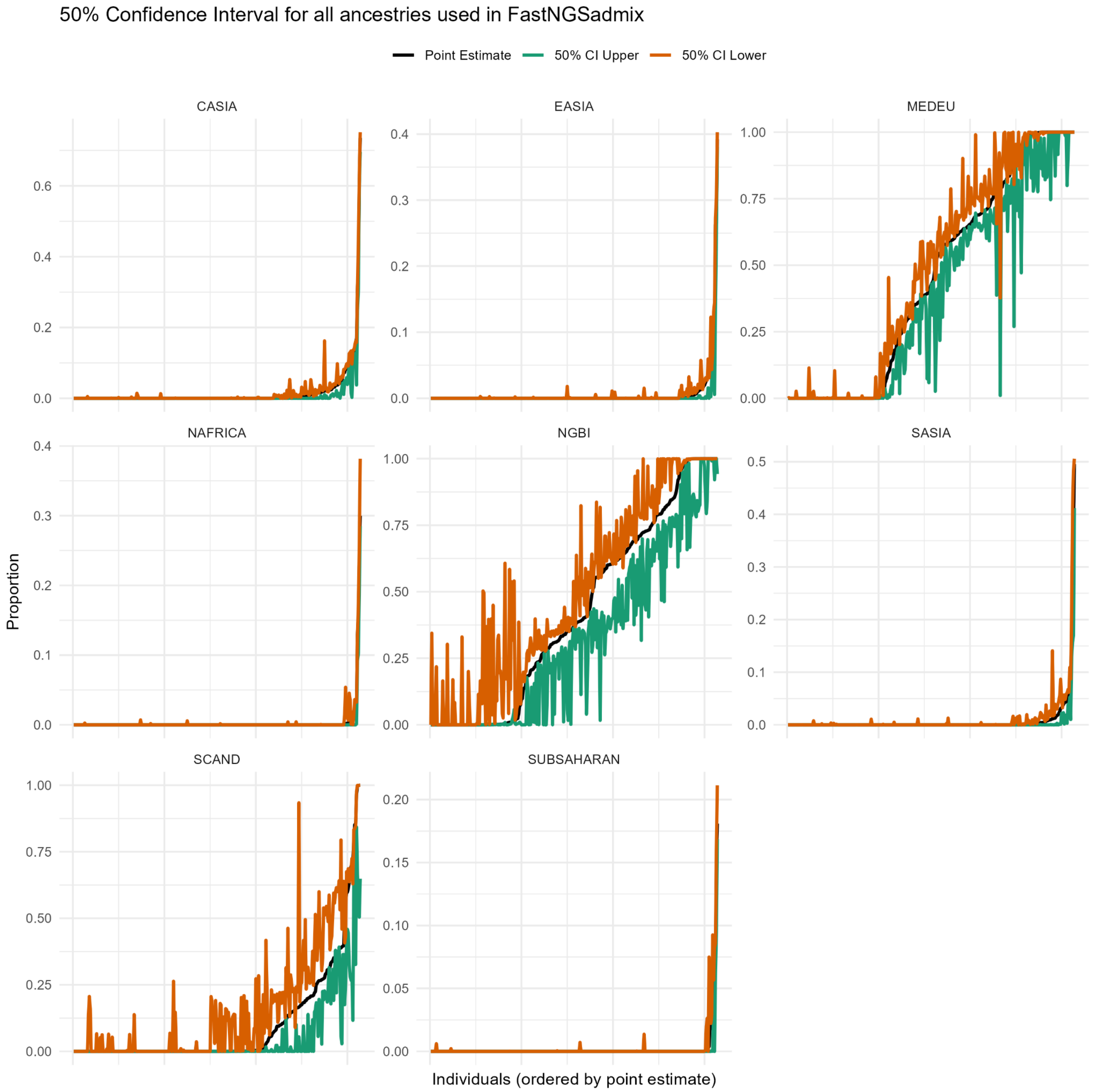

Supplementary Fig. 15. Bootstrap results for all ancestries, including point estimates and 50% confidence intervals

### S4. Comparing fastNGSadmix results with qpAdm results

To evaluate whether fastNGSadmix and qpAdm produce comparable results, we assessed the correlation between ancestry proportions estimated by the two methods (Supplementary Fig. 16). We first removed all unmodeled individuals identified by qpAdm, as well as individuals with coverage lower than 0.01×. Each ancestry component was then compared separately.

Using Pearson's correlation and linear regression in R, we found that for the non-European components, SASIA and SUBSAHARAN did not exhibit significant correlations between the two methods. This is likely due to the very small number of individuals with detectable levels of these ancestries, as well as the extremely low estimated proportions—conditions under which correlation analysis is not meaningful. In contrast, CASIA, EASIA, and NAFRICA all showed significant correlations, suggesting greater consistency between the two methods. However, the regression slopes (covariates) deviate from the expected value of ~1, likely due to the small number of individuals carrying these non-European ancestries.

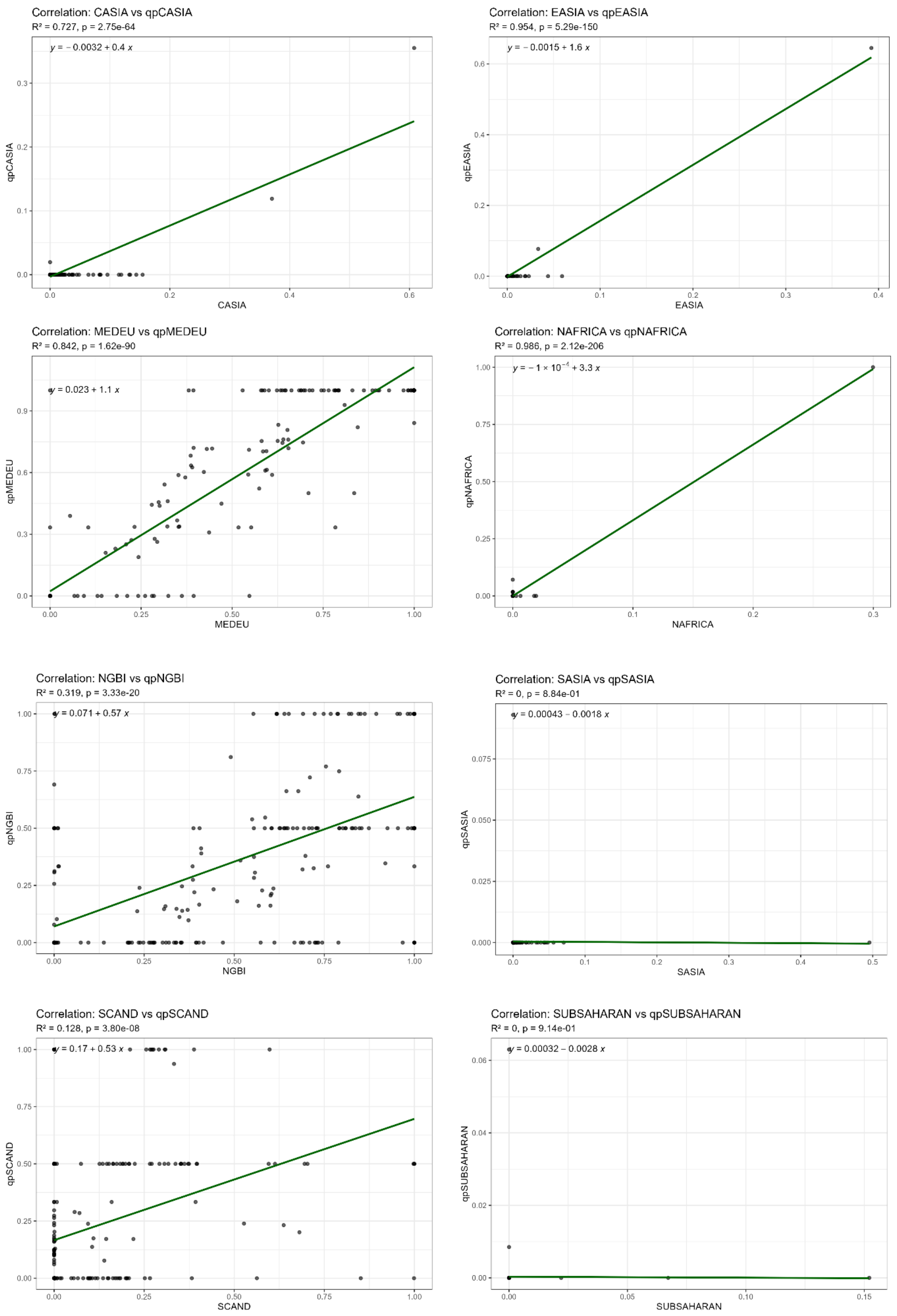

Supplementary Fig. 16. Correlation and linear regression analyses between fastNGSadmix and qpAdm results (starting with 'qp') for eight penecontemporaneous reference populations. Analyses included all individuals with data from both methods and a coverage greater than 0.01x.

For the European components, MEDEU showed a strong correlation between fastNGSadmix and qpAdm, with a regression slope of 1.1, indicating a high degree of similarity. NGBI and SCAND also showed significant correlations, but with regression slopes closer to 0.5. This may reflect the difficulty in distinguishing between these two ancestries. When NGBI and SCAND were combined into a single component, the regression slope increased to 1.1 (Supplementary Fig. 17), suggesting a more consistent and reliable estimation when these ancestries are not separated

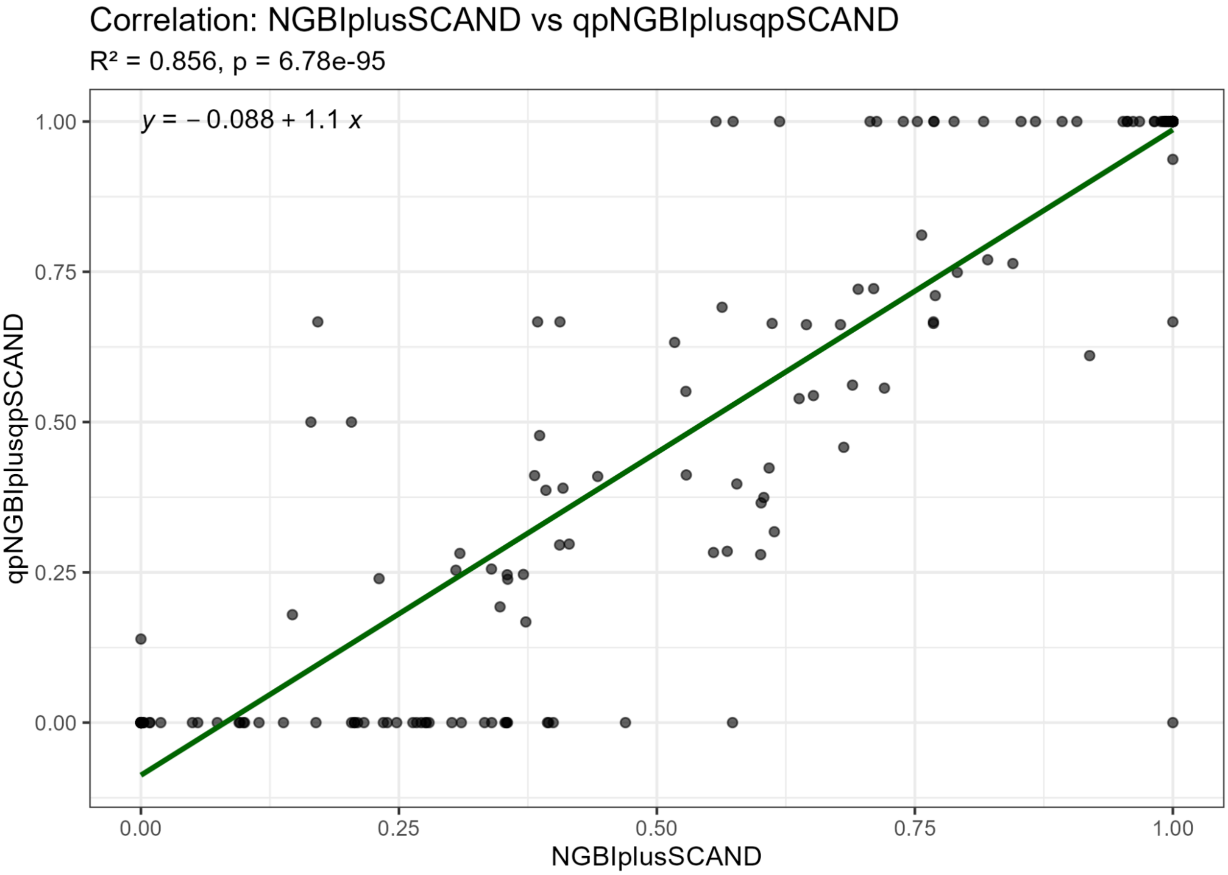

Supplementary Fig. 17. Correlation and linear regression analyses between fastNGSadmix and qpAdm results (starting with 'qp') for northern components (NGBI plus SCAND).

**S5. IBD analysis**

We first examined IBD connections between post-Roman individuals from the Little Hungarian Plain (LHP) and those from other populations. In total, we identified 244 pairs of IBD segments longer than 12 cM shared between post-Roman LHP individuals and individuals from other regions. These involved in total 118 individuals outside of the LHP (Supplementary Table S8), including 4 individuals from Croatia (n = 5 connections), 4 from Denmark (n = 6), 4 from Estonia (con = 6), 16 from Germany (n = 46), 20 from Hungary (n = 22), 6 from Ireland (n = 17), 9 from

Italy (n = 16), 1 from Kazakhstan (n = 1), 7 from the Netherlands (n = 54), 2 from Norway (n = 2), 6 from Sweden (n = 22), 1 from Turkey (n = 2), and 27 from the UK (n = 46).

| Ref_pop | Size | nb_conn_6th_inds | nb_conn_6th_pairs | total_pairwise_comps | prop_con_6th |
|---|---|---|---|---|---|
| Albania | 1 | | | 130 | 0.00% |
| Armenia | 4 | | | 520 | 0.00% |
| Austria | 8 | | | 1040 | 0.00% |
| Botswana | 1 | | | 130 | 0.00% |
| Bulgaria | 1 | | | 130 | 0.00% |
| Croatia | 48 | 4 | 5 | 6240 | 0.08% |
| Czech Republic | 1 | | | 130 | 0.00% |
| Denmark | 14 | 4 | 5 | 1820 | 0.27% |
| Estonia | 31 | 4 | 6 | 4030 | 0.15% |
| France | 5 | | | 650 | 0.00% |
| Germany | 46 | 16 | 46 | 5980 | 0.77% |
| Greece | 1 | | | 130 | 0.00% |
| Greenland | 1 | | | 130 | 0.00% |
| Hungary | 299 | 20 | 22 | 38870 | 0.06% |
| Iceland | 3 | | | 390 | 0.00% |
| India | 1 | | | 130 | 0.00% |
| Ireland | 12 | 6 | 17 | 1560 | 1.09% |
| Italy | 46 | 9 | 16 | 5980 | 0.27% |
| Kazakhstan | 20 | 1 | 1 | 2600 | 0.04% |
| Kenya | 1 | | | 130 | 0.00% |
| Kyrgyzstan | 13 | | | 1690 | 0.00% |

| | | | | | |
|---|---|---|---|---|---|
| Lebanon | 1 | | | 130 | 0.00% |
| Mongolia | 17 | | | 2210 | 0.00% |
| Montenegro | 1 | | | 130 | 0.00% |
| Netherlands | 16 | 7 | 54 | 2080 | 2.60% |
| Norway | 13 | 2 | 2 | 1690 | 0.12% |
| Poland | 3 | | | 390 | 0.00% |
| Portugal | 3 | | | 390 | 0.00% |
| Romania | 1 | | | 130 | 0.00% |
| Russia | 16 | | | 2080 | 0.00% |
| Serbia | 13 | | | 1690 | 0.00% |
| Slovakia | 2 | | | 260 | 0.00% |
| Slovenia | 1 | | | 130 | 0.00% |
| Spain | 6 | | | 780 | 0.00% |
| Sweden | 17 | 6 | 22 | 2210 | 1.00% |
| Taiwan | 15 | | | 1950 | 0.00% |
| Turkey | 25 | 1 | 2 | 3250 | 0.06% |
| UK | 154 | 27 | 46 | 20020 | 0.23% |
| ALL | 861 | 107 | 244 | 111930 | 0.22% |

Supplementary Table S8. Summary of IBD analysis between LHP 6th CE individuals and other populations

However, when focusing on IBD segments longer than 20 cM, only 23 pairs of relatedness remained, involving 19 individuals from outside the LHP. Notably, two individuals from Hegykő (Hegykő_47 and Hegykő_49, a parent-offspring pair) shared an approximately 20 cM segment with an individual from West Heslerton, UK (I11587). Another individual from Hegykő (Hegykő_36) shared ~20 cM with an individual from Dover Buckland, UK (BUK009), while Hegykő_35 shared ~25 cM with an individual from Lower Saxony, Germany (DRU016). Hegykő_11 also shared ~20 cM with an individual from Tian Shan, Kazakhstan.

In Szeleste, two individuals (Szeleste_498 and Szeleste_370-2) shared ~40 cM with the same individual from Haven, Germany (HVN004). Additionally, we found five connections between Szeleste individuals and those from Dover Buckland, UK: Szeleste_468 with BUK041, Szeleste_370-1 with BUK049, Szeleste_388 with BUK049, Szeleste_486 with BUK038, and Szeleste_437-1 with BUK053. Notably, Szeleste_370-1 and Szeleste_388 form another parent-offspring pair. Two Szeleste individuals also showed IBD sharing with individuals from Croatia: Szeleste_598-B with I26859 and Szeleste_534 with I26749. Szeleste_498 also shared segments with three individuals from Collegno, Italy (COL_069, COL_017, and COL_143), all from the same pedigree. Szeleste_437-2 shared ~30 cM with COL_017. Furthermore, Szeleste_380 had IBD connections with two individuals from Rákóczifalva, Hungary (RKF207 and RKF237) and one from Szólád, Hungary (Sz_28).

Additionally, Gyirmót_26 shared IBD segments with two individuals from Szólád, Hungary (AV1 and AV2).

Despite these findings, the longest IBD segment identified between LHP individuals and those from other populations was only 44 cM, and the highest sum of IBD sharing is 121cM, which only suggests ~6th degree relatedness.

We next examined IBD connections between Roman-period individuals from the LHP and other populations (Supplementary Table S9). In total, we identified 220 IBD-sharing pairs. Notably, two individuals—Gy-Szt_29 and Gy-Szt_32—accounted for a disproportionately large number of these connections. Gy-Szt_29 shared IBD segments with 49 individuals from diverse regions including Germany, the Netherlands, Kazakhstan, Ireland, Romania, Hungary, Denmark, Sweden, and Mongolia. Gy-Szt_32 exhibited shared IBD with 125 individuals spanning a remarkably wide geographic range, including Germany, Kyrgyzstan, Italy, Kazakhstan, the Netherlands, Hungary, Turkey, Croatia, India, Taiwan, Spain, Russia, Ireland, Kenya, Romania, Mongolia, Denmark, Sweden, Norway, and even Botswana. Several of these shared segments exceeded 55 cM in length. Despite passing quality filters, the extraordinary number and extent of connections suggest these two individuals may be problematic, and we treated them with caution in subsequent analyses.

| Ref | Size | nb_conn_3-5th_inds | nb_conn_3-5th_pairs | total_pairwise_comps | prop_con_3-5th |
|---|---|---|---|---|---|
| Albania | 1 | | | 31 | 0.00% |
| Armenia | 4 | | | 124 | 0.00% |
| Austria | 8 | | | 248 | 0.00% |
| Botswana | 1 | | | 31 | 0.00% |
| Bulgaria | 1 | | | 31 | 0.00% |

| | | | | | |
|---|---|---|---|---|---|
| Croatia | 48 | 1 | 1 | 1488 | 0.07% |
| Czech Republic | 1 | | | 31 | 0.00% |
| Denmark | 14 | 1 | 1 | 434 | 0.23% |
| Estonia | 31 | | | 961 | 0.00% |
| France | 5 | | | 155 | 0.00% |
| Germany | 46 | 2 | 7 | 1426 | 0.49% |
| Greece | 1 | | | 31 | 0.00% |
| Greenland | 1 | | | 31 | 0.00% |
| Hungary | 299 | 7 | 7 | 9269 | 0.08% |
| Iceland | 3 | | | 93 | 0.00% |
| India | 1 | | | 31 | 0.00% |
| Ireland | 12 | | | 372 | 0.00% |
| Italy | 46 | 1 | 1 | 1426 | 0.07% |
| Kazakhstan | 20 | | | 620 | 0.00% |
| Kenya | 1 | | | 31 | 0.00% |
| Kyrgyzstan | 13 | | | 403 | 0.00% |
| Lebanon | 1 | | | 31 | 0.00% |
| Mongolia | 17 | | | 527 | 0.00% |
| Montenegro | 1 | 1 | 1 | 31 | 3.23% |
| Netherlands | 16 | 5 | 12 | 496 | 2.42% |
| Norway | 13 | | | 403 | 0.00% |
| Poland | 3 | | | 93 | 0.00% |
| Portugal | 3 | | | 93 | 0.00% |
| Romania | 1 | | | 31 | 0.00% |
| Russia | 16 | | | 496 | 0.00% |

| | | | | | |
|---|---|---|---|---|---|
| Serbia | 13 | 3 | 3 | 403 | 0.74% |
| Slovakia | 2 | 1 | 1 | 62 | 1.61% |
| Slovenia | 1 | | | 31 | 0.00% |
| Spain | 6 | | | 186 | 0.00% |
| Sweden | 17 | 3 | 7 | 527 | 1.33% |
| Taiwan | 15 | | | 465 | 0.00% |
| Turkey | 25 | | | 775 | 0.00% |
| UK | 154 | | | 4774 | 0.00% |
| ALL | 861 | 25 | 41 | 26691 | 0.15% |

Supplementary Table S9. Summary of IBD analysis between LHP 3-5th CE individuals and other populations

After excluding Gy-Szt_29 and Gy-Szt_32, we identified 44 IBD-sharing pairs involving 27 individuals outside the LHP. These include 1 individual from Croatia (n =1), 1 individual from Denmark (n = 1), 2 from Germany (n = 7), 7 from Hungary (n = 7), 1 from Italy (n = 1), 1 from Montenegro (n = 1), 5 from Netherlands (n = 12), 3 from Serbia (n = 3), 1 form Slovakia (n = 1), and 3 from Sweden (n = 7). None of the connections has IBD segments longer than 20cM.

**S6. Comparison between fastNGSadmix results and Local Ancestry Inference results using flare**

To assess the reliability of Local Ancestry Inference using Flare, we compared its global ancestry estimates with those from fastNGSadmix using Pearson's correlation and linear regression in R. The analysis (Supplementary Fig. 18) included all LHP individuals with coverage ≥1x, consistent with the threshold used in the IBD analysis. The results indicate that both Northern (nEUR) and Southern (sEUR) European ancestries are highly correlated between the two methods ($p < 0.0001$ for both). However, Flare tends to estimate ancestry proportions more conservatively—rarely reaching exact values of 0 or 1—whereas fastNGSadmix often assigns these boundary values. We also observed a significant correlation for non-European ancestries; however, the regression fit was weaker compared to that of Northern and Southern ancestries, likely due to their low representation within the population.

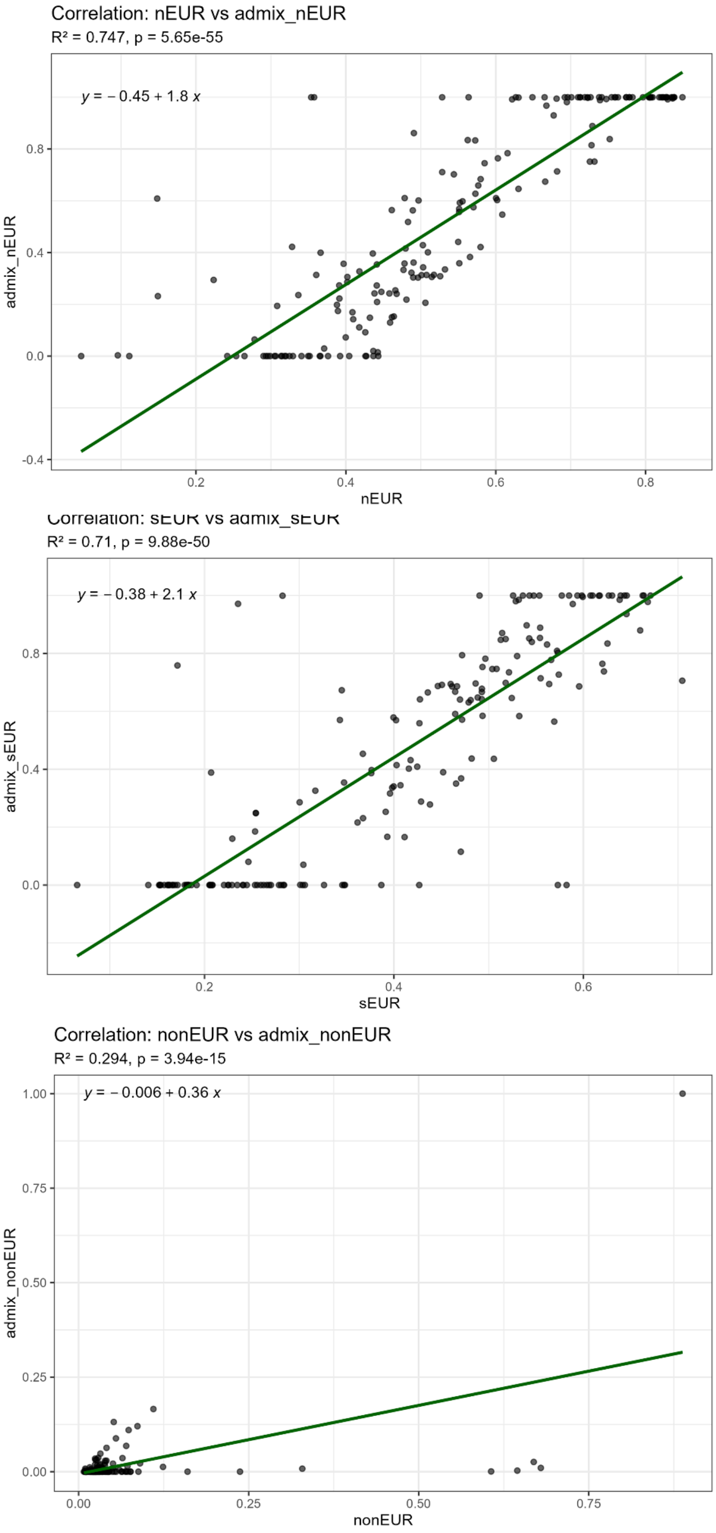

Supplementary Fig. 18. Correlation and linear regression analyses between fastNGSadmix and Local Ancestry Inference results. nEUR - Northern European; sEUR - Southern European; nonEUR - non-European

### S7. Compare the results of lcMLkin and KIN

We compared the initial degree of relatedness inferred by KIN with the results generated by lcMLkin (Dataset S9, column "lcMLkin_compare"). Among the 213 related pairs (up to third degree) identified by KIN, 95 had fewer than 10,000 shared SNPs and were therefore excluded from the lcMLkin analysis. Of the remaining 117 pairs, 105 (90%) showed consistent results between KIN and lcMLkin. An additional six pairs matched the second-best guess from KIN, and two were consistent with KIN's second-best guess within the same degree category. For the five remaining pairs, four were classified as third-degree relatives by KIN but as second-degree relatives according to pedigree-informed lcMLkin results. The last pair was identified as third-degree by KIN but as unrelated by lcMLkin.

### References

1. Haas, J. & Jámbor, Á. Quaternary Evolution. in *Geology of Hungary* 201–213 (Springer Berlin Heidelberg, Berlin, Heidelberg, 2012).
2. *Landscapes and Landforms of Hungary*. (Springer International Publishing, Cham, Switzerland, 2015).
3. Csillag, G. & Sebe, K. Long-Term Geomorphological Evolution. in *World Geomorphological Landscapes* 29–38 (Springer International Publishing, Cham, 2015).
4. Gábris, G. & Nádor, A. Long-term fluvial archives in Hungary: response of the Danube and Tisza rivers to tectonic movements and climatic changes during the Quaternary: a review and new synthesis. *Quat. Sci. Rev.* **26**, 2758–2782 (2007).
5. *Magyarország kistájainak katasztere*. (HUN-REN CSFK Geographical Institute | Library |

Institutional publications, Budapest, 2010).

6. Bentley, R. A., Price, T. D. & Stephan, E. Determining the ‘local’ 87Sr/86Sr range for archaeological skeletons: a case study from Neolithic Europe. *J. Archaeol. Sci.* **31**, 365–375 (2004).

7. Price, T. D., Burton, J. H. & Bentley, R. A. The characterization of biologically available strontium isotope ratios for the study of prehistoric migration. *Archaeometry* **44**, 117–135 (2002).

8. Alexander Bentley, R. Strontium Isotopes from the Earth to the Archaeological Skeleton: A Review. *Journal of Archaeological Method and Theory* **13**, 135–187 (2006).

9. Makarewicz, C. A. & Sealy, J. Dietary reconstruction, mobility, and the analysis of ancient skeletal tissues: Expanding the prospects of stable isotope research in archaeology. *J. Archaeol. Sci.* **56**, 146–158 (2015).

10. Maurer, A.-F. *et al.* Bioavailable 87Sr/86Sr in different environmental samples--effects of anthropogenic contamination and implications for isoscapes in past migration studies. *Sci. Total Environ.* **433**, 216–229 (2012).

11. Depaermentier, M. L. C., Kempf, M., Bánffy, E. & Alt, K. W. Modelling a scale-based strontium isotope baseline for Hungary. *J. Archaeol. Sci.* **135**, 105489 (2021).

12. Knipper, C. Die Strontiumisotopenanalyse: Eine naturwissenschaftliche Methode zur Erfassung von Mobilität in der Ur- und Frühgeschichte. *jrgzm* **51**, 589–686 (2004).

13. Giblin, J. I. Strontium isotope analysis of Neolithic and Copper Age populations on the Great Hungarian Plain. *J. Archaeol. Sci.* **36**, 491–497 (2009).

14. Katzenberg, M. A. Stable isotope analysis: a tool for studying past diet, demography, and life history. *Biological anthropology of the human skeleton* **2**, 413–441 (2008).

15. Lee-Thorp, J. A. On isotopes and old bones. *Archaeometry* **50**, 925–950 (2008).

16. Ventresca Miller, A. R. & Makarewicz, C. A. Intensification in pastoralist cereal use coincides with the expansion of trans-regional networks in the Eurasian Steppe. *Sci. Rep.*

**9**, 8363 (2019).

17. Martin, L. *et al.* The place of millet in food globalization during Late Prehistory as evidenced by new bioarchaeological data from the Caucasus. *Sci. Rep.* **11**, 13124 (2021).
18. Hedges, R. E. M. & Reynard, L. M. Nitrogen isotopes and the trophic level of humans in archaeology. *J. Archaeol. Sci.* **34**, 1240–1251 (2007).
19. Ambrose, S. H. Effects of diet, climate and physiology on nitrogen isotope abundances in terrestrial foodwebs. *J. Archaeol. Sci.* **18**, 293–317 (1991).
20. van Klinken, G. J. Bone collagen quality indicators for palaeodietary and radiocarbon measurements. *J. Archaeol. Sci.* **26**, 687–695 (1999).
21. Alt, K. W. *et al.* Lombards on the move – an integrative study of the migration period cemetery at szólád, Hungary. *PLoS One* **9**, e110793 (2014).
